# An Improved Chromosome-scale Genome Assembly and Population Genetics resource for *Populus tremula*

**DOI:** 10.1101/805614

**Authors:** Kathryn M. Robinson, Bastian Schiffthaler, Hui Liu, Sara M. Westman, Martha Rendón-Anaya, Teitur Ahlgren Kalman, Vikash Kumar, Camilla Canovi, Carolina Bernhardsson, Nicolas Delhomme, Jerry Jenkins, Jing Wang, Niklas Mähler, Kerstin H. Richau, Victoria Stokes, Stuart A’Hara, Joan Cottrell, Kizi Coeck, Tim Diels, Klaas Vandepoele, Chanaka Mannapperuma, Eung-Jun Park, Stephane Plaisance, Stefan Jansson, Pär K. Ingvarsson, Nathaniel R. Street

## Abstract

Aspen (*Populus tremula* L.) is a widely distributed keystone species and a model system for forest tree genomics, with extensive resources developed for population genetics and genomics. Here we present an updated resource comprising a chromosome-scale assembly of *P. tremula* and population genetics and genomics data integrated into the PlantGenIE.org web resource. We demonstrate use of the diverse data types included to explore the genetic basis of natural variation in leaf size and shape as examples of traits with complex genetic architecture.

We present a chromosome-scale genome assembly generated using long-read sequencing, optical and high-density genetic maps containing 39,894 annotated genes with functional annotations for 73,765 transcripts from 37,184 gene loci. We conducted whole-genome resequencing of the Umeå Aspen (UmAsp) collection comprising 227 aspen individuals. We utilised the assembly, the UmAsp re-sequencing data and existing whole genome re-sequencing data from the Swedish Aspen (SwAsp) and Scottish Aspen (ScotAsp) collections to perform genome-wide association analyses (GWAS) using Single Nucleotide Polymorphisms (SNPs) for leaf physiognomy phenotypes. We conducted Assay of Transposase Accessible Chromatin sequencing (ATAC-Seq) and identified genomic regions of accessible chromatin and subset SNPs to these regions, which improved the GWAS detection rate. We identified candidate long non-coding RNAs in leaf samples and quantified their expression in an updated co-expression network (AspLeaf, available in PlantGenIE.org), which we further used to explore the functions of candidate genes identified from the GWAS.

We examined synteny to the reference *P. trichocarpa* assembly and identified *P. tremula*-specific regions. Analysis of whole-genome duplication indicated differential substitution rates for the two *Populus* species, indicating more rapid evolution in *P. tremula*. A GWAS of 26 leaf physiognomy traits and all SNPs in each of the three aspen collections found significant associations for only two traits in ScotAsp collection and one in UmAsp, whereas subsetting SNPs to those in open chromatin regions revealed associations for a further four traits among all three aspen collections. The significant SNPs were associated with genes annotated for developmental and growth functions, which represent candidates for further study. Of particular interest was a 177-kbp region of chromosome 9 harbouring SNPs associated with multiple leaf phenotypes in ScotAsp, with the set of SNPs in linkage disequilibrium explaining 24 to 30 % of the phenotypic variation in leaf indent depth variation.

We have incorporated the assembly, population genetics, genomics and leaf physiognomy GWAS data into the PlantGenIE.org web resource, including updating existing genomics data to the new genome version. This enables easy exploration and visualisation of the genomics data and exploration of GWAS results. We provide all raw and processed data used for the presented analyses to facilitate reuse in future studies.

## Introduction

The *Populus* genus encompasses around thirty broad-leaved, fast-growing tree species that occur naturally across most of the Northern hemisphere. *Populus* species are used extensively in short-rotation forestry and landscaping worldwide and are pioneer, keystone species. The black cottonwood, *P. trichocarpa,* was the first tree genome to be sequenced (Tuskan *et al*., 2006) after which the genomes of several other poplars, aspens and cottonwoods have been published (Yang *et al*., 2017; Lin *et al*., 2018; Ma *et al*., 2019; An *et al*., 2020; Wu *et al*., 2020; Zhang *et al*., 2020; Bai *et al*., 2021; Chen *et al*., 2023a; Bae *et al*., 2023, Zhou *et al*., 2023, Shi *et al*., 2024), firmly establishing *Populus* as a model system for forest tree research with a mature genomics resource (Jansson & Douglas, 2007). The aspens (section Populus) include *P. tremula* and *P. alba,* which have ranges spanning northern Eurasia*, P. tremuloides* and *P. grandidenta,* native to North America, and *P. adenopoda, P. qiongdaoensis* and *P. davidiana,* distributed in northern and eastern Asia (Slavov & Zhelev, 2010; Hou *et al*., 2018). They are recognised by their capacity for clonal regeneration, particularly after environmental perturbation such as fire or intense browsing (Myking *et al*., 2011). Other distinguishing features of aspens are their characteristic leaf tremble, and abundant variation in spring and autumn leaf colouration.

The availability of a reference genome can be transformative in enabling research of a species, opening possibilities for a range of functional genomics, population genetics and comparative genomics studies. We previously described a reference genome for *P. tremula* (Lin *et al*., 2018) produced using short-read, second generation sequencing technologies. While this genome assembly provided high quality and comprehensive representation of the gene space, it was highly fragmented and lacked long-range contiguity. Such fragmentation is a common limitation of using short read sequencing technologies to assemble highly heterozygous, repeat-rich or polyploid genomes (Jiao & Schneeberger, 2017). These limitations can be alleviated or overcome, depending on the scale of the challenge, by use of third generation sequencing technologies such as those commercialised by Pacific Bioscience or Oxford Nanopore Technologies, which produce vastly longer individual sequence reads (Jiao & Schneeberger, 2017). These long reads simplify assembly, with individual sequencing reads often being sufficiently long to span a repeat element or a heterozygous region, although such haplotype resolved assembly ability also introduces its own set of challenges (Amarasinghe *et al*., 2020; Michael & VanBuren, 2020). These technologies can be combined with newly developed or improved methods for scaffolding, such as Hi-C or optical mapping, to further improve long-range assembly contiguity (Ghurye & Pop, 2019; Ghurye *et al*., 2019; Pan *et al*., 2019). The vastly improved contiguity achieved also facilitates use of genetic maps to anchor and orient assembled scaffolds to produce chromosome scale assemblies. Improved contiguity is essential for synteny and other comparative genome-based analyses and highly contiguous and accurate assemblies provide a more reliable resource for performing gene family and orthology analyses and for designing guide sequencing to perform genome editing using approaches such as CRISPR-Cas.

We previously reported the evolutionary divergence of *P. trichocarpa* from the aspens, showing how natural variation has shaped genetic relationships among the European/Eurasian (*P. tremula*) and American (*P. tremuloides* and *P. grandidentata*) aspens (Wang *et al*., 2016a; Wang *et al*., 2016b; Lin *et al*., 2018; Apuli *et al*., 2020). An important resource for such work is the Swedish Aspen (SwAsp) collection, which exhibits considerable heritable variation in numerous phenotypes including phenology, leaf shape, specialised metabolite composition and ecological interactions (Luquez *et al*., 2008; Robinson *et al*., 2012; Bernhardsson *et al*., 2013; Wang *et al*., 2018; Mähler *et al*., 2020) in addition to gene expression (Mähler *et al*. 2017). While we previously reported GWAS resulting in the discovery of a major locus for an adaptive phenological trait (Wang *et al*., 2018), most of the phenotypes considered to date have not yielded significant SNP-phenotype associations (Grimberg *et al*. 2018; Mähler *et al*., 2020), likely indicative of complex and highly polygenic genetic architecture (e.g., Mähler *et al*., 2020). However, other factors such as the fragmented nature of the v1.1 genome assembly (Lin *et al*., 2018), rare or non-SNP variants and a relatively small population size are also likely to contribute to the limited ability to detect significant associations (Street & Ingvarsson, 2011).

Here we present a chromosome-scale genome assembly for *P. tremula*, which we refer to as *P. tremula* v2.2, generated using long-read sequences and optical and genetic maps. We demonstrate utility of this improved genome assembly by performing SNP calling and GWAS for selected leaf physiognomy traits with complex genetic architecture in three collections of wild aspen trees grown in common gardens, including the Umeå Aspen (UmAsp) collection for which we here present whole-genome resequencing data. To facilitate community access and utilisation of the various datasets available for *P. tremula* we have integrated them into the PlantGenIE.org web resource (Sundell *et al*., 2015) in addition to making all raw and processed data available at public repositories. We provide examples of how these genetics and genomics datasets can be used to explore or develop hypotheses and how the tools available at PlantGenIE.org can be used to gain additional biological insight for identified candidate genes.

## Materials and Methods

We extracted DNA from the individual used to generate the v1.1 assembly presented in Lin *et al*. (2018). For genome assembly and correction, we generated two libraries: “PacBio data”: 28,874,072,954 bases (filtered subreads, ∼6∽0x coverage), Pacific Biosciences on the RSII platform (sequencing performed by Science for Life Laboratory, Uppsala, Sweden); and “Illumina data”: 108,353,739,802 bases (2∽26x coverage), Illumina HiSeq2500. We also utilised five existing RNA-Seq datasets to support gene annotation. We purified nuclei and produced an ATAC-library. We used five RNA-Seq datasets from *P. tremula* as supporting evidence for gene annotation. We called ATAC-Seq peaks using MACS2 v2.2.7.1 (Zhang *et al*., 2008).

To flag sequences originating from the chloroplast, we matched all unplaced scaffolds to published chloroplast sequences (Kersten *et al*., 2016), using blast+.

We created a custom *de novo* repeat library using RepeatModeler v1.0.11 and subsequently masked the genome using RepeatMasker4.0.8. (http://www.repeatmasker.org). We used Trinity assemblies from all RNA-Seq datasets in conjunction with all annotated transcripts from the v1 assembly as evidence for gene annotation. We provided proteins from the v1 *P. tremula* assembly and the v3.0 assembly of *P. trichocarpa* (Tuskan *et al*., 2006) as protein evidence. We performed the lift-over alignments of v1.1 scaffolds to the v2.2 assembly using minimap 2 (v2.2.2, Li, 2018). We aligned the PacBio data to both the v1.1 and v2.2 assemblies using pbmm2 (v1.1.0), which uses minimap2 internally. We then performed variant detection using pbsv (v2.2.2, https://github.com/PacificBiosciences/pbsv). We aligned the transcripts and protein-coding sequences retrieved from MAKER to the NCBI nt (Wheeler *et al*., 2007) and UniRef90 (The UniProt Consortium, 2019) databases. For transcripts, we used Blast+ version 2.6.0+ (Altschul *et al*., 1990) with the non-default parameters: -e-value 1e^-5^. For proteins, we used Diamond version 0.9.26 (Buchfink *et al*., 2014). We identified and extracted the sequences aligning solely to the NCBI nt database to complement the UniRef90 alignments using an ad-hoc script (available upon request). We then imported the resulting alignment files into Blast2GO (Götz *et al*., 2008) version 5.2. Finally, we used Blast2GO to generate Gene Ontology (both GO and GO-Slim), Pfam (El-Gebali *et al*., 2018) and KEGG (Kanehisa & Goto, 2000) annotations.

To calculate summary statistics of the assembly, we used QUAST v5.0.2 (Gurevich *et al*., 2013), aligning a 20X coverage subset (generated by truncating the library to a total count of 8 * 10e^9^ nucleotides) of the aspen V1 2×150 PE library data (ENA: PRJEB23581) to calculate mapping percentages. We ran BUSCO v3.0.2 for both the genomic and transcript sequences.

To identify homologous chromosomes between *P. tremula* and *P. trichocarpa* genomes, we used minimap2 (Li, 2018). We performed an all-versus-all BLASTP using protein sequences of *P. tremula* and *P. trichocarpa* to identify homologous gene pairs between the two species. We used MCscanX (Wang *et al*., 2012) to identify syntenic gene blocks. We aligned the protein sequences for duplicate gene pairs in syntenic blocks using MAFFT (Katoh & Standley, 2013). We used the PAML package (Yang, 2007) to estimate the *Ks* and *Ka*/*Ks* for each gene pair.

We used 32 plant genomes, (Supplementary table S1, Appendix S1) to perform gene family analysis. We used Orthofinder v2.2.7 (Emms & Kelly, 2015) to cluster the genes into gene families. Gene family trees were constructed using the PLAZA pipeline (Van Bel *et al*., 2018), for multiple sequence alignment and tree inference. We used muscle as the multiple sequence alignment method and fasttree as the tree construction method. The species tree was inferred using STAG (https://github.com/davidemms/STAG). To estimate the divergence time, we first calibrated the species tree based on the divergence dates from Timetree (http://www.timetree.org/) and inferred the divergence time on each clade using r8s (Sanderson, 2003). We inferred the expansion and contraction of the gene families using CAFÉ (De Bie *et al*., 2006) and the species tree.

We reprocessed the RNA-Seq data in developing terminal leaves of aspen from Mähler *et al*. (2020), using the v2.2 genome assembly with salmon/1.0.0 (Patro *et al*., 2017). We mapped expression quantitative trait loci (eQTL) using two different methods: (1) following the method and settings used in Mähler *et al*. (2020) using Matrix eQTL (Shabalin, 2012), and (2) using fastJT (Lin *et al*., 2019) which has no underlying assumption of the phenotypic distribution. For both methods, the eQTL were considered significant at FDR < 0.05.

We implemented a pipeline to identify putative long intergenic non-coding RNAs (lincRNAs) on the pre-processed RNA-Seq data. We first *in silico-*normalised the reads to reduce data redundancy and then reconstructed the transcriptome using a *de novo* assembler, Trinity (v. 2.8.3; Grabherr *et al*., 2011; Haas *et al*., 2013), on which other programs were run. We retained only transcripts being expressed in the dataset, that were identified as having no coding potential by PLEK (predictor of long non-coding RNAs and messenger RNAs based on an improved k-mer scheme; v. 1.2; settings -minlength 200; A. Li *et al*., 2014), CNCI (Coding-Non-Coding Index; v. 2; Sun *et al*., 2013), and CPC2 (Coding Potential Calculator version 2; v. 2.0 beta; settings -r TRUE; Kang *et al*., 2017) and being longer than 200 nt. We kept only transcripts identified as having no coding potential by TransDecoder (version 2.8.3; https://github.com/TransDecoder/TransDecoder/wiki; Haas *et al*., 2013), and at a distance > 1000 nt from any annotated gene using BEDTools closest (v. 2.30.0; https://bedtools.readthedocs.io/en/latest/content/tools/closest.html; Quinlan & Hall, 2010). The DESeq2 package (v. 1.42.0; Love *et al*., 2014) was used for differential expression analysis with the formula based on consecutive leaf developmental series, both for genes and lincRNAs. Then, lincRNAs and gene expression data were transformed to homoscedastic, asymptotical log_2_ counts using the variance stabilising transformation as implemented in DESeq2 (setting blind=FALSE). We retained all the genes and lincRNAs being different from na in the dds results and used their values from the ‘aware’ variance stabilising transformation as an input for the co-expression network. Thereafter, ten network inference methods were run using the Seidr toolkit (Schiffthaler *et al*., 2023). The networks were aggregated using the inverse rank product method (Zhong *et al*., 2014) and edges were filtered according to the noise-corrected backbone (Coscia & Neffke, 2017). We selected backbone8 to be used for further analyses.

We measured phenotypic traits in three aspen (*P. tremula*) collections, two of which originate from Sweden and one from Scotland. The Swedish Aspen (SwAsp) collection of 113 individuals, collected across ten degrees of latitude and longitude (Luquez *et al*., 2008), is replicated in two common gardens in Sweden, one in the north (Sävar, ∼64 °N) and one in the south (Ekebo, ∼56 °N). The Umeå Aspen (UmAsp) collection comprises 242 individuals originating from the Umeå municipality in northern Sweden (Fracheboud *et al*., 2009; Robinson *et al*., 2014) growing in a common garden at Sävar (∼64 °N). The Scottish Aspen (ScotAsp) collection of 138 trees originating from across Scotland, was cloned and grown in plots of five trees per clone, in a common garden at Forest Research, Roslin, UK (∼56 °N) Harrison (2009).

Details of the SwAsp and ScotAsp DNA sequencing and SNP calling have been described previously (Rendón-Anaya *et al*., 2021, Supplementary table S2). Samples sequenced from the previously generated data set comprising 94 genotypes from the SwAsp collection (Wang *et al*., 2018) have been complemented with a further five genotypes re-sequenced for *P. tremula* v2.2. We called SNPs and generated VCF files independently for SwAsp, UmAsp and ScotAsp (with 99, 227 and 105 unrelated individuals, respectively), containing biallelic, high quality sites along the 19 chromosomes. We also created a VCF for each collection containing a subset of all SNPs by intersecting with open chromatin regions identified by ATAC-Seq. The intersection was performed using bcftools (Danecek *et al*., 2021).

We measured leaf physiognomy (shape and size) parameters in six leaves of three clonal replicate trees in the UmAsp common garden, and fifteen leaves sampled across five clonal replicates per genotype in the ScotAsp Roslin common garden. We sampled mature, undamaged leaves, scanned them using a flatbed scanner, and measured using LAMINA software (Bylesjö *et al*., 2008) following methods described in Mähler *et al*. (2020). We present leaves sampled from the SwAsp common gardens as reported in Mähler *et al*. (2020) for leaf area, leaf circularity and leaf indent depth, and a further 23 leaf shape and size metrics for the analysis with SNPs called from *P. tremula* v2.2. We estimated Best Linear Unbiased Predictor (BLUP) values for each of the 26 phenotypes used in the GWAS (i.e. 26 phenotypes from each collection) using a custom pipeline. We used these BLUP estimates as phenotypic values to carry out GWAS in 99 SwAsp, 227 UmAsp, or 105 ScotAsp individuals for which SNP data were available and that remained after removing some samples due to high relatedness (IDB, identity by descent > 0.4). For the GWAS, we filtered SNPs with a minor allele frequency above 5% and Hardy-Weinberg equilibrium *P-*value threshold of 1e^-6^ using PLINK version 1.9. (Purcell *et al*., 2007). We investigated genome-wide associations using linear mixed models in GEMMA v0.98.1 (Zhou & Stephens, 2012), on (1) all SNPs, and (2) on SNPs subset to open chromatin regions. We used a 5% false discovery rate (*q*-value) to define associations as significant, calculated in the ‘qvalue’ package in R (Storey *et al*., 2021). We annotated the SNPs using ANNOVAR v2019Oct24 to produce GWAS summary tables, adding *A. thaliana* homologues of the *P. tremula* v2.2 gene models from PlantGenIE.org. We estimated the proportion of phenotypic variation explained (PVE) by an individual SNP using the equation stated in Wang *et al*. (2018). We calculated marker-based heritability (*h*^2^, Kruijer *et al*., 2015) using ‘marker_h2’ function, in the ‘heritability’ package version 1.3 (Kruijer, 2019) in R. We visualised the gene ontology enrichments using the PlantGenIE.org tool and additional visualisation using default R scripts exported from REVIGO (Supek *et al*., 2011)

For population genetic analysis, we discarded SNPs in SwAsp, UmAsp, and ScotAsp failing the Hardy–Weinberg equilibrium test (*P*-value < 1e^-6^) and/or with missing rate > 5%. We used SNPs with minor allele frequency >10% and missing rate <20% for linkage disequilibrium (LD) analysis. We calculated squared correlation coefficients (*r*^2^) between all pairs of SNPs that were within 50 Kbp using PopLDdecay v3.41 (Zhang *et al*., 2019). To analyse the population structure based on the PCA, we pruned SNPs by removing one SNP from each pair of SNPs with a between SNP correlation coefficient (*r*^2^) > 0.2 in windows of 50 SNPs with a step of 5 SNPs using PLINK v1.90b6.16 (Purcell *et al*., 2007). We then used the smartpca program in EIGENSOFT v6.1.4 (Patterson *et al*., 2006) to perform a principal components analysis (PCA) on the reduced set of genome-wide independent SNPs.

We calculated the composite likelihood ratio (CLR) statistic in 10 Kbp non-overlapping windows using SweepFinder2 and iHH12 (Integrated Haplotype Homozygosity Pooled) using selscan v1.3. To identify regions under positive selection, we used sliding windows containing at least 10 SNPs as input to a range of inference methods. We considered windows with the lowest 5% Tajima’s D values or highest 5% values for other as those windows displaying evidence of signals of positive selection. We assumed genes or SNPs within these selected regions to be under selection. We ran Betascan (Siewert *et al*., 2017) (-fold -m 0.1) to detect possible signals of balancing selection in the ScotAsp, UmAsp and SwAsp collections.

## Resource overview

The resource comprises the new *P. tremula* v2.2 genome assembly and gene annotation, genomic data sets (for example, *P. tremula*-specific regions, open chromatin regions, and lincRNAs), population genetics resources (SNPs and regions/SNPs under selection), both raw and processed leaf physiognomy phenotype data, and results of the leaf physiognomy GWAS analyses. The genomic resources are available at PlantGenIE.org where the genome assembly, gene annotation, gene expression data and associated co-expression networks are available through interactive tools and as flat files on the File Transfer Protocol (FTP) site (https://plantgenie.org/FTP). The assembly, open chromatin regions, lincRNAs, gene models, SNP variants and sites under selection have been made available as tracks in the JBrowse genome browser tool. The SwAsp leaf bud gene expression data and expression quantitative trait loci (eQTL) analysis presented in Mähler *et al*. (2017) have been updated to *P. tremula* v2.2. The AspLeaf dataset (Mähler *et al*. 2020) was used to identify lincRNAs and an updated co-expression network including these has been included in the exNet tool. Expression data sets can also be viewed in the exImage, exPlot and exHeatmap tools. The Potra v2.2 SNPs in SwAsp, UmAsp and ScotAsp are available in VCF format at the European Variation Archive (EVA, https://www.ebi.ac.uk/eva/). The phenotype data can be accessed at the SciLifeLab Data Repository (https://figshare.scilifelab.se/): this includes original raw leaf images used for phenotyping, the images processed by the LAMINA software, raw and processed leaf physiognomy metrics, and genotypic BLUP values, in addition to the raw GWAS output tables. Scripts used to generate the results presented here, including a BLUP pipeline for preparing phenotype data for GWAS, are available at https://github.com/bschiffthaler/aspen-v2 and at https://github.com/sarawestman/Genome_paper.

## Results and discussion

### A high-quality reference genome for *Populus tremula*

The previously available version of the *P. tremula* genome (v1.1) was highly fragmented despite having good representation of the gene space (Lin *et al*., 2018; Supplementary table 3). Such fragmentation was a common characteristic of assemblies produced using short-read sequencing technologies and was especially problematic for repeat-rich and highly heterozygous genomes. The extent of fragmentation prohibited or limited analyses requiring long-range contiguity, such as synteny, made gene-family analysis error-prone and presented challenges for accurate SNP calling in hard to assemble regions. Here, we used a combination of long-read sequencing, optical and genetic maps to generate a high-quality and highly contiguous genome assembly for *P. tremula.* Integration with a genetic map Apuli *et al*. (2020) enabled anchoring and orienting of assembled contigs to form pseudo-chromosomes (Figure 1A), with the final assembly having a contig N50 of 16.9 MB (Table 1), representing an order of magnitude improvement compared to the previous v1.1 genome assembly (Lin *et al*., 2018).

**Figure 1.**
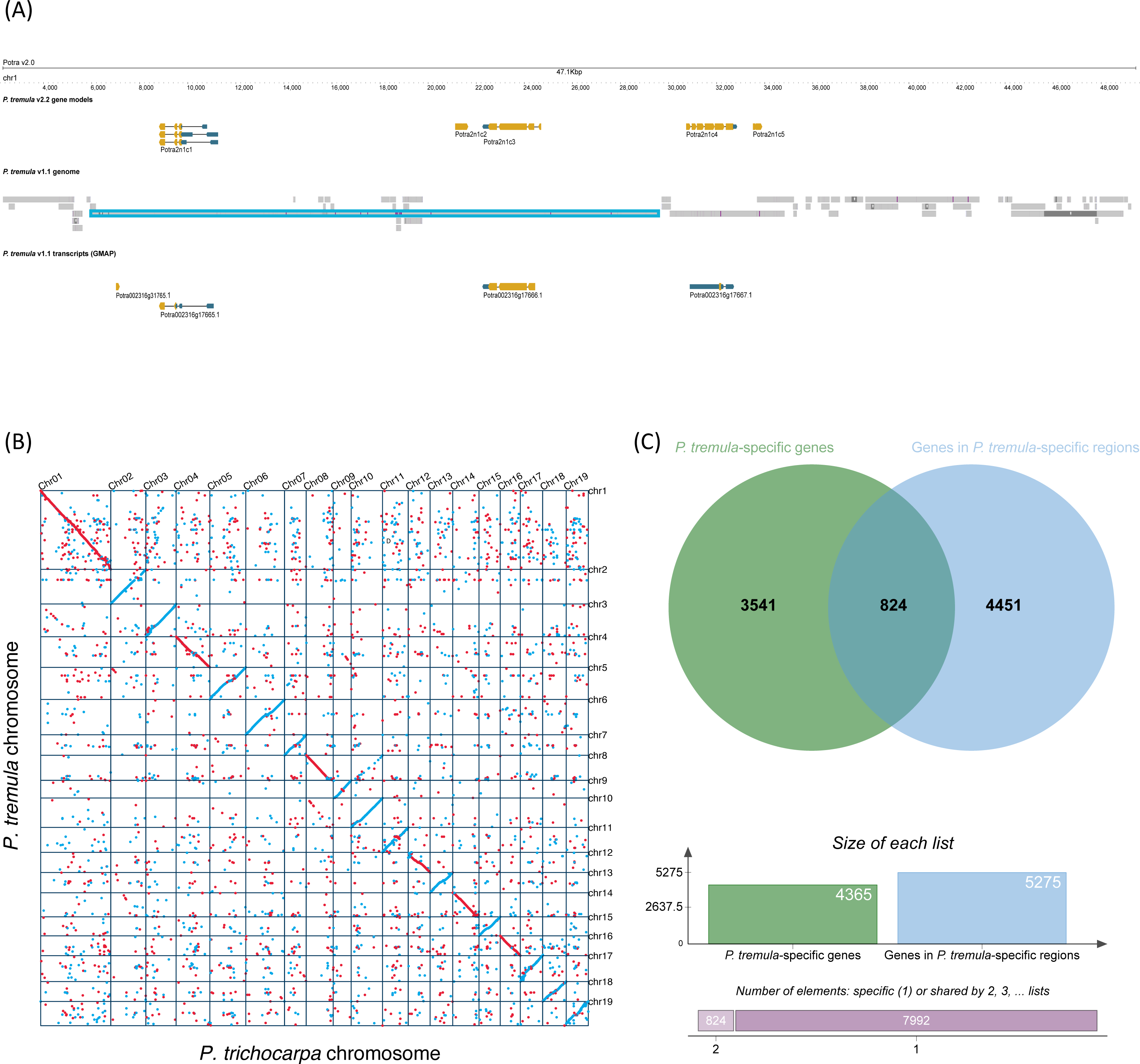
Overview of the *P. tremula* v2.2. genome. **(A)** Comparison of a 47.1 Kbp region of *P. tremula* chromosome 1 showing the *P. tremula* v2.2 gene models and a liftover of the *P. tremula* v1.1 genome and transcripts, rendered in the JBrowse tool in PlantGenIE. The region in turquoise highlights an example of a longer scaffold in *P. tremula* v1.1 containing a gene. **(B)** Synteny and structural rearrangements between *P. tremula* and *P. trichocarpa*. **(C)** Venn diagram created using the Venn tool in PlantGenIE, showing the intersection of genes in *P. tremula-*specific regions and the *P. tremula-*specific genes identified from synteny analysis.

**Table 1.** Summary statistics for *Populus tremula* genome assemblies v1.1 (Lin *et al*., 2018) and v2.2. GC content statistics were calculated using the unmasked genome.

| Statistic | *P. tremula* v1.1 | *P. tremula* v2.2 |
| --- | --- | --- |
| # contigs (>= 0 bp) | 204318 | 1601 |
| # contigs (>= 1000 bp) | 31632 | 1584 |
| # contigs (>= 5000 bp) | 7267 | 1339 |
| # contigs (>= 10000 bp) | 5151 | 986 |
| # contigs (>= 25000 bp) | 3209 | 491 |
| # contigs (>= 50000 bp) | 1789 | 255 |
| Total length (>= 0 bp) | 386236512 | 408834716 |
| Total length (>= 1000 bp) | 328536064 | 408824553 |
| Total length (>= 5000 bp) | 277117215 | 407999588 |
| Total length (>= 10000 bp) | 262322877 | 405364617 |
| Total length (>= 25000 bp) | 231504505 | 397478443 |
| Total length (>= 50000 bp) | 180499961 | 389097052 |
| # contigs | 12044 | 1489 |
| Largest contig | 418873 | 53234430 |
| Total length | 294670244 | 408605800 |
| GC (%) | 33.56 | 33.87 |
| N50 | 69979 | 16928776 |
| N75 | 29987 | 13637973 |
| L50 | 1227 | 9 |
| L75 | 2826 | 15 |
| # N's per 100 Kbp | 5428.58 | 6573.91 |
| Reads aligned (%) | 97.77% | 96.40% |
| Reads properly paired (%) | 92.33% | 94.19% |

### Genome Assembly

The v2.2 *P. tremula* genome assembly contains 19 pseudo-chromosomes and 1,582 unplaced scaffolds with a combined length of 408,834,716 bp and an N50 of 16.9 Mb (Supplementary table S3). Alignment of ∼95 million Illumina reads (∼20X coverage) yielded a mapping rate of 96.4% (compared to 97.77% in v1.1) with 94.19% (compared to 92.33% in v1.1) of paired-end reads mapped as proper pairs. The increase in proper pairs and decrease in overall mapping reflects expectations for an assembly with higher contiguity but lower per-base accuracy, which is a characteristic of the PacBio sequencing reads utilised. Analysis of the genome using Benchmarking Universal Single-Copy Orthologue (BUSCO) with the embryophyta_odb10 ortholog set (Simão *et al*., 2015) to assess gene-space completeness identified 96% (96% in v1.1) complete BUSCOs, of which 81.7% (82.5% in v1.1) were single copy and 15.1% (14.3% in v1.1) duplicated (Supplementary table S3). The long terminal repeat (LTR) index for the assembly (Ou *et al*., 2018) was 6.65, with 1.42% of intact LTRs and 20.66% of total LTRs, indicative of a high-quality assembly. The improved contiguity of the new assembly is clear when examining multiple sequences from v1.1 that align, for example, to a region of chromosome 1 (Figure 1A).

### Gene annotation

There are 39,894 identified gene models, 37,184 of which are located on pseudo-chromosomes and 2,710 on unplaced scaffolds. There are 77,949 annotated transcripts, 73,765 on pseudo-chromosomes and 4,184 on unplaced scaffolds (∼1.95 transcripts per gene). Functional annotations were assigned for 73,765 transcripts in 37,184 genes. Analysis of the predicted transcripts using BUSCO with the embryophyta_odb10 ortholog set showed 98.1% (96.8% in v1.1) complete BUSCOs, of which 35.7% (30.2% in v1.1) were single copy and 62.4% (66.6% in v1.1) duplicated (Supplementary table S3). Similarly, the PLAZA core Gene Family (coreGF) set of genes (Veeckman *et al*. 2016; Bucchini *et al*., 2021) indicated 99% completeness (99% in v1.1; Supplementary table S3).

### Comparative genomics analyses

We utilised the improved assembly to perform gene family and comparative genomics analyses, identifying syntenic and species-specific genomic regions of *P. tremula* compared to *P. trichocarpa.* (Figure 1B). There were a large number of aspen- and *P. tremula*-specific genes and genomic regions (Supplementary table S4) in addition to a set of highly diverged regions (Supplementary table S5), although we acknowledge that lineage specific (orphan) genes should be viewed with caution (Weisman *et al*., 2020). Similar analyses to identify orphan genes in *P. trichocarpa* were recently reported (Yates *et al*., 2021), but using a far more stringent definition of orphan genes, showing that orphan genes are polymorphic in a GWAS population and integrated within co-expression networks. Species- and clade-specific gene families were identified (Supplementary tables S6), and *P. tremula*-specific genomic regions were enriched for the terms “non-membrane-bounded organelle” and “cell differentiation” among the set of expanded gene families (Supplementary tables S7 S8, S9). We used the tools in PlantGenIE to explore these groups of genes and their gene ontology (GO) enrichments. For example, we used the Venn tool to view the intersection of lists of *P. tremula* specific genes and genes in *P. tremula*-specific regions (Figure 1C). While this exploration did not provide us with specific insights into leaf physiognomy traits, which we focus on below, the comparative genomics resource is available for exploration in future studies of *P. tremula*.

### Long intergenic non-coding RNAs

Long non-coding RNAs (lncRNAs) are arbitrarily defined as transcripts longer than 200 nt, not producing functional proteins. If they are located entirely in the intergenic space, they are sub-classified as long intergenic non-coding RNAs (lincRNAs). In general, lncRNAs have low expression levels and tissue-specific expression. They are also characterised by a rapid evolution and a low sequence conservation between species (Chen & Zhu, 2022; Palos *et al*., 2023). Recent reports have shown that lncRNAs participate in plant developmental regulation (Kramer *et al*., 2022; Chen *et al*., 2023b). We identified 902 putative lincRNAs in developing aspen leaves (Supplementary table S10) and integrated them into the Aspen Leaf (AspLeaf)expression data resources at PlantGenIE.org.

### Population genetics of SwAsp, UmAsp and ScotAsp

The original locations of the samples (Figure 2A) differed among the aspen collections in climatic variables, with Scottish samples drawn from a milder, maritime climate and Swedish samples from a colder, more continental climate (Supplementary table 11). Based on whole-genome re-sequencing data, and after removal of related samples and the batch correction described in Rendón-Anaya *et al*. (2021), we identified 12,054,692 SNPs for 99 individuals from SwAsp, 16,938,820 SNPs for 227 individuals from UmAsp, and 19,655,602 SNPs for 105 individuals from ScotAsp, on chromosomes, after discarding SNPs with missing rate >5% and failing the Hardy–Weinberg equilibrium test (*P*-value < 1e^-6^). Of these SNPs, 27.4% were found within gene boundaries for SwAsp, 31.1% for UmAsp, and 31.8% for ScotAsp while 33.6% were in gene flanking regions for SwAsp, 30.8% for UmAsp, and 30.6% for ScotAsp. The remaining sites were in intergenic regions. The SNP density was 33.3 SNPs/Kbp for SwAsp, 48.8 SNPs/Kbp for UmAsp and 54.3 SNPs/Kbp for ScotAsp across the 19 chromosomes and was highest in the flanking regions and lowest in the CDS regions for the three aspen collections. The three collections harboured substantial levels of nucleotide diversity (π) across the genome (0.0061 in SwAsp, 0.0080 in UmAsp, and 0.0082 in ScotAsp). While the majority of SNPs (13,153,803) were shared between Swedish (UmAsp or SwAsp) and Scottish aspens (Figure 2B), 9,539,889 SNPs were shared only between SwAsp and UmAsp, indicating the potential utility of a combined Swedish aspen resource, and there were 6,501,799 SNPs unique to ScotAsp, highlighting its differences from the Swedish collections.

**Figure 2.**
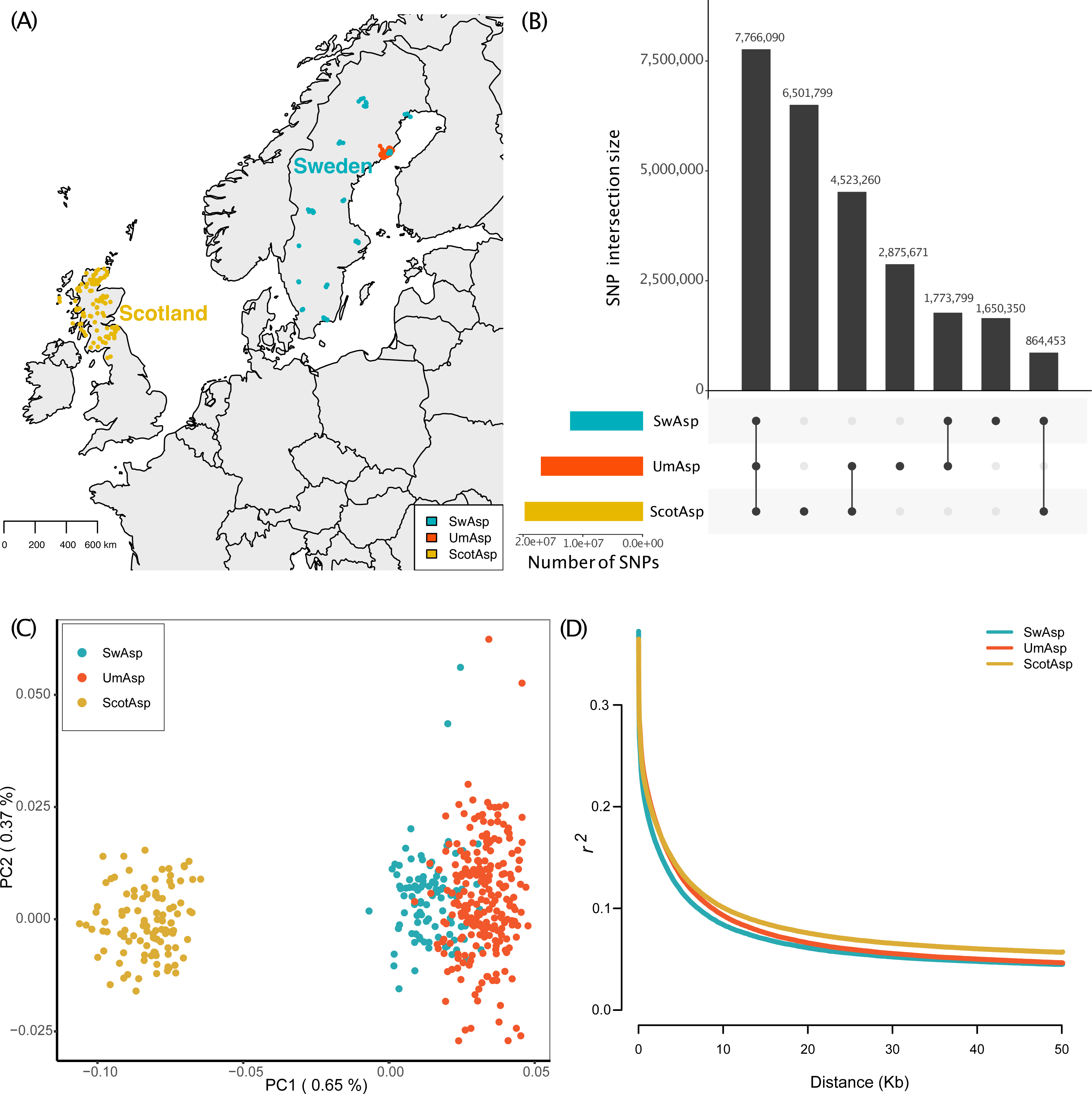
Overview of genome-wide association study using three aspen collections. **(A)** Map indicating the original sampling locations of the individual wild trees in the aspen collections from Scotland (ScotAsp), Sweden (SwAsp) and the Umeå municipality in Sweden (UmAsp) that were included the genetic analyses after removal of related samples. **(B)** Number of biallelic SNPs, filtered by Hardy-Weinberg Equilibrium *P-*value > 1e^-6^ and missingness < 5%, in the ScotAsp, SwAsp and UmAsp collections. Coloured bars on the left indicate total number of SNPs in each collection, linked points indicate membership of intersections among collection, with numbers in intersections shown in the vertical bars above. Single points indicate sets of SNPs exclusive to one population. **(C)** Principal components plot of the first two principal components (PCs) of pruned, unrelated SNPs (LD *r^2^* <0.2) to show population structure in the ScotAsp, SwAsp and UmAsp collections. Proportion of variance explained by each PC is indicated in parentheses. **(D)** Rates of linkage disequilibrium decay in the ScotAsp, SwAsp and UmAsp collections.

We used seven measures calculated in 10 Kbp non-overlapping windows to identify regions under selection in SwAsp and UmAsp using ScotAsp as an outgroup. Signatures of positive selection were identified for 589 and 653 regions, corresponding to 7.46 Mbp and 8.60 Mbp in SwAsp and UmAsp, respectively (Table S12). Only 1.57 Mbp of regions under selection were shared between SwAsp and UmAsp. Based on genome annotation, we identified 621 and 633 genes under selection in SwAsp and UmAsp, respectively (Table S12) of which 123 genes were in common.

Population structure based on PCA clearly separated at least two independent clusters of individuals, one corresponding to ScotAsp with the other comprising SwAsp and UmAsp (Figure 2C), indicating that ScotAsp is a suitable outgroup to identify signatures of selection in SwAsp and UmAsp. This clustering pattern is consistent with previous observations by de Carvalho *et al*. (2010) and Rendón-Anaya *et al*. (2021), which have shown that the aspens from the British Isles are diverged from aspens in continental Europe. In agreement with previous results (Lin *et al*., 2018), genome-wide mean linkage disequilibrium (LD) measured by *r*^2^ was largest between adjacent SNP pairs (0.36 to 0.37) in the three aspen groups and decreased rapidly to 0.1 within 10 Kbp (Figure 2D). The population genetic data are available in JBrowse in PlantGenIE, where tracks can be loaded and viewed in the context of other genomic features and significant GWAS results.

### Natural genetic variation in leaf physiognomy phenotypes

The leaves of *P. tremula* are rounded with irregular serrations, hereafter termed indents. In previous leaf physiognomy analyses we reported natural genetic variation in ten traits (Bylesjö *et al*., 2008), and three representative traits (‘leaf area’, ‘circularity’, and ‘ident depth’; Mähler *et al*., 2020) in the SwAsp collection. Here we present 26 traits (Appendix S2) measured in the two SwAsp common gardens in each of two years, and in the UmAsp and ScotAsp common gardens in a single year. The raw image files of the sampled leaves, the annotated images indicating measured parameters (Figure 3A) and data output from the measurement software (LAMINA, Bylesjö *et al*., 2008), are available at the associated Figshare data repository (see data availability statement for details). There was a clear shared genetic component among indent traits and shape traits (Figure 3B), with high genetic correlations between ‘Squared Perimeter/Area’ and each of ‘indent depth’ and ‘indent depth standard deviation (SD)’. Leaf size traits were all positively genetically correlated, with high correlations among length traits and among width traits, and to a moderate extent between length and width traits (Figure 3B). Narrow-sense (‘chip’) heritability was greater for shape and indent traits than size traits (3C, Appendix S2), with the exception of the composite trait ‘indent density,’ which had low heritability (*h*^2^ = 0.138). There was clear separation of size traits from shape and indent traits in the first principal component (PC1) of a PCA of all 26 leaf metrics in the three populations. While PC1 explained 43.5 % of the variation in this combined data set and an overall intersection among ScotAsp, SwAsp and UmAsp, there was a tendency towards larger leaves (i.e. smaller values of PC1) in ScotAsp. The heritability estimates and shared multivariate space among the three aspen collections, together with number of common SNPs, favour the integration of these traits and collections in genetic analyses. The processed phenotype data, including composite leaf physiognomy traits (Appendix S2) and BLUPs are available at Figshare.

**Figure 3.**
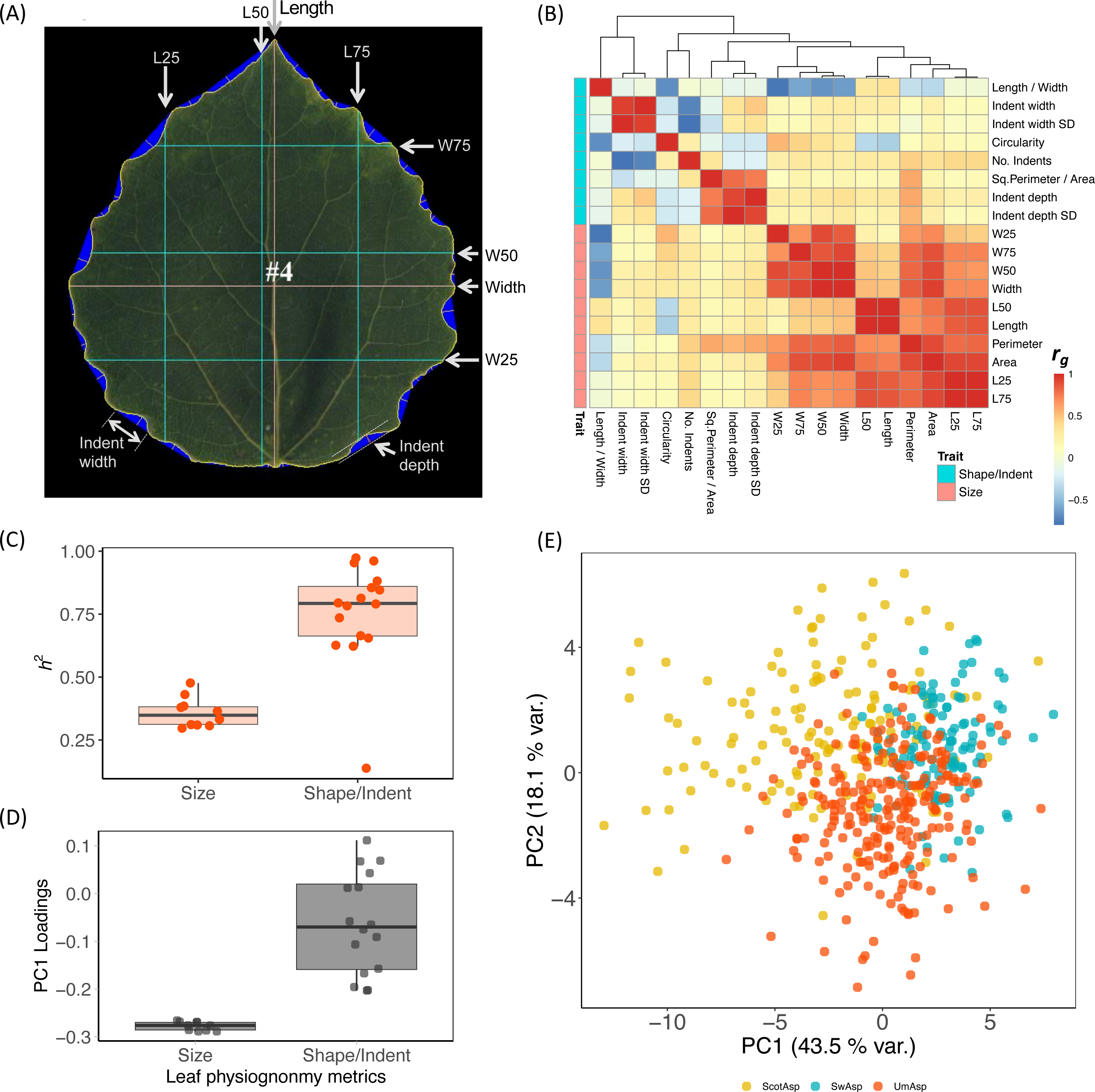
Overview of leaf physiognomy metrics. **(A)** Example processed leaf image from LAMINA software, with annotations indicating the Indent width, Indent depth, the Length and Width axes, and length and width at 25% (L25, W25) and 75% (L75 and L25) along each perpendicular axis. **(B)** Heatmap of genetic correlations of measured leaf physiognomy traits in the Umeå aspen (UmAsp) collection. Composite traits are excluded to reduce redundancy. Scale bar indicates genetic correlation *r_G_* values. Coloured bars indicate category of either ‘Size’ (leaf shape metrics) or ‘Shape/Indent’ (size and indent metrics). Hierarchical clustering between the clusters uses the complete linkage method. **(C)** Marker-based heritability, *h2,* of 26 shape and size/indent leaf physiognomy metrics in the UmAsp collection. All trait metrics are described in Appendix S1. **(D)** Principal components loadings plot indicating the loading scores of size and shape metrics indicated in Appendix S2 and the associated Principal Components Analysis (PCA) plot **(E)** for 26 leaf physiognomy metrics in Scottish (ScotAsp), Swedish (SwAsp) and UmAsp (UmAsp) collections. Proportion of variance explained by each component is in parenthesis. In all cases, means are omitted for Indent length and Indent width to avoid redundancy, since medians are included for these traits.

### GWAS in open chromatin regions enhances detection of SNP-phenotype associations

Leaf physiognomy traits appear to be highly polygenic, yet highly heritable, with variation among individuals resulting from numerous small-scale effects (Mähler *et al*., 2020). In such cases it is common that no significant genetic associations are identified, with huge sample sizes needed to detect such small-scale effects. Other factors, such as incomplete genome assembly, can also prohibit detection of sequence-based genetic markers in hard-to-assemble regions of the genome. While we previously reported our GWAS study in three leaf physiognomy traits in SwAsp using the previous genome assembly version, here we conducted GWAS on 26 leaf physiognomy metrics in each of the SwAsp, UmAsp and ScotAsp populations, taking advantage of the substantially higher number of SNP markers called using the improved v2.2 genome assembly. In the GWAS including all SNPs (All-SNP GWAS), we detected significant (*q-*value < 0.05) associations for vertical size 75% (L75) in UmAsp, while in ScotAsp there were associations for ‘indent width SD’ (the standard deviation of indent width) and circularity (Supplementary table S13). No significant associations were identified in the SwAsp All-SNP GWAS. A GWAS that includes several million SNPs in a relatively small population may fail to detect associations for complex traits due to adjustments for multiple testing. Inspired by work in maize (Rodgers-Melnick *et al*., 2016) demonstrating that open chromatin regions harbour much of the genetic variation for quantitative traits, we generated ATAC-Seq data from *P. tremula* leaves to identify regions of open chromatin (open chromatin regions, OCRs). We then subset SNPs to only those regions, and ran GWAS using these SNP subsets (OCR GWAS). This resulted in 212,902 SNPs in ScotAsp, 185,616 in SwAsp and 220,009 in UmAsp. The genomic context distribution differed between the All-SNP and OCR-SNP sets with a greater proportion of SNPs in up/downstream, UTR and exonic regions, and a lower proportion of SNPs in intergenic regions, in the OCR-SNP set (Figure 4). OCR GWAS resulted in ten, five and four significant associations (*q*-value > 0.05) in ScotAsp, UmAsp and SwAsp respectively for three, two and two leaf traits respectively (Supplementary table S13). These associations ranked highly in the All-SNP GWAS, despite in most cases falling below the *q-*value threshold (Table 2). Significant OCR-GWAS associations intersected with significant All-SNP GWAS associations in the case of only one ScotAsp trait (indent width SD).

**Figure 4.**
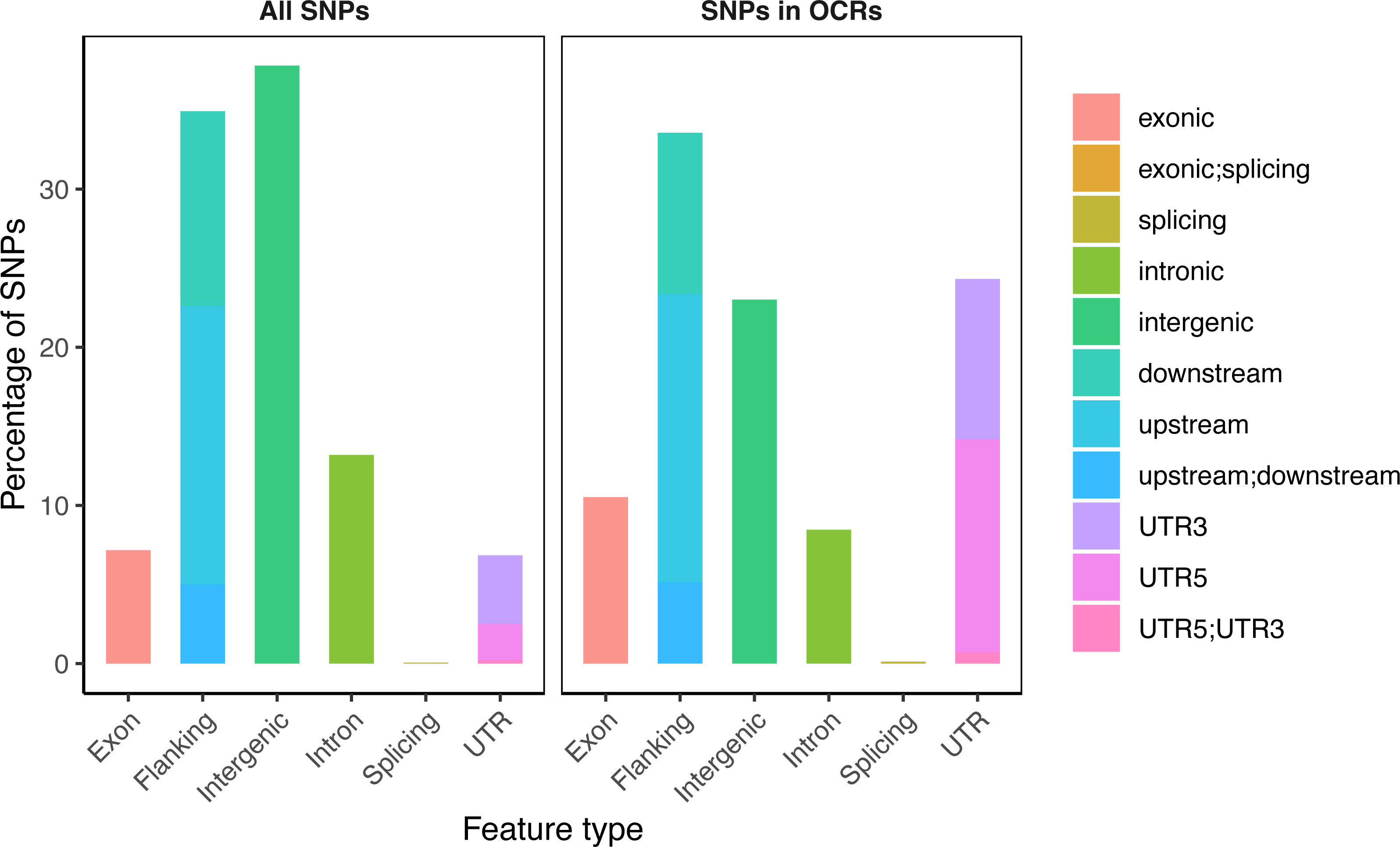
Comparison of the percentage of SNPs located in different genomic contexts in two GWAS backgrounds in the SwAsp collection. GWAS was first conducted using “All SNPs” (all the genome-wide SNPs filtered on SNP quality including Excess Heterozygosity, Hardy Weinberg *P*-value, and minor allele frequency > 0.05) (left panel). The set of 6,806,717 “All SNPs” was filtered to those only those 185,616 SNPs in open chromatin regions (“SNPs in OCRs”, right panel). The percentage of SNPs in each genomic context category was calculated from the total number in the set used for the GWAS. Genomic contexts were assigned using ANNOVAR with flanking regions defined as 2000 bp.

**Table 2.** Comparison of of SNP-trait associations ranked by signficance (by association *P-* value) in the genome wide association (GWAS) analysis of leaf physiognomy traits in the ScotAsp, SwAsp and UmAsp collections. For each association significant at *q-*value < 0.05, the rank of the SNP is shown in the OCR GWAS (GWAS using SNPs filtered to open chromatin regions) compared to the rank in the All-SNP GWAS (using SNPs filtered only by excess heterozygosity, Hardy-Weinberg Equlibrium *P-*value and minor allele freqency, see materials and methods for details).

| Population | Trait | SNP | OCR GWAS rank | All-SNP GWAS rank |
| --- | --- | --- | --- | --- |
| ScotAsp | Indent density | chr9_10684216_G_A | 1 | 3 |
|  |  | chr9_10684306_T_A | 2 | 4 |
|  |  | chr9_10684389_C_T | 3 | 5 |
|  | Indent depth SD | chr9_10684114_T_G | 1 | 2 |
|  |  | chr9_10684216_G_A | 2 | 4 |
|  |  | chr9_10684306_T_A | 3 | 5 |
|  | Indent width SD | chr9_10684216_G_A | 1 | 15 |
|  |  | chr9_10684306_T_A | 2 | 16 |
|  |  | chr9_10684389_C_T | 3 | 19 |
|  |  | chr9_10646677_G_C | 4 | 90 |
| SwAsp | Length 75% / Width | chr5_13736867_T_G | 1 | 5 |
|  |  | chr5_13736875_A_C | 2 | 7 |
| SwAsp | Indent width SD | chr1_31893859_G_A | 1 | 6 |
|  |  | chr1_31894106_A_T | 2 | 7 |
| UmAsp | Indent density | chr16_10033126_C_G | 1 | 4 |
|  |  | chr16_10033093_C_T | 2 | 10 |
|  |  | chr16_10033097_C_T | 3 | 11 |
|  |  | chr16_10032951_G_T | 4 | 15 |
| UmAsp | Squared Perimeter/ Area | chr11_432652_C_A | 1 | 2 |

### Genome-wide associations suggestive of leaf development processes

We looked for signals of leaf developmental processes in the GWAS results, first by examining expression patterns in the AspLeaf data set for all genes with significant SNPs in the GWAS. Of the 25 genes associated with SNPs in the GWAS, 24 had expression data in the AspLeaf data resource, and 18 of those had a clear gradient of expression from the apex or youngest leaf to the oldest leaf. Next, we examined the annotations of genes associated with significant SNPs in both the All-SNP and OCR GWAS and noted that the majority are annotated with functions that include plant developmental processes (Table 3). Significant SNPs for ScotAsp indent width SD in the All-SNP GWAS (Supplementary table S13) included a pescadillo homologue (Potra2n4c9542), important in leaf growth, in particular through control of ribosomal biogenesis affecting leaf cell division, expansion, and pavement cell differentiation and (Cho *et al*., 2013; Ahn *et al*., 2016), and a *FAB1*-like gene (Potra2n1c1433) also annotated as a phosphatidylinositol-3-phosphate 5-kinase, important for auxin signalling and normal plant development (Hirano *et al*., 2011; Baute *et al*., 2015). The three significant SNP associations for ScotAsp circularity in the All-SNP GWAS were in linkage disequilibrium and located on chromosome 17 in intronic and exonic regions of Potra2n17c30934, which is annotated as a “PATRONUS 1-like isoform X1 protein” and carries the GO identifier, “regulation of mitotic cell cycle.” PATRONUS1 is reported to have an important role in cell division in plants (Cromer *et al*., 2019). In the UmAsp All-SNP GWAS, the 11 significant SNPs were located in upstream, exonic, intronic and 5’ UTR regions of Potra2n3c7046 and in an intergenic region with Potra2n3c7047 (Supplementary table S13). The *A. thaliana* homologues of these two genes are, respectively, AT5G13390 (‘No exine formation 1’), important in pollen wall development (Ariizumi *et al*., 2004), and AT1G28130, an Auxin-responsive GH3 family protein that regulates auxin metabolism and distribution and plant development (Zheng *et al*., 2016; Guo *et al*., 2022). In the SwAsp OCR-GWAS, two SNPs in the 5’ UTR region of Potra2n1c2680 were associated with indent width SD. The *A. thaliana* homologue of Potra2n1c2680 is involved in several plant developmental processes (Xiao *et al*., 2021).

**Table 3.**
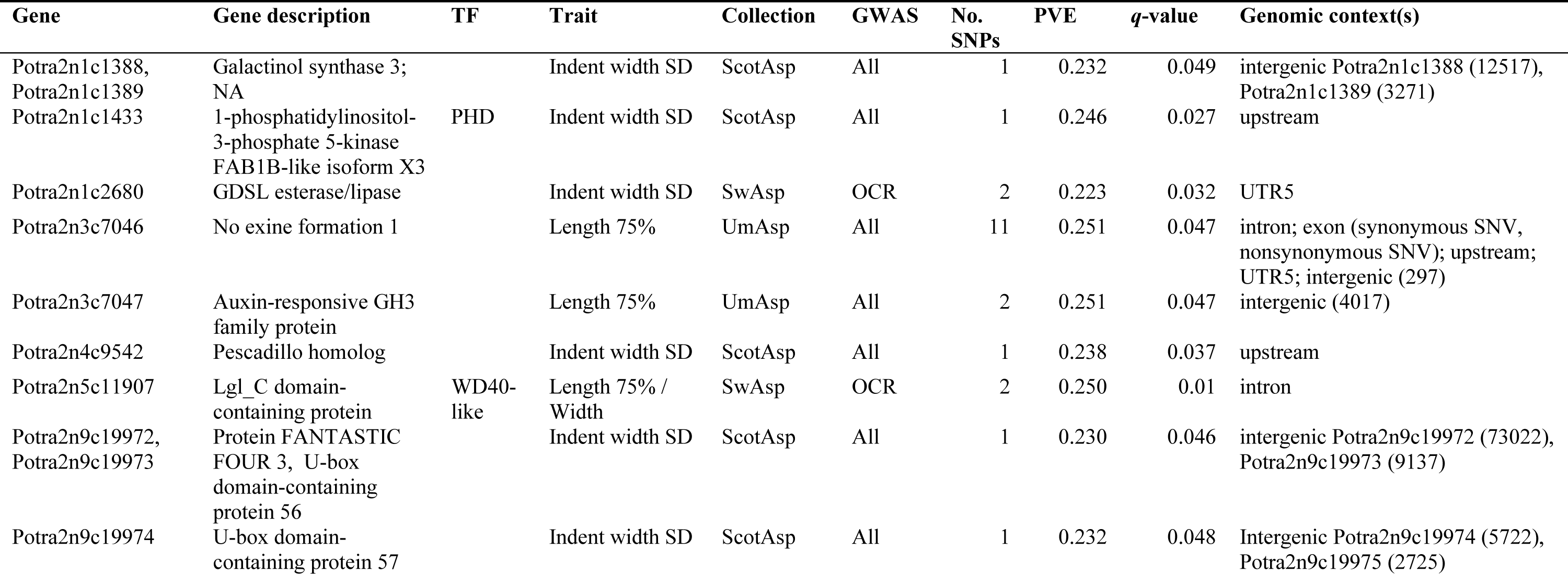
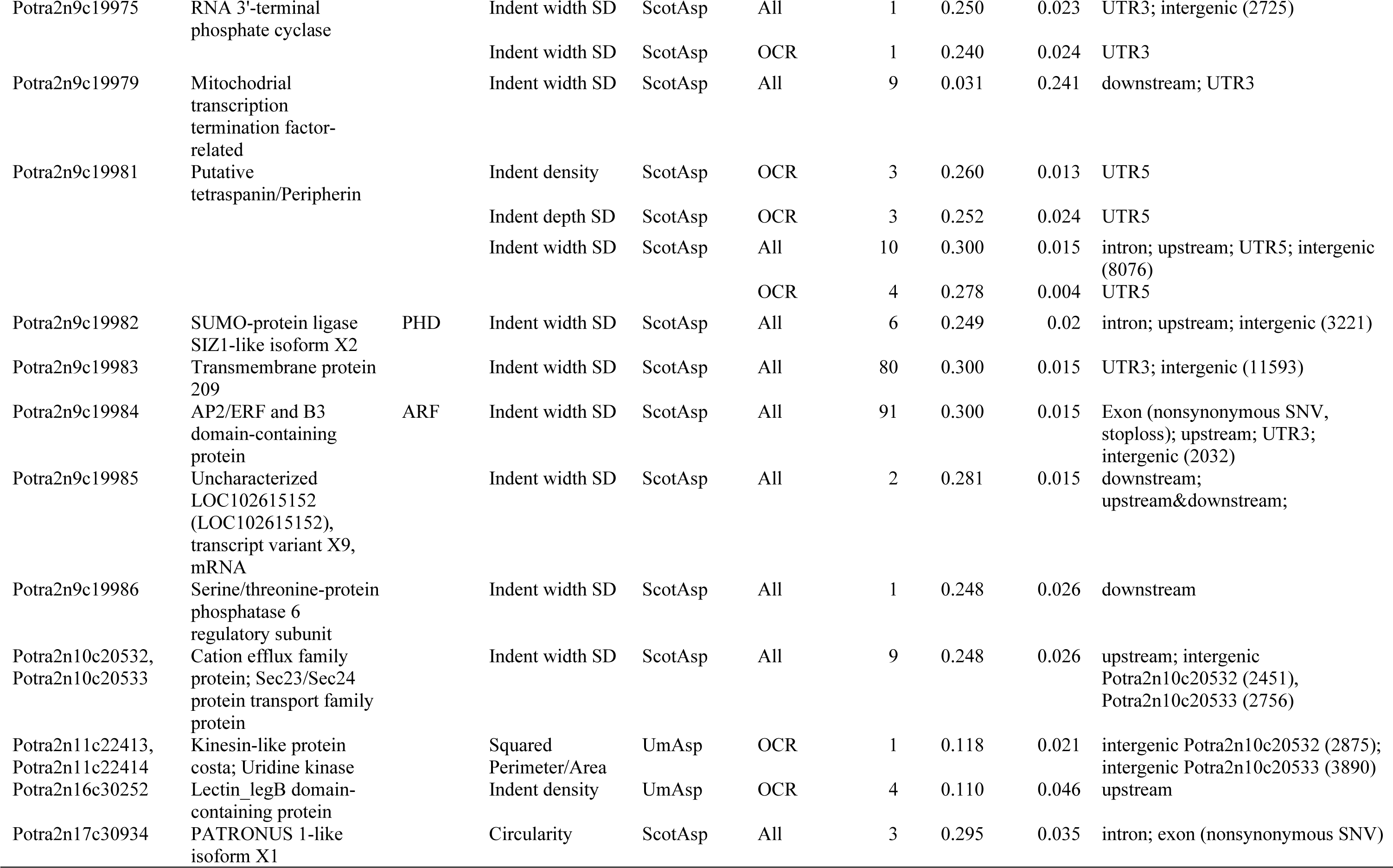
Summary of significant (*q*-value < 0.05) association mapping statistics of leaf physiognomy traits in separate Genome-Wide Association Studies (GWAS), in each of the Swedish (SwAsp), Umeå (UmAsp) and Scottish (ScotAsp) aspen collections. Traits are described in detail in Appendix S2. ‘Gene’ = *Potra* v2.2 gene model associated with the genomic context of the significant SNP(s). ‘Gene Description’ and ‘TF’ respectively indicate the functional description of the gene and its transcription factor family (if applicable). ‘GWAS’ indicates the SNP background for the GWAS: ‘All’ = all filtered (minor allele frequency >0.05) SNPs in the genome; ‘OCR’ = all SNPs subset to only those in open chromatin regions. ‘No. SNPs’ = number of significant SNP-phenotype associations at *q-*value < 0.05. ‘PVE’ = maximum proportion of phenotypic variation explained by an individual SNP associated with the gene and trait. ‘*q*-value’ = minimum association *q*-value for any SNP associated with the gene and trait. ‘Genomic context’ = position of the gene relative to genomic features; if intergenic the minimum distance (bp) is stated.

### Combined data resources reveal variation in leaf base angle

Two SNPs from the SwAsp OCR GWAS were associated with leaf base angle (W75/Width). Both of these SNPs are located in an intron of Potra2n5c11907 (Supplementary Figure 3A), which is annotated as LLGL scribble cell polarity complex, a transcription factor that in *A. thaliana* is a Transducin/WD40 repeat-like superfamily protein (AT4G35560). The *WD40*-like transcription factors have roles in various developmental processes including organ size determination (Gachomo *et al*., 2014; Guerriero *et al*., 2015; Yang *et al*., 2018). The phenotypic BLUP values for leaf base angle were significantly partitioned by the chr5_13736867 SNP allele groups in the SwAsp collection (Supplementary Figure 3B), and the individuals with the greatest and smallest phenotypic values (SwAsp 114 and SwAsp 4, which belong to the contrasting allele groups), can be identified using the phenotype files at the Figshare data repository. The cropped LAMINA output images of these can be downloaded from Figshare and leaf shape features compared. In this particular case, the two genotypes differed markedly in leaf base angle (Supplementary Figure 3B). The phenotypic values were, however, somewhat variable for the allele groups for this SNP, which is consistent with the polygenic nature of leaf shape determination. As such, not all ScotAsp genotypes with two recessive alleles for this SNP had a steep leaf base angle. Following a similar approach, the expression of Potran2n5c11907 in the data set from developing SwAsp leaf buds can be partitioned by SNP alleles for chr5_13736867 (Supplementary Figure 3C). While in this case the interpretation is not straightforward, it serves to demonstrate the integration of available data resources to explore characteristics of identified candidate genes and to help prioritise among candidates.

### A 177-kbp region associated with leaf shape phenotypes in Scottish aspen

The All-SNP associations for ScotAsp indent width SD included 122 SNPs on chromosome 9, which intersected with significant SNPs in the ScotAsp OCR GWAS for the same trait, as well as indent density and indent depth SD (Supplementary table S13). These SNPs were located within a region spanning ∼177 kbp on chromosome 9 (Supplementary Figure 3). Many of the SNPs in this region were in linkage disequilibrium (LD) and of the 15 genes in this region, 12 were associated with SNPs in the ScotAsp GWAS at *q-*value < 0.05 (Supplementary table S13; Table 3). The proportion of variance explained (PVE) by any single SNP among these significant associations was moderate, ranging from 0.23 to 0.30 (Supplementary table S13). These SNPs were distributed across various genomic contexts in the 12 genes, all with functions suggestive of roles in leaf development. These included a *SIZ1*-like isoform that is a *PHD* transcription factor (Potra2n9c199982), involved in cell division and expansion (Catala *et al*., 2007; Miura *et al*., 2010; Mouriz *et al*., 2015), an *ARF10* auxin response factor (Potra2n9c199984) involved in auxin signalling during leaf development (Hendelman *et al*., 2012; Liu *et al*., 2007; Ben-Gera *et al*., 2016), and two periphrins/tetraspanins (Potra2n9c199975, Potra2n9c199981), involved in numerous cell proliferation and tissue patterning processes (Wang *et al*., 2015; Reimann *et al*., 2017). Expression in the AspLeaf dataset, as observed using the exImage tool at PantGenIE.org, showed a gradient of relative expression across the developmental stages of the terminal leaves in eight of these 12 genes, which was most pronounced for Potra2n9c199975, Potra2n9c199981, Potra2n9c199982, Potra2n9c199984 and Potra2n9c199985. This suggests that these genes are developmentally regulated. While the 12 genes were significantly associated with only four traits, each of the genes occurred in the top-ranked 1000 genes of at least three, and up to 13, ScotAsp traits (Supplementary table S13), suggesting that these genes contribute to multiple leaf physiognomy phenotypes in Scottish aspens. In contrast to ScotAsp, these 12 genes were not highly ranked in the SwAsp and UmAsp GWAS and would thus appear to make a negligible contribution to leaf size and shape variation in Swedish aspens. In SwAsp only Potra2n9c199985 and Potra2n9c19972 were present in the top-ranked 1000 genes for two traits, Area and Length:Width ratio respectively (Supplementary table S13), and for UmAsp, none of these 12 genes were ranked in the top 1000 genes. This reflects the demographics of SNPs at this locus in the other collections; of the 122 significant SNPs for this trait in ScotAsp, only 25 SNPs were present in the SwAsp in the SwAsp VCF, with a median minor allele frequency (MAF) of 0.122, and while 119 of the 122 SNPs were present in UmAsp the median MAF was 0.092, indicating that the variation at these sites is higher in ScotAsp (median MAF = 0.302). Only two of the top 1000 genes for Indent width SD for this trait intersected among all three collections, however these (Potra2n1c1769 and Potra2n3c8236) did not have apparent annotations relevant to leaf developmental processes. This example suggests that there is substantial control of natural variation in leaf shape phenotypes determined by SNPs at this locus on chromosome 9 and that this is specific to Scottish aspen. Since ScotAsp separates from SwAsp and UmAsp in the SNP PCA (Figure 2C), it is not unexpected that the complex leaf phenotypes in the Swedish and Scottish populations do not share this GWAS locus.

### Combined resources aid genomic exploration of SNP-phenotype associations

Many of the gene annotations in the GWAS results have plausible biological links to leaf physiognomy traits. However, these interpretations are speculative, especially for associations of relatively low PVE and where few individuals are homozygous for the minor allele. In such cases it can be useful to consult several lines of evidence to evaluate the plausibility of the functional link. To demonstrate utility of the Potra v2.2 genomics data available within PlantGenIE.org for such explorative analyses, we examined the associations to Potra2n10c20533, one of the genes associated with indent width SD in the ScotAsp All-SNP GWAS, appearing as a small peak on the Manhattan plot for this trait (Figure 5A). The annotation of Potra2n10c20533 is a putative protein transport protein Sec24A, with the most sequence-similar gene in *A. thaliana* (AT3G07100) having a role in endoplasmic reticulum maintenance and cell size regulation in sepals (Nakano *et al*., 2009; Qu *et al*., 2014). The Potra2n10c20533 gene harbours a significant SNP 1962 bp upstream from the annotated transcription start site, and eight SNPs in the intergenic region between Potra2n10c20532 (a Cation efflux family protein associated with manganese tolerance in *A. thaliana;* Peiter *et al*., 2007) and Potra2n10c20533 (Supplementary table S13). To reveal the potential biological function of these associations, we used the exNet tool at a lenient threshold *P*-value 10^-1^ to identify first degree neighbours of Potra2n10c20533 in the AspLeaf dataset co-expression network (Figure 5B). Functional enrichment of these co-expressed genes identified GO categories for cell expansion (Figure 5C). The network visualisation using exNet (Figure 5B) showed that the set of 205 co-expressed genes included 29 transcription factors (TFs; i.e. 14 % were TFs) and two lincRNAs. Using the gene expression visualisation tools available at PlantGenIE.org we explored the expression of these lincRNAs within the AspLeaf datasets, revealing a gradient of expression across the terminal leaf development stages (Figure 5D). This revealed that the two lincRNAs were negatively correlated to more than 100 of the first-degree neighbours of Potra2n10c20533. Use of the JBrowse tool at PlantGenIE.org also enabled us to view the significant SNPs in the GWAS region around Potra2n10c20533 in the context of the Potra v2.2. gene models and co-locating eQTL (Figure 5E). Mapping of eQTL was conducted using two different methods; using Matrix eQTL, we identified 466,966 significant (FDR < 0.05) eQTL, whereas the more conservative method using fastJT identified 173,080 significant eQTL (Supplementary table S14). The JBrowse tool enabled the easy visualisation of *trans* eQTL acting on ten genes identified using Matrix eQTL that co-located with intergenic SNPs in the GWAS between Potra2n10c20532 and Potra2n10c20533. Use of the Enrichment tool of PlantGenIE.org for this set of ten *trans* eQTL genes showed Pfam enrichment terms for categories relevant to plant organ development (Figure 5F), including Phosphatidylinositol-4-phosphate 5-Kinases (Watari *et al*., 2022), MORN repeats (Lee *et al*., 2010), and K-Box regions (Uchida *et al*., 2007). In the same intergenic region, there was one *cis* eQTL (FDR = 0.023), for the expression of Potra2n10c20525, which is annotated as dirigent protein; this gene class is involved in cell wall biosynthesis and growth as well as stress resistance (Paniagua *et al*., 2017). Overall, these relatively straightforward uses of the available data sets enable us to establish that the significant SNPs on chromosome 10 in ScotAsp are associated with lincRNAs and transcripts potentially involved in leaf development processes, that these vary in expression during leaf development, and that local SNPs are associated with the expression (eQTL) of developmental genes in SwAsp. While speculative, this demonstrates how *in silico* tools can be used to integrate evidence from a diverse range of genomics and population genetics data to develop hypotheses and to prioritise among candidate genes for downstream characterisation work.

**Figure 5.**
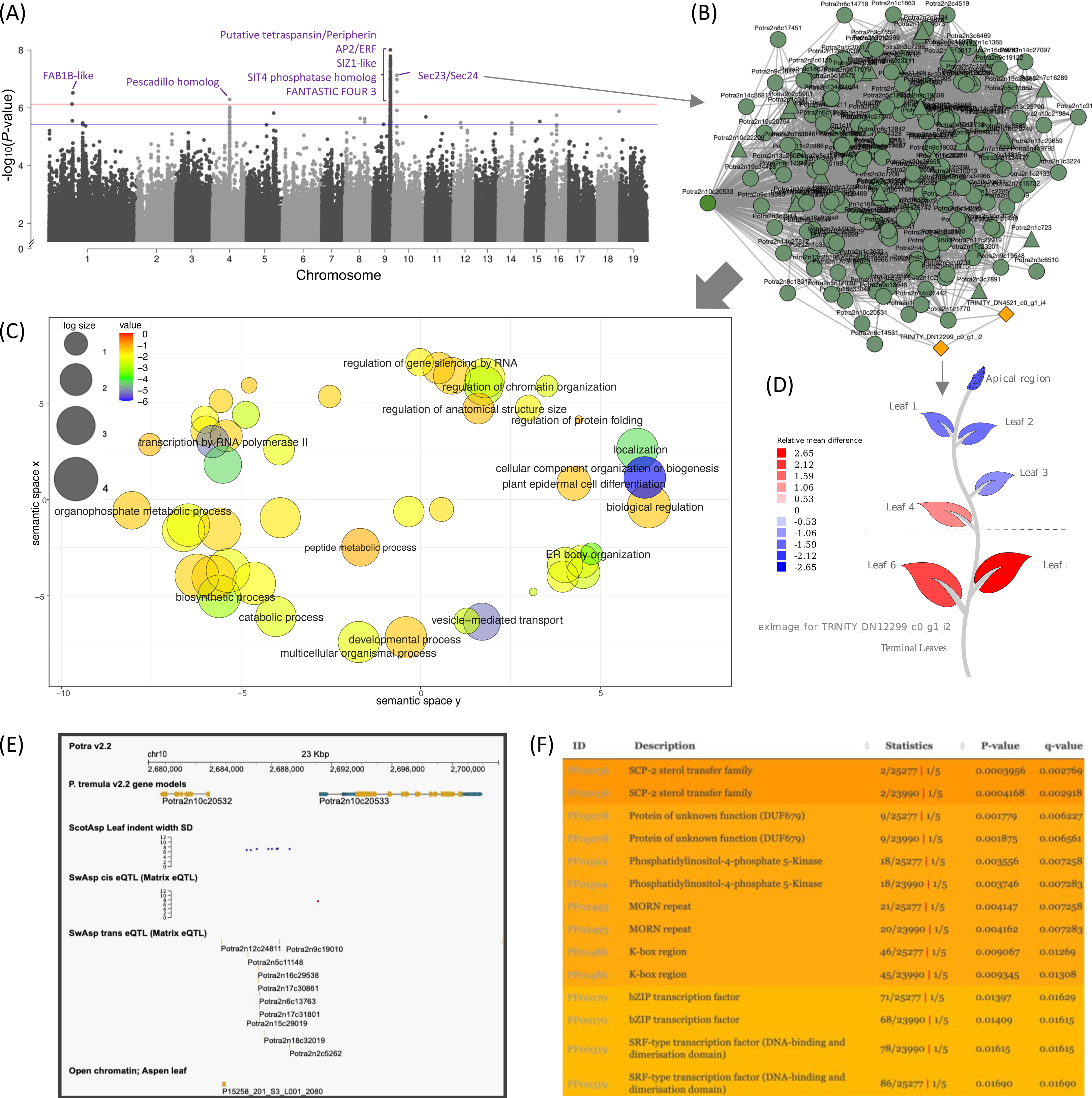
Genome-wide Single Nucleotide Polymorphism (SNP) associations for leaf shape in the Scottish Aspen collection (ScotAsp) and exploration of results in PlantGenIE. **(A)** Manhattan plot distribution of SNP associations for leaf indent width standard deviation (SD) in ScotAsp, where the red line indicates significance at *q*-value < 0.05 and the blue line is ‘suggestive significance’ at *q*-value < 0.1. Significant SNPs/groups of SNPs are annotated with the name of the associated Potra v2.2 gene; full details in Table 3. **(B)** A peak in the Manhattan plot indicates significant Single Nucleotide Polymorphisms from the Genome-Wide Association Study for leaf indent width SD on chromosome 10 comprises nine associated SNPs significant at *q*-value < 0.05, eight of which are located in an intergenic region between Potra2n10c20532 and Potra2n10c20533, and one located upstream of Potra2n10c20533, annotated as a Sec23/Sec24 transport family protein. The genes co-expressed with Potra2n10c20533 were examined in the exNet tool using the “Expand network” button to visualise first degree neighbours selected at *P*-value threshold 10^-1^ with genes shown as circles, transcription factors shown as triangles, and lincRNAs shown as yellow diamonds. **(C)** The resulting list of co-expressed 205 genes was tested using the Enrichment Tool to perform a gene ontology (GO) over-enrichment test and visualised using REVIGO. Circles representing the GO categories are scaled to the size of the term in the gene ontology database and coloured by enrichment -log_10_(*P*-value). **(D)** Co-expressed genes of Potra2n10c20533 were examined using the exImage tool, where it is possible to view the expression of the gene in the AspLeaf dataset of gene expression in terminal leaves; the example here is a lincRNA, TRINITY_DN12299_c0_g1_i2, with contrasting relative expression across the leaf development series. Shading of the exImage dataset is scaled as the relative mean difference between the greatest and least expression values. **(E)** Example of the use of JBrowse showing the region of chromosome 10 including Potra2n10c20532 and Potra2n10c20533, with tracks showing the co-location of significant GWAS results (*q-*value <0.05) for leaf indent width SD in ScotAsp, one *cis*-eQTL in Potra2n10c20533, ten *trans*-eQTL in the intergenic regions acting on ten individual genes, and an open chromatin region. **(F)** Use of the Enrichment tool showing Pfam enrichments for the set of ten genes acted on by *trans-*eQTL shown in (E).

### Conclusions

The improved genome assembly and population genetics data presented here, and from numerous existing studies, have been updated to the v2.2 genome and integrated into PlantGenIE (Sundell *et al*., 2015) to serve as a comprehensive community resource to facilitate hypothesis exploration and generation. To demonstrate the value and utility of the improved genome resource detailed here we performed GWAS for leaf physiognomy phenotypes in three aspen collections. We demonstrate use of the PlantGenIE.org resource to explore the Potra2n10c20533 gene that harbours a SNP associated to the standard deviation of leaf indent width. This is coupled with a complete phenotype data resource for the leaf physiognomy traits studied. The data presented is all publicly available, as summarised in Figure 6. Genomic resources in aspen have facilitated the characterisation of adaptive traits (Wang *et al*., 2018), omnigenic traits (Mähler *et al*., 2020), and the use of GWAS as a tool to guide candidate gene discovery (Grimberg *et al*., 2018) in addition to functional genomics insights into wood formation (Sundell *et al*., 2017) and sex determination (Muller *et al*., 2020), among others. Integration of these data in PlantGenIE.org enables rapid exploration of hypotheses, for example the potential functional role of candidate genes and can help in selecting among candidates for downstream studies to investigate and elucidate their functional and adaptive significance.

**Figure 6.**
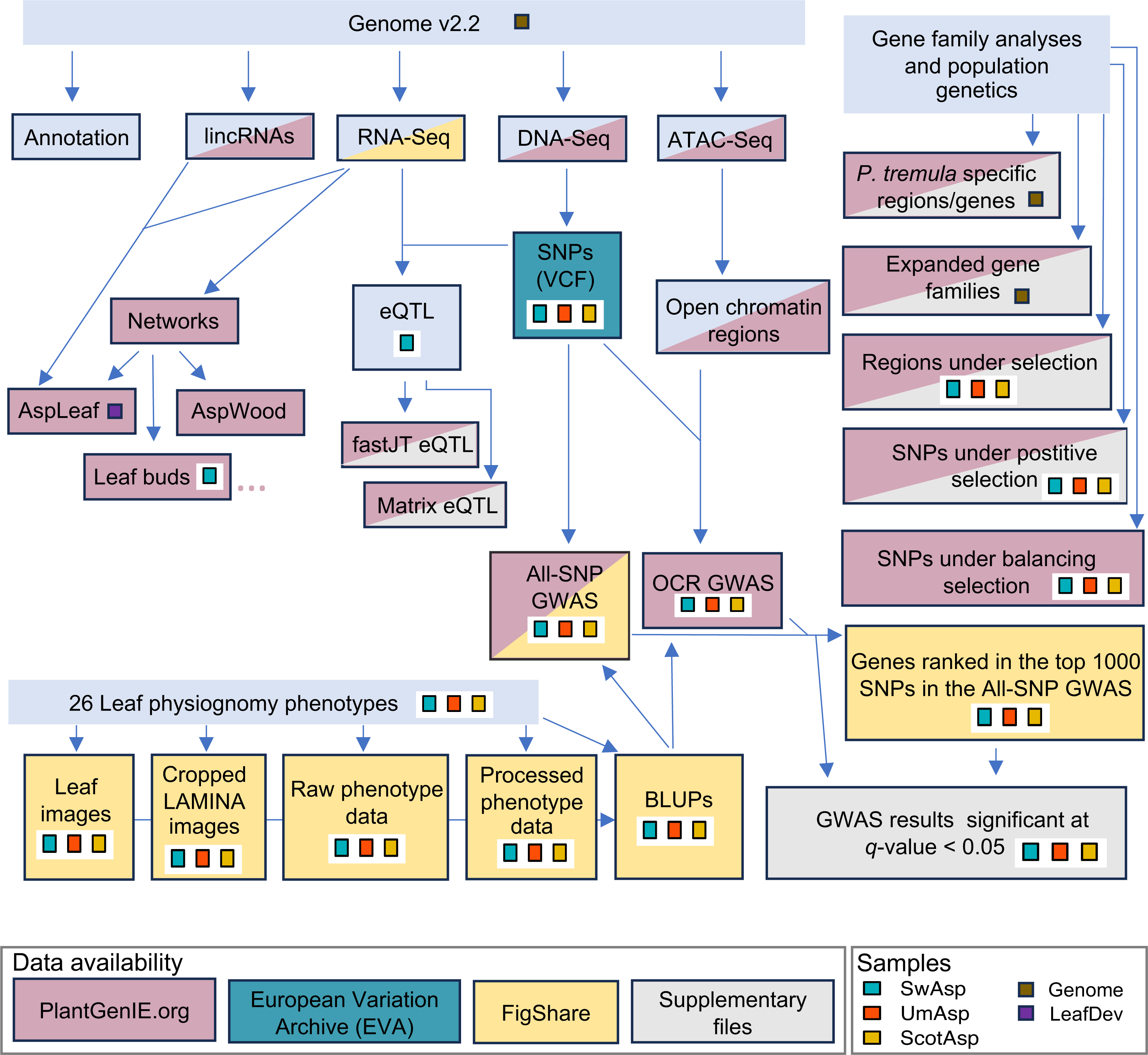
Overview of data accessibility for the genome and population genetics resource. The datasets that we present are grouped into three main sections: the Genome, Gene family analyses and population genetics, and Phenotype data. Boxes with each data set presented here are linked by arrows showing related data types and coloured by source of data accessibility: PlantGenIE.org = as a browsable tool / flat file available in at PlantGenIE.org; ENA = file available at the European Nucleotide Archive; FigShare = files available for download from FigShare at the SciLife Data Repository; Supplementary file = supplementary files available with this article at the publisher’s website. Samples from which the data files are derived are: SwAsp, the Swedish Aspen collection; UmAsp, the Umeå aspen collection; ScotAsp, the Scottish Aspen collection; Genome, the original tree that was sequenced for the genome assembly; LeafDev, the gene expression data set from developing aspen leaves described in Mähler *et al*. (2020).

## Author contributions

Conducted the experiments: KMR, BS, VK, KHR, SA, KC, NRS.

Analysed and/or interpreted the data: KMR, BS, HL, SMW, MR-A, TAK, VK, CC, CB, ND, TD, CM, JW, NM, SA, KC, NRS.

Provided materials and resources: JJ, VS, JC, KV, E-JP, SP, SJ, PKI, NRS.

Wrote the manuscript: KMR, BS and NRS with contributions from all authors. All authors read and approved the manuscript.

## Acknowledgements

Optical map production and analysis were performed by the VIB Nucleomics Core (www.nucleomics.be). The authors would like to acknowledge support from Science for Life Laboratory, the National Genomics Infrastructure, NGI, and Uppmax for providing assistance in massive parallel sequencing and computational infrastructure. The UmAsp sequences were funded by a grant from The Swedish Biodiversity Program at SciLife to Stefan Jansson. Computations were performed on resources provided by SNIC through Uppsala Multidisciplinary Centre for Advanced Computational Science (UPPMAX) under projects SNIC 2019/8-60, b2017115, b2010014, sllstore2017050, sllstore2017059, SNIC 2017/1-499, SNIC 2017/7-219, SNIC 2018/3-552, SNIC 2019/3-597, and SNIC 2021/5-62. We particularly thank Olga Pettersson at the Science for Life Laboratory for efforts, discussion and input. We thank Michiel Van Bel at the VIB_UGent Center for Plant Systems Biology for assistance with the PLAZA platform. We thank the UPSC bioinformatics platform (UPSCb) for computational infrastructure. The SwAsp and UmAsp common gardens were hosted and maintained by Skogforsk. We thank Zulema Carracedo Lorenzo for assistance in the field and laboratory. The ScotAsp clone garden was planted and maintained by Forest Research Technical Services Unit, with funding from the Forestry Commission. This work was supported by the Knut and Alice Wallenberg Foundation, VINNOVA UPSC Centre for Forest Biotechnology, the Research and Development Program for Forestry Technology (Project S111416L0710) provided by Korea Forest Service, and the Trees for the Future (T4F) project, The Swedish Research Council for Environment, Agricultural Science, and The Swedish Research Council to NRS. Hui Liu thanks the China Scholarship Council (CSC) for the financial support (No. 201906510022).

## Data availability and FAIR (Findable Accessible Interoperable Reusable) compliance

Data used to generate the genome assembly are available as European Nucleotide Archive (ENA; https://www.ebi.ac.uk/ena/browser/home) accession PRJEB41363, and the UmAsp re-sequencing data are available as accession PRJEB47451. The ATAC-Seq data are available as accession: in progress. The genome data are available to browse at https://plantgenie.org/. The VCF files for the UmAsp, SwAsp and ScotAsp collections are available at the European Variant Archive: in progress. Significant genome-wide association results (at *q*-value < 0.05), SNP variant files and regions under positive and balancing selection are available at https://plantgenie.org/JBrowse_new. Raw and processed phenotype files, top-ranked GWAS results and ATAC-Seq peaks are available at: the SciLife Data Repository Figshare: doi:10.17044/scilifelab.25335448.

## Supporting information files

**Appendix S1.** References to Table S2 of the plant genomes included in the gene family analysis.

**Appendix S2.** An overview of the metrics measured along the proximodistal and centrolateral leaf axis by default in the LAMINA software. Additional leaf physiognomy measurements to LAMINA defaults. All leaf metrics in the LAMINA analyses, phenotype data, trait names, types, and inclusion Genome-Wide Association study. Narrow-sense or ‘chip’-heritability estimates for leaf size and shape traits measured in the Umeå Aspen (UmAsp) collection.

**Supplementary table S1.** Plant genomes included in the gene family analysis.

**Supplementary table S2.** Details of samples from the UmAsp collection for DNA-Seq analysis and details of SwAsp and ScotAsp samples.

**Supplementary table S3.** Benchmarking Universal Single-Copy Orthologue (BUSCO) genome statistics and core Gene Family (coreGF) transcript statistics for *Populus tremula* assemblies v1.1 (Lin *et al*., 2018) and v2.2.

**Supplementary table S4**. **Genomic** regions (I), genes within those regions (II) and GO enrichment results (III) of *P. tremula* specific genes identified from synteny analysis.

**Supplementary table S5.** Structural rearrangements.

**Supplementary table S6.** Genes (I) GO enrichment (II) of *P. tremula* genes with Ka/Ks>1.

**Supplementary table S7.** Species- and clade-specific gene families.

**Supplementary table 8**. Genes (I) and GO enrichment results (II) of *P. tremula* specific genes identified from gene family analysis.

**Supplementary table S9**. Genes (I) and GO enrichment results (II) of *P. tremula* expanded gene families.

**Supplementary table S10**. Novel lincRNAs in aspen leaves.

**Supplementary table S11**. Climate data for the SwAsp, UmAsp and ScotAsp.

**Supplementary table S12**. Genomic regions under selection in SwAsp and UmAsp (I), genes within those regions (II) and GO (III) and Pfam (IV) enrichment of those genes.

**Supplementary table S13**. Significant genome-wide association results, comparison of ranks in the All-SNP GWAS for SNPs significant in the OCR GWAS, and details of linkage disequilibrium in the region of chromosome 9 in the All-SNP indent width SD GWAS in ScotAsp. SNPs and genes in linkage disequilibrium in this region.

**Supplementary table S14**. Significant eQTL at FDR < 0.05.

**Supplementary table S15**. Lists of the top-ranked genes from the All-SNP GWAS.

**Supplementary figure S1.** Ks distribution of *P. tremula* and *P. trichocarpa*.

**Supplementary figure S2.** Phylogenetic tree used to infer the analyse expansion and contraction of gene families.

**Supplementary figure S4.** JBrowse view for a 246 Kbp region of chromosome 9 with tracks displayed including Potra v2.2 gene models, Manhattan view of significant SNPs in the GWAS in All-SNP and OCR GWAS, *P. tremula*-specific regions, SNPs under balancing selection, and eQTL associations.

## Notes

https://github.com/bschiffthaler/aspen-v2/

### Competing Interest Statement

The authors have declared no competing interest.

### Summary of Updates

Complete re-work to include population genetics data and results

## Reference

Adebali O, Ortega DR, Zhulin IB. 2015. CDvist: a webserver for identification and visualization of conserved domains in protein sequences. Bioinformatics 31: 1475–1477. 10.1093/bioinformatics/btu836

Alpers DH, Appel SH, Tomkins GM. 1965. A spectrophotometric assay for thiogalactoside transacetylase. The Journal of Biological Chemistry 240: 10–13. 10.1016/S0021-9258(18)97606-4

Altschul SF, Gish W, Miller W, Myers EW, Lipman DJ. 1990. Basic local alignment search tool. Journal of Molecular Biology 215: 403–410. 10.1016/S0022-2836(05)80360-2

Amarasinghe SL, Su S, Dong X, Zappia L, Ritchie ME, Gouil Q. 2020. Opportunities and challenges in long-read sequencing data analysis. Genome Biology 21: 1–16. 10.1186/s13059-020-1935-5

An X, Gao K, Chen Z, et al. 2020. Hybrid origin of *Populus tomentosa* Carr. identified through genome sequencing and phylogenomic analysis. bioRxiv, 2020.04.07.030692. Available at: https://www.biorxiv.org/content/10.1101/2020.04.07.030692v1

Apuli R-P, Bernhardsson C, Schiffthaler B, Robinson KM, Jansson S, Street NR, Ingvarsson PK. 2020. Inferring the genomic landscape of recombination rate variation in European Aspen (*Populus tremula*). Genes*|Genomes|Genetics* 10: 299–309. 10.1534/g3.119.400504

Ariizumi T, Hatakeyama K, Hinata K, Inatsugi R, Nishida I, Sato S, Kato T, Tabata S, Toriyama K. 2004. Disruption of the novel plant protein NEF1 affects lipid accumulation in the plastids of the tapetum and exine formation of pollen, resulting in male sterility in Arabidopsis thaliana. The Plant Journal 39: 170–181. 10.1111/j.1365-313X.2004.02118.x

Bae, EK, Kang, MJ, Lee, SJ, Park EJ, Kim KT. 2023. Chromosome-level genome assembly of the Asian aspen *Populus davidiana* Dode. Sci Data 10: 431. 10.1038/s41597-023-02350-5

Bai S, Wu H, Zhang J, Pan Z, Zhao W, Li Z, Tong C. 2021. Genome Assembly of Salicaceae *Populus deltoides* (Eastern Cottonwood) I-69 Based on Nanopore Sequencing and Hi-C Technologies. Journal of Heredity 112: 303–310. 10.1093/jhered/esab010

Baute J, Herman D, Coppens F, De Block J, Slabbinck B, Dell’Acqua M, Pè ME, Maere S, Nelissen H, Inzé D. 2015. Correlation analysis of the transcriptome of growing leaves with mature leaf parameters in a maize RIL population. Genome Biol. 16: 168. doi: 10.1186/s13059-015-0735-9

Bååth R. 2016. bayesboot: an implementation of Rubin’s (1981) Bayesian Bootstrap. https://CRAN.R-project.org/package=bayesboot.

Ben-Gera H, Dafna A, Alvarez JP, Bar M, Mauerer M, Ori N. 2016. Auxin-mediated lamina growth in tomato leaves is restricted by two parallel mechanisms. Plant J. 86: 443–57. doi: 10.1111/tpj.13188

Bernhardsson C, Robinson KM, Abreu IN, Jansson S, Albrectsen BR, Ingvarsson PK. 2013. Geographic structure in metabolome and herbivore community co-occurs with genetic structure in plant defence genes. Ecology Letters 16: 791–798. 10.1111/ele.12114

Boeckler GA, Gershenzon J, Unsicker SB. 2011. Phenolic glycosides of the Salicaceae and their role as anti-herbivore defenses. Phytochemistry 72: 1497–1509. 10.1016/j.phytochem.2011.01.038

Bolger AM, Lohse M, Usadel B. 2014. Trimmomatic: A flexible trimmer for Illumina sequence data. Bioinformatics 30: 2114–2120. 10.1093/bioinformatics/btu170

Bresadola L, Caseys C, Castiglione S, Buerkle CA, Wegmann D, Lexer C. 2019. Admixture mapping in interspecific *Populus* hybrids identifies classes of genomic architectures for phytochemical, morphological and growth traits. New Phytologist 223: 2076–2089. 10.1111/nph.15930

Browning BL, Zhou Y, Browning SR. 2018. A one-penny imputed genome from next generation reference panels. Am. J. Hum. Genet. 103: 338–348. doi:10.1016/j.ajhg.2018.07.015

Bucchini F, Del Cortona A, Kreft L, Botzki A, Van Bel M, Vandepoele K. 2021.TRAPID 2.0: a web application for taxonomic and functional analysis of de novo transcriptomes. Nucleic Acids Research 49: e101. 10.1093/nar/gkab565

Buchfink B, Xie C, Huson DH. 2014. Fast and sensitive protein alignment using DIAMOND. Nature Methods 12: 59–60. 10.1038/s41592-021-01101-x

Bylesjö M, Segura V, Soolanayakanahally RY, Rae AM, Trygg J, Gustafsson P, Jansson S, Street NR. 2008. LAMINA: a tool for rapid quantification of leaf size and shape parameters. BMC Plant Biol. 81: 1–9. 10.1186/1471-2229-8-82

Campbell MS, Holt C, Moore B, Yandell M. 2014. Genome Annotation and Curation Using MAKER and MAKER-P. Current Protocols in Bioinformatics 2014: 4.11.1-4.11.39. 10.1002/0471250953.bi0411s48

Catala R, Ouyang J, Abreu IA, Hu Y, Seo H, Zhang X, Chua NH. 2007. The Arabidopsis E3 SUMO ligase SIZ1 regulates plant growth and drought responses. Plant Cell. 19: 2952–66. doi: 10.1105/tpc.106.049981

Chang S, Puryear J, Cairney J. 1993. A simple and efficient method for isolating RNA from pine trees. Plant Mol Biol Rep 11: 113–116. 10.1007/BF02670468

Chedgy RJ, Köllner TG, Constabel CP. 2015. Functional characterization of two acyltransferases from *Populus trichocarpa* capable of synthesizing benzyl benzoate and salicyl benzoate, potential intermediates in salicinoid phenolic glycoside biosynthesis. Phytochemistry 113: 149–159. 10.1016/j.phytochem.2014.10.018

Chen S, Yu Y, Wang X, Wang S, Zhang T, Zhou Y, He R, Meng N, Wang Y, Liu W, Liu Z, Liu J, Guo Q, Huang H, Sederoff RR, Wang G, Qu G, Chen S. 2023. Chromosome-level genome assembly of a triploid poplar *Populus alba* ‘Berolinensis’. Molecular Ecology Resources 23: 1092–1107. 10.1111/1755-0998.13770

Chen L, Zhu QH. 2022. The evolutionary landscape and expression pattern of plant lincRNAs. RNA Biology 19: 1190–1207. 10.1080/15476286.2022.2144609

Chin CS, Peluso P, Sedlazeck FJ, Nattestad M, Concepcion GT, Clum A, Dunn C, O’Malley R, Figueroa-Balderas R, Morales-Cruz A, et al. 2016. Phased diploid genome assembly with single-molecule real-time sequencing. Nature Methods 13: 1050–1054. 10.1038/nmeth.4035

Cho HK, Ahn CS, Lee HS, Kim JK, Pai HS. 2013. Pescadillo plays an essential role in plant cell growth and survival by modulating ribosome biogenesis. Plant J. 76: 393–405. doi: 10.1111/tpj.12302

Coscia M, Neffke FMH. 2017. Network backboning with noisy data. Proceedings - International Conference on Data Engineering, 425–436. 10.1109/ICDE.2017.100

Cromer L, Jolivet S, Singh DK, Berthier F, De Winne N, De Jaeger G, Komaki S, Prusicki MA, Schnittger A, Guérois R, Mercier R. 2019. Patronus is the elusive plant securin, preventing chromosome separation by antagonizing separase. Proc Natl Acad Sci USA 116: 16018–16027. doi: 10.1073/pnas.1906237116

Danecek P, Auton A, Abecasis G, Albers CA, Banks E, DePristo MA, Handsaker RE, Lunter G, Marth GT, Sherry ST, McVean G, Durbin R, 1000 Genomes Project Analysis Group. 2011. The variant call format and VCFtools. Bioinformatics 27: 2156–2158. 10.1093/bioinformatics/btr330

Danecek P, Bonfield JK, Liddle J, Marshall J, Ohan V, Pollard MO, Whitwham A, Keane T, McCarthy SA, Davies RM, Li H. 2021. Twelve years of SAMtools and BCFtools. GigaScience 10: giab008, 10.1093/gigascience/giab008

Davies C, Ellis CJ, Iason GR, Ennos RA. 2014. Genotypic variation in a foundation tree (*Populus tremula* L.) explains community structure of associated epiphytes. Biol. Lett. 10: 20140190. 10.1098/rsbl.2014.0190

De Bie T, Cristianini N, Demuth JP, Hahn MW. 2006. CAFE: a computational tool for the study of gene family evolution. Bioinformatics 22: 1269–71. 10.1093/bioinformatics/btl097

de Carvalho D, Ingvarsson PK, Joseph J, Suter L, Sedivy C, Macaya-Sanz D, Cottrell J, Heinze B, Schanzer I, Lexer C. 2010. Admixture facilitates adaptation from standing variation in the European aspen (*Populus tremula* L.), a widespread forest tree. Mol. Ecol. 19:1638–1650. 10.1111/j.1365-294X.2010.04595.x

Dobin A, Davis CA, Schlesinger F, Drenkow J, Zaleski C, Jha S, Batut P, Chaisson M, Gingeras TR. 2013. STAR: ultrafast universal RNA-seq aligner. Bioinformatics 29: 15–21. 10.1093/bioinformatics/bts635

Dóczi R, Hatzimasoura E, Bögre L. 2011. Mitogen-activated protein kinase activity and reporter gene assays in plants. In: Dissmeyer N, Schnittger A. (eds) Plant Kinases. Methods in Molecular Biology (Methods and Protocols), vol 779. Humana, Totowa, NJ. 10.1007/978-1-61779-264-9_5

Donaldson JR, Stevens MT, Barnhill HR, Lindroth RL. 2006. Age-related shifts in leaf chemistry of clonal aspen (Populus tremuloides). Journal of Chemical Ecology 32: 1415– 1429. 10.1007/s10886-006-9059-2

El-Gebali S, Mistry J, Bateman A, Eddy SR, Luciani A, Potter SC, Qureshi M, Richardson LJ, Salazar GA, Smart A, et al. 2018. The Pfam protein families database in 2019. Nucleic Acids Research 47: 427–432. 10.1093/nar/gkaa913

Emms DM, Kelly S. 2015. OrthoFinder: solving fundamental biases in whole genome comparisons dramatically improves orthogroup inference accuracy. Genome Biology 16. 10.1186/s13059-015-0721-2

Fox J, Weisberg S. 2019. An R Companion to Applied Regression, Third edition. Sage, Thousand Oaks CA. https://socialsciences.mcmaster.ca/jfox/Books/Companion/

Fracheboud Y, Luquez V, Björkén L, Sjödin A, Tuominen H, Jansson S. 2009. The control of autumn senescence in European aspen. Plant physiology 149: 1982–1991. 10.1104/pp.108.133249

Gachomo EW, Jimenez-Lopez JC, Baptiste LJ, Kotchoni SO. 2014. GIGANTUS1 (GTS1), a member of Transducin/WD40 protein superfamily, controls seed germination, growth and biomass accumulation through ribosome-biogenesis protein interactions in Arabidopsis thaliana. BMC Plant Biol. 14: 37. doi: 10.1186/1471-2229-14-37

Gasteiger E, Gattiker A, Hoogland C, Ivanyi I, Appel RD, Bairoch A. 2003. ExPASy: the proteomics server for in-depth protein knowledge and analysis. Nucleic Acids Research 31: 3784–3788. 10.1093/nar/gkg563

Gautier M, Klassmann A, Vitalis R. 2017. rehh 2.0: a reimplementation of the R package rehh to detect positive selection from haplotype structure. Molecular Ecology Resources 17: 78–90. 10.1111/1755-0998.12634

Gehlenborg N. 2014. UpSetR: A More Scalable Alternative to Venn and Euler Diagrams for Visualizing Intersecting Sets. R package version 1.4.0. https://CRAN.R-project.org/package=UpSetR.

Gel B, Serra E. 2017. karyoploteR: an R/Bioconductor package to plot customizable genomes displaying arbitrary data and 2 CIBERONC. Bioninformatics 33: 3088–3090. 10.1093/bioinformatics/btx346

Ghurye J, Pop M. 2019. Modern technologies and algorithms for scaffolding assembled genomes. PLoS Computational Biology 15: e1006994. 10.1371/journal.pcbi.1006994

Ghurye J, Rhie A, Walenz BP, Schmitt A, Selvaraj S, Pop M, Phillippy AM, Koren S. 2019. Integrating Hi-C links with assembly graphs for chromosome-scale assembly (I Ioshikhes, Ed.). PLOS Computational Biology 15: e1007273. 10.1371/journal.pcbi.1007273

Goel M, Sun H, Jiao W-B, Schneeberger K. 2019. SyRI: finding genomic rearrangements and local sequence differences from whole-genome assemblies. Genome biology 20: 277. 10.1186/s13059-019-1911-0

Götz S, García-Gómez JM, Terol J, Williams TD, Nagaraj SH, Nueda MJ, Robles M, Talón M, Dopazo J, Conesa A. 2008. High-throughput functional annotation and data mining with the Blast2GO suite. Nucleic acids research 36: 3420–35. 10.1093/nar/gkn176

Grabherr MG, Haas BJ, Yassour M, Levin JZ, Thompson DA, Amit I, Adiconis X, Fan L, Raychowdhury R, Zeng Q, Chen Z, Mauceli E, Hacohen N, Gnirke A, Rhind N, Di Palma F, Birren BW, Nusbaum C, Lindblad-Toh K, Friedman N, Regev A. 2011. Trinity: reconstructing a full-length transcriptome without a genome from RNA-Seq data. Nature Biotechnology 29: 644. 10.1038/NBT.1883

Grimberg Å, Lager I, Street NR, Robinson KM, Marttila S, Mähler N, Ingvarsson PK, Bhalerao RP. 2018. Storage lipid accumulation is controlled by photoperiodic signal acting via regulators of growth cessation and dormancy in hybrid aspen. New Phytologist 219: 619–630. 10.1111/nph.15197

Gruner K, Leissing F, Sinitski D, Thieron H, Axstmann C, Baumgarten K, Reinstädler A, Winkler P, Altmann M, Flatley A, Jaouannet M, Zienkiewicz K, Feussner I, Keller H, Coustau C, Falter-Braun P, Feederle R, Bernhagen J, Panstruga R. 2021. Chemokine-like MDL proteins modulate flowering time and innate immunity in plants. J Biol Chem. 296: 100611. doi: 10.1016/j.jbc.2021.100611

Guerriero G, Hausman JF, Ezcurra I. 2015. WD40-Repeat Proteins in Plant Cell Wall Formation: Current Evidence and Research Prospects. Front Plant Sci. 6: 1112. doi: 10.3389/fpls.2015.01112

Guo R, Hu Y, Aoi Y, Hira H, Ge C, Dai X, Kasahara H, Zhao Y. 2022. Local conjugation of auxin by the GH3 amido synthetases is required for normal development of roots and flowers in Arabidopsis. Biochem Biophys Res Commun. 589: 16–22. doi: 10.1016/j.bbrc.2021.11.109

Gurevich A, Saveliev V, Vyahhi N, Tesler G. 2013. Genome analysis QUAST: quality assessment tool for genome assemblies. Bioinformatics 29: 1072–1075. 10.1093/bioinformatics/btt086

Haas BJ, Papanicolaou A, Yassour M, Grabherr M, Blood PD, Bowden J, Couger MB, Eccles D, Li B, Lieber M, et al. 2013. De novo transcript sequence reconstruction from RNA-seq using the Trinity platform for reference generation and analysis. Nature Protocols 8: 1494–1512. 10.1038/nprot.2013.084

Harrison A. 2009. Aspen growth trials: showing the species’ potential in Scotland. The biodiversity and management of aspen woodlands (eds Cosgrove P, Amphlett A). Grantown-on-Spey, UK: The Cairngorms Local Biodiversity Action Plan. pp. 49–51.

Hendelman A, Buxdorf K, Stav R, Kravchik M, Arazi T. 2012. Inhibition of lamina outgrowth following Solanum lycopersicum AUXIN RESPONSE FACTOR 10 (SlARF10) derepression. Plant Mol Biol. 78: 561–76. doi: 10.1007/s11103-012-9883-4

Hirano T, Matsuzawa T, Takegawa K, Sato MH. 2011. Loss-of-function and gain-of-function mutations in FAB1A/B impair endomembrane homeostasis, conferring pleiotropic developmental abnormalities in Arabidopsis. Plant Physiol. 155: 797–807. doi: 10.1104/pp.110.167981.

Hoff KJ, Lange S, Lomsadze A, Borodovsky M, Stanke M. 2016. BRAKER1: Unsupervised RNA-Seq-Based Genome Annotation with GeneMark-ET and AUGUSTUS Bioinformatics 32: 767–769. 10.1093/bioinformatics/btv661

Hou Z, Wang Z, Ye Z, Du S, Liu S, Zhang J. 2018. Phylogeographic analyses of a widely distributed Populus davidiana: Further evidence for the existence of glacial refugia of cool-temperate deciduous trees in northern East Asia. Ecology and Evolution 8: 13014–13026. 10.1002/ece3.4755

Huang S, Kang M, Xu A. 2017. HaploMerger2: rebuilding both haploid sub-assemblies from high-heterozygosity diploid genome assembly. Bioinformatics 33: 2577–2579. 10.1093/bioinformatics/btx220

Jansson S, Douglas CJ. 2007. *Populus*: A Model System for Plant Biology. Annual Review of Plant Biology 58: 435–458. 10.1146/annurev.arplant.58.032806.103956

Jiao WB, Schneeberger K. 2017. The impact of third generation genomic technologies on plant genome assembly. Current Opinion in Plant Biology 36: 64–70. 10.1016/j.pbi.2017.02.002

Kanehisa M, Goto S. 2000. KEGG: Kyoto Encyclopedia of Genes and Genomes. Nucleic Acids Research 28: 27–30. 10.1093/nar/28.1.27

Katoh K, Standley DM. 2013. MAFFT multiple sequence alignment software version 7: Improvements in performance and usability. Molecular Biology and Evolution 30: 772–780. 10.1093/molbev/mst010

Kersten B, Faivre Rampant P, Mader M, Le Paslier M-C, Bounon R, Berard A, Vettori C, Schroeder H, Leplé J-C, Fladung M. 2016. Genome Sequences of *Populus tremula* Chloroplast and Mitochondrion: Implications for Holistic Poplar Breeding. PLoS ONE 11: e0147209. 10.1371/journal.pone.0147209. 10.1371/journal.pone.0147209

Kolde R. 2019. pheatmap: Pretty Heatmaps. R package version 1.0.12. https://CRAN.R-project.org/package=pheatmap.

Kopylova E, Noé L, Touzet H. 2012. SortMeRNA: fast and accurate filtering of ribosomal RNAs in metatranscriptomic data. Bioinformatics 28: 3211–7. 10.1093/bioinformatics/bts611

Korf I. 2004. Gene finding in novel genomes. BMC Bioinformatics 5: 59. 10.1186/1471-2105-5-59

Kruijer W. 2019. heritability: Marker-Based Estimation of Heritability Using Individual Plant or Plot Data. R package version 1.3. https://CRAN.R-project.org/package=heritability.

Kruijer W, Boer MP, Malosetti M, Flood PJ, Engel B, Kooke R, Keurentjes JJB, Van Eeuwijk FA. 2015. Marker-based estimation of heritability in immortal populations. Genetics 199: 379–398. 10.1534/genetics.114.167916

Kulasekaran S, Cerezo-Medina S, Harflett C, Lomax C, de Jong F, Rendour A, Ruvo G, Hanley SJ, Beale MH, Ward JL. 2021. A willow UDP-glycosyltransferase involved in salicinoid biosynthesis (J Rohwer, Ed.). Journal of Experimental Botany 72: 1634–1648. 10.1093/jxb/eraa562

Kang YJ, Yang DC, Kong L, Hou M, Meng YQ, Wei L, Gao G. 2017. CPC2: a fast and accurate coding potential calculator based on sequence intrinsic features. Nucleic Acids Research 45: W12–W16. 10.1093/NAR/GKX428

Lee J, Han CT, Hur Y. 2010. Overexpression of BrMORN, a novel ‘membrane occupation and recognition nexus’ motif protein gene from Chinese cabbage, promotes vegetative growth and seed production in Arabidopsis. Mol Cells. 29: 113–22. doi: 10.1007/s10059-010-0006-2

Li H. 2013. Aligning sequence reads, clone sequences and assembly contigs with BWA-MEM. arXiv:1303.3997v2 [q-bio.GN]

Li H. 2018. Minimap2: Pairwise alignment for nucleotide sequences. Bioinformatics 34: 3094–3100. 10.1093/bioinformatics/bty191

Li A, Zhang J, Zhou Z. 2014. PLEK: A tool for predicting long non-coding RNAs and messenger RNAs based on an improved k-mer scheme. BMC Bioinformatics 15: 1–10. 10.1186/1471-2105-15-311

Lin J, Sibley A, Shterev I, Nixon A, Innocenti F, Chan C, Owzar K. 2019. fastJT: An R package for robust and efficient feature selection for machine learning and genome-wide association studies. BMC Bioinformatics 20: 333. 10.1186/s12859-019-2869-3

Lin YC, Wang J, Delhomme N, Schiffthaler B, Sundström G, Zuccolo A, Nystedt B, Hvidsten TR, de la Torre A, Cossu RM, et al. 2018. Functional and evolutionary genomic inferences in Populus through genome and population sequencing of American and European aspen. Proceedings of the National Academy of Sciences of the United States of America 115: E10970–E10978. 10.1073/pnas.1801437115

Liu PP, Montgomery TA, Fahlgren N, Kasschau KD, Nonogaki H, Carrington JC. 2007. Repression of AUXIN RESPONSE FACTOR10 by microRNA160 is critical for seed germination and post-germination stages. Plant J. 52: 133–46. doi: 10.1111/j.1365-313X.2007.03218.x

Lomsadze A, Ter-Hovhannisyan V, Chernoff YO, Borodovsky M. 2005. Gene identification in novel eukaryotic genomes by self-training algorithm. Nucleic acids research 33: 6494–6506. 10.1093/nar/gki937

Love MI, Huber W, Anders S. 2014. Moderated estimation of fold change and dispersion for RNA-seq data with DESeq2. Genome Biology 15: 1–21. 10.1186/s13059-014-0550-8

Luquez V, Hall D, Albrectsen BR, Karlsson J, Ingvarsson P, Jansson S. 2008. Natural phenological variation in aspen (*Populus tremula*): The SwAsp collection. Tree Genetics and Genomes 4: 279–292. 10.1007/s11295-007-0108-y

Ma, J., Wan, D., Duan, B., Bai, X., Bai, Q., Chen, N. and Ma, T. (2019) Genome sequence and genetic transformation of a widely distributed and cultivated poplar. Plant Biotechnol. J., 17, 451–460. 10.1111/pbi.12989

Mähler N, Wang J, Terebieniec BK, Ingvarsson PK, Street NR, Hvidsten TR. 2017. Gene co-expression network connectivity is an important determinant of selective constraint. PLoS Genetics 13: e1006402. 10.1371/journal.pgen.1006402

Mähler N, Schiffthaler B, Robinson KM, Terebieniec BK, Vučak M, Mannapperuma C, Bailey MES, Jansson S, Hvidsten TR, Street NR. 2020. Leaf shape in *Populus tremula* is a complex, omnigenic trait. Ecol Evol 10:11922–11940. 10.1002/ece3.6691

Marçais G, Kingsford C. 2011. A fast, lock-free approach for efficient parallel counting of occurrences of k-mers. Bioinformatics 27: 764–770. 10.1093/bioinformatics/btr011

McKenna, A., Hanna, M., Banks, E., et al. 2010. The Genome Analysis Toolkit: A MapReduce framework for analyzing next-generation DNA sequencing data. Genome Res. 20: 1297–1303. 10.1101/gr.107524.110

Michael TP, VanBuren R. 2020. Building near-complete plant genomes. Current Opinion in Plant Biology 54: 26–33. 10.1016/j.pbi.2019.12.009

Michelson IH, Ingvarsson PK, Robinson KM, Edlund E, Eriksson ME, Nilsson O, Jansson S. 2018. Autumn senescence in aspen is not triggered by day length. Physiologia Plantarum 162: 123–134. 10.1111/ppl.12593

Miura K, Lee J, Miura T, Hasegawa PM. 2010. SIZ1 controls cell growth and plant development in Arabidopsis through salicylic acid. Plant Cell Physiol. 51:103–13. doi: 10.1093/pcp/pcp171

Mouriz A, López-González L, Jarillo JA, Piñeiro M. 2015. PHDs govern plant development. Plant Signal Behav. 10:e993253. doi: 10.4161/15592324.2014.993253

Müller NA, Kersten B, Leite Montalvão AP, Mähler N, Bernhardsson C, et al. 2020. A single gene underlies the dynamic evolution of poplar sex determination. Nat. Plants 6: 630–637. 10.1038/s41477-020-0672-9

Myking T, Bøhler F, Austrheim G, Solberg EJ. 2011. Life history strategies of aspen (Populus tremula L.) and browsing effects: A literature review. Forestry 84: 61–71. 10.1093/forestry/cpq044

Nakano RT, Matsushima R, Ueda H, Tamura K, Shimada T, Li L, Hayashi Y, Kondo M, Nishimura M, Hara-Nishimura I. 2009. GNOM-LIKE1/ERMO1 and SEC24a/ERMO2 are required for maintenance of endoplasmic reticulum morphology in *Arabidopsis thaliana*. Plant Cell 21: 3672–85. doi: 10.1105/tpc.109.068270

Ou S, Chen J, Jiang N. 2018. Assessing genome assembly quality using the LTR Assembly Index (LAI). Nucleic acids research 46: e126. 10.1093/nar/gky730

Pakull B, Kersten B, Lüneburg J, Fladung M. 2015. A simple PCR-based marker to determine sex in aspen. Plant Biol J 17: 256–261. 10.1111/plb.12217

Palos K, Yu L, Railey CE, Nelson Dittrich AC, Nelson ADL. 2023. Linking discoveries, mechanisms, and technologies to develop a clearer perspective on plant long noncoding RNAs. The Plant Cell 35: 1762–1786. 10.1093/plcell/koad027

Pan W, Jiang T, Lonardi S. 2019. OMGS: Optical Map-Based Genome Scaffolding. Journal of Computational Biology 27: cmb.2019.0310. 10.1089/cmb.2019.0310

Paniagua C, Bilkova A, Jackson P, Dabravolski S, Riber W, Didi V, Houser J, Gigli-Bisceglia N, Wimmerova M, Budínská E, Hamann T, Hejatko J. 2017. Dirigent proteins in plants: modulating cell wall metabolism during abiotic and biotic stress exposure. J Exp Bot. 68: 3287–3301. doi: 10.1093/jxb/erx141

Patro R, Duggal G, Love MI, Irizarry RA, Kingsford C. 2017. Salmon provides fast and bias-aware quantification of transcript expression. Nature Methods 14: 417–419. 10.1038/nmeth.4197

Patterson N, Price AL, Reich D. 2006. Population Structure and Eigenanalysis. PLoS Genet 2: e190. 10.1371/journal.pgen.0020190

Peiter E, Montanini B, Gobert A, Pedas P, Husted S, Maathuis FJ, Blaudez D, Chalot M, Sanders D. A secretory pathway-localized cation diffusion facilitator confers plant manganese tolerance. 2007. Proc Natl Acad Sci USA. 104: 8532–7. doi: 10.1073/pnas.0609507104

Peterson RA, Cavanaugh JE. 2020. Ordered quantile normalization: a semiparametric transformation built for the cross-validation era. Journal of Applied Statistics 47: 2312–2327. doi: 10.1080/02664763.2019.1630372.

Philippe RN, Bohlmann J. 2007. Poplar defense against insect herbivores. Canadian Journal of Botany 85: 1111–1126. 10.1139/B07-109

Pinosio S, Giacomello S, Faivre-Rampant P, Taylor G, Jorge V, Le Paslier MC, Zaina G, Bastien C, Cattonaro F, Marroni F, Morgante M. 2016. Characterization of the poplar pan-genome by genome-wide identification of structural variation. Molecular Biology and Evolution 33: 2706–2719. 10.1093/molbev/msw161

Purcell S, Neale B, Todd-Brown K, Thomas L, Ferreira MAR, Bender D, Maller J, Sklar P, De Bakker PIW, Daly MJ, et al. 2007. PLINK: A tool set for whole-genome association and population-based linkage analyses. American Journal of Human Genetics 81: 559–575. 10.1086/519795

Qu X, Chatty PR, Roeder AHK. 2014. Endomembrane Trafficking Protein SEC24A Regulates Cell Size Patterning in Arabidopsis. Plant Physiology 166: 1877–1890. 10.1104/pp.114.246033

Quinlan AR, Hall IM. 2010. BEDTools: a flexible suite of utilities for comparing genomic features. Bioinformatics 26: 841–842. 10.1093/BIOINFORMATICS/BTQ033

R Core Team. 2019. A Language and Environment for Statistical Computing. R Foundation for Statistical Computing, Vienna, Austria. https://www.R-project.org/

Reimann R, Kost B, Dettmer J. 2017. TETRASPANINs in Plants. Front Plant Sci. 18: 545. doi: 10.3389/fpls.2017.00545

Rendón-Anaya M, Wilson J, Sveinsson S, Cottrell J, Bailey ME, Ruņģis D, Lexer C, Jansson S, Robinson KM, Street NR, Ingvarsson PK. 2021. Adaptive Introgression Facilitates Adaptation to High Latitudes in European Aspen (*Populus tremula* L.). Molecular Biology and Evolution 38: 5034–5050. 10.1093/molbev/msab229

Robinson KM, Delhomme N, Mähler N, Schiffthaler B, Onskog J, Albrectsen BR, Ingvarsson PK, Hvidsten TR, Jansson S, Street NR. 2014. *Populus tremula* (European aspen) shows no evidence of sexual dimorphism. BMC plant biology 14: 276. 10.1186/s12870-014-0276-5

Robinson KM, Hauzy C, Loeuille N, Albrectsen BR. 2015. Relative impacts of environmental variation and evolutionary history on the nestedness and modularity of tree-herbivore networks. Ecology and Evolution 5: 2898– 2915. 10.1002/ece3.1559

Robinson KM, Ingvarsson PK, Jansson S, Albrectsen BR. 2012. Genetic variation in functional traits influences arthropod community composition in aspen (*Populus tremula* L.). PLoS ONE 7: 1–12. 10.1371/journal.pone.0037679

Rodgers-Melnick E, Vera DL, Bass HW, Buckler ES. 2016. Open chromatin reveals the functional maize genome. Proc Natl Acad Sci USA 113: E3177–84. doi: 10.1073/pnas.1525244113

Sanderson MJ. 2003. r8s: inferring absolute rates of molecular evolution and divergence times in the absence of a molecular clock. Bioinformatics 19: 301–2. 10.1093/bioinformatics/19.2.301

Shabalin AA. 2012. Matrix eQTL: ultra fast eQTL analysis via large matrix operations. Bioinformatics. 28: 1353–8. doi: 10.1093/bioinformatics/bts163

Schiffthaler B, Van Zalen E, Serrano AR, Street NR, Delhomme N. (2023). SeiÃ°r: Efficient calculation of robust ensemble gene networks. Heliyon, 9: e16811. 10.1016/j.heliyon.2023.e16811

Shevchenko A, Wilm M, Vorm O, Mann M.1996. Mass spectrometric sequencing of proteins from silver-stained polyacrylamide gels. Analytical Chemistry 68: 850–858. 10.1021/ac950914h

Shi TL, Jia KH, Bao YT, Nie S, Tian XC, Yan XM, Chen ZY, Li ZC, Zhao SW, Ma HY, Zhao Y, Li X, Zhang RG, Guo J, Zhao W, El-Kassaby YA, Müller N, Van de Peer Y, Wang XR, Street NR, Porth I, An X, Mao JF. 2024. High-quality genome assembly enables prediction of allele-specific gene expression in hybrid poplar. Plant Physiol. 27:kiae078. doi: 10.1093/plphys/kiae078

Siewert KM, Voight BF. 2017. Detecting Long-Term Balancing Selection Using Allele Frequency Correlation. Mol Biol Evol 2017, 34: 2996–3005. 10.1093/molbev/msx209

Simão FA, Waterhouse RM, Ioannidis P, Kriventseva E V., Zdobnov EM. 2015. BUSCO: user guide. Bioinformatics 31: 3210–3212. 10.1093/bioinformatics/btv351

Slaten ML, Chan YO, Shrestha V, Lipka AE, Angelovici R. 2020. HAPPI GWAS: Holistic Analysis with Pre- and Post Integration GWAS. Bioinformatics 36: 4655–4657. 10.1093/bioinformatics/btaa589

Slavov GT, Zhelev P. 2010. Salient Biological Features, Systematics, and Genetic Variation of Populus. In: Genetics and Genomics of Populus. Springer New York, 15–38.

Sobuja N, Nissinen K, Virjamo V, Salonen A, Sivadasan U, Randriamanana T, Ikonen V-P, Kilpeläinen K, Julkunen-Tiitto R, Nybakken L, Mehtätalo L, Peltola H. 2021.Accumulation of phenolics and growth of dioecious *Populus tremula* (L.) seedlings over three growing seasons under elevated temperature and UVB radiation. Plant Physiology and Biochemistry 165: 114–122. 10.1016/j.plaphy.2021.05.012

Soolanayakanahally R, Guy R, Street N, Robinson K, Silim S, Albrectsen B, Jansson S. 2015. Comparative physiology of allopatric *Populus* species: geographic clines in photosynthesis, height growth, and carbon isotope discrimination in common gardens. Frontiers in Plant Science 6: 528. 10.3389/fpls.2015.00528

Stanke M, Diekhans M, Baertsch R, Haussler D. 2008. Using native and syntenically mapped cDNA alignments to improve de novo gene finding. Bioinformatics 24: 637–44. 10.1093/bioinformatics/btn013

Storey JD, Bass AJ, Dabney A, Robinson D. 2021. qvalue: Q-value estimation for false discovery rate control. R package version 2.26.0, http://github.com/jdstorey/qvalue.

Street NR, Ingvarsson PK. 2011. Association genetics of complex traits in plants. New Phytologist 189:909–922. 10.1111/j.1469-8137.2010.03593.x

Sundell D, Mannapperuma C, Netotea S, Delhomme N, Lin Y-C, Sjödin A, Van de Peer Y, Jansson S, Hvidsten TR, Street NR. 2015. The Plant Genome Integrative Explorer Resource: PlantGenIE.org. New Phytologist 208: 1149–1156. 10.1111/nph.13557

Sundell D, Street NR, Kumar M, Mellerowicz EJ, Kucukoglu M, Johnsson C, Kumar V, Mannapperuma C, Delhomme N, Nilsson O, et al. 2017. Aspwood: High-spatial-resolution transcriptome profiles reveal uncharacterized modularity of wood formation in *Populus tremula*. Plant Cell 29: 1585–1604. 10.1105/tpc.17.00153

Supek F, Bošnjak M, Škunca N, Šmuc T. 2011. REVIGO summarizes and visualizes long lists of gene ontology terms. PLoS One. 6: e21800. doi: 10.1371/journal.pone.0021800

Taguchi G, Ubukata T, Nozue H, Kobayashi Y, Takahi M, Yamamoto H, Hayashida N. 2010. Malonylation is a key reaction in the metabolism of xenobiotic phenolic glucosides in Arabidopsis and tobacco. The Plant Journal 63: 1031–1041. 10.1111/j.1365-313X.2010.04298.x

Tajima F. 1989. Statistical method for testing the neutral mutation hypothesis by DNA polymorphism. Genetics 123: 585–595. 10.1093/genetics/123.3.585

Tang H, Zhang X, Miao C, Zhang J, Ming R, Schnable JC, Schnable PS, Lyons E, Lu J. 2015. ALLMAPS: Robust scaffold ordering based on multiple maps. Genome Biology 16: 3. 10.1186/s13059-014-0573-1

The UniProt Consortium. 2019. UniProt: a worldwide hub of protein knowledge The UniProt Consortium. Nucleic Acids Research 47: D506–D515. 10.1093/nar/gky1049

Tibbits JFG, McManus LJ, Spokevicius A V., Bossinger G. 2006. A rapid method for tissue collection and high-throughput isolation of genomic DNA from mature trees. Plant Molecular Biology Reporter 24: 81–91. 10.1007/BF02914048

Turner S. 2017. qqman: Q-Q and Manhattan Plots for GWAS Data. R package version 0.1.4. https://CRAN.R-project.org/package=qqman.

Tuskan GA, DiFazio S, Jansson S, Bohlmann J, Grigoriev I, Hellsten U, Putnam M, Ralph S, Rombauts S, Salamov A, et al. 2006. The genome of black cottonwood, Populus trichocarpa (Torr. & Gray). Science 313: 1596–1604. 10.1126/science.1128691

Tuskan GA, DiFazio S, Faivre-Rampant P, Gaudet M, Harfouche A, Jorge V, Labbé JL, Ranjan P, Sabatti M, Slavov G, Street N, Tschaplinski TJ, Yin T. 2012. The obscure events contributing to the evolution of an incipient sex chromosome in *Populus*: a retrospective working hypothesis. Tree Genetics & Genomes 8, 559–571. 10.1007/s11295-012-0495-6

Tylewicz S, Tsuji H, Miskolczi P, Petterle A, Azeez A, Jonsson K, Shimamoto K, Bhalerao RP. 2015. Dual role of tree florigen activation complex component FD in photoperiodic growth control and adaptive response pathways. Proceedings of the National Academy of Sciences 112: 3140–3145. 10.1073/pnas.1423440112

Van Bel M, Diels T, Vancaester E, Kreft L, Botzki A, Van de Peer Y, Coppens F, Vandepoele K. 2018. PLAZA 4.0: an integrative resource for functional, evolutionary and comparative plant genomics. Nucleic acids research 46: D1190–D1196. 10.1093/nar/gkx1002

Veeckman E, Ruttink T, Vandepeople K. 2016. Are we there yet? Reliably estimating the completeness of plant genome sequences. The Plant Cell 28: 1759–1768. 10.1105/tpc.16.00349

Walker BJ, Abeel T, Shea T, Priest M, Abouelliel A, Sakthikumar S, Cuomo CA, Zeng Q, Wortman J, Young SK, et al. 2014. Pilon: An integrated tool for comprehensive microbial variant detection and genome assembly improvement. PLoS ONE 9: e112963. 10.1371/journal.pone.0112963

Wang J, Ding J, Tan B, Robinson KM, Michelson IH, Johansson A, Nystedt B, Scofield DG, Nilsson O, Jansson S, et al. 2018. A major locus controls local adaptation and adaptive life history variation in a perennial plant. Genome Biology 19: 1–17. 10.1186/s13059-018-1444-y

Wang J, Street NR, Scofield DG, Ingvarsson PK. 2016a. Natural selection and recombination rate variation shape nucleotide polymorphism across the genomes of three related *Populus* species. Genetics 202: 1185–1200. 10.1534/genetics.115.183152

Wang J, Street NR, Scofield DG, Ingvarsson PK. 2016b. Variation in linked selection and recombination drive genomic divergence during allopatric speciation of European and American aspens. Molecular Biology and Evolution 33: 1754–1767. 10.1093/molbev/msw051

Wang F, Muto A, Van de Velde J, Neyt P, Himanen K, Vandepoele K, Van Lijsebettens M. 2015. Functional Analysis of the Arabidopsis TETRASPANIN Gene Family in Plant Growth and Development. Plant Physiol. 169: 2200–14. doi: 10.1104/pp.15.01310

Wang Y, Tang H, Debarry JD, Tan X, Li J, Wang X, Lee T, Jin H, Marler B, Guo H, et al. 2012. MCScanX: a toolkit for detection and evolutionary analysis of gene synteny and collinearity. Nucleic acids research 40: e49. 10.1093/nar/gkr1293

Watari M, Kato M, Blanc-Mathieu R, Tsuge T, Ogata H, Aoyama T. 2022. Functional Differentiation among the Arabidopsis Phosphatidylinositol 4-Phosphate 5-Kinase Genes PIP5K1, PIP5K2 and PIP5K3. Plant Cell Physiol. 63: 635–648. doi: 10.1093/pcp/pcac025

Wheeler DL, Barrett T, Benson DA, Bryant SH, Canese K, Chetvernin V, Church DM, DiCuccio M, Edgar R, Federhen S, et al. 2007. Database resources of the National Center for Biotechnology Information. Nucleic acids research 35: D5–D12. 10.1093/nar/gkl1031

Weir BS, Cockerham CC. 1984. Estimating F-statistics for the analysis of population structure. Evolution 38: 1358–1370. 10.1111/j.1558-5646.1984.tb05657.x

Weisman CM, Murray AW, Eddy SR. 2020. Many, but not all, lineage-specific genes can be explained by homology detection failure. PLoS Biol. 18: e3000862. 10.1371/journal.pbio.3000862

Wickham H, François R, Henry L, Müller K. 2020. dplyr: A Grammar of Data Manipulation. R package version 0.8.4. https://CRAN.R-project.org/package=dplyr.

Wright MN, Ziegler A. 2017. ranger: A Fast Implementation of Random Forests for High Dimensional Data in C++ and R. Journal of Statistical Software 77: 1–17. 10.18637/jss.v077.i01

Wu, H., Yao, D., Chen, Y., Yang, W., Zhao, W., Gao, H. and Tong, C. 2020. De Novo Genome assembly of *Populus simonii* further supports that *Populus simonii* and *Populus trichocarpa* belong to different sections. G3 Genes|Genomes|Genetics 10: 455–466. 10.1534/g3.119.400913

Xiao C, Guo H, Tang J, Li J, Yao X, Hu H. 2021. Expression Pattern and Functional Analyses of Arabidopsis Guard Cell-Enriched GDSL Lipases. Front Plant Sci. 12: 748543. doi: 10.3389/fpls.2021.748543

Yang L, Liu H, Zhao J, Pan Y, Cheng S, Lietzow CD, Wen C, Zhang X, Weng Y. 2018. LITTLELEAF (LL) encodes a WD40 repeat domain-containing protein associated with organ size variation in cucumber. Plant J. 95: 834–847 doi: 10.1111/tpj.13991

Yang, W., Wang, K., Zhang, J., Ma, J., Liu, J. and Ma, T. 2017. The draft genome sequence of a desert tree Populus pruinosa. Gigascience 6: 1–7. 10.1093/gigascience/gix075

Yang Z. 2007. PAML 4: Phylogenetic Analysis by Maximum Likelihood. Molecular Biology and Evolution 24: 1586–1591. 10.1093/molbev/msm088

Yates TB, Feng K, Zhang J, Singan V, Jawdy SS, Ranjan P, Abraham PE, Barry K, Lipzen A, Pan C, Schmutz J, Chen J-G, Tuskan GA, Muchero W. 2021. The Ancient Salicoid Genome Duplication Event: A Platform for Reconstruction of De Novo Gene Evolution in Populus trichocarpa. Genome Biology and Evolution 13: evab198. 10.1093/gbe/evab198

Zhang Y, Liu T, Meyer CA, Eeckhoute J, Johnson DS, Bernstein BE, Nusbaum C, Myers RM, Brown M, Li W. 2008. Model-based analysis of ChIP-Seq (MACS). Genome biology 9: 1–9.

Zhang, M., Zhang, Y., Scheuring, C. et al. 2012. Preparation of megabase-sized DNA from a variety of organisms using the nuclei method for advanced genomics research. Nat. Protoc. 7: 467–478

Zhang J, Yang Y, Zheng K, Xie M, Feng K, Jawdy SS, Gunter LE, Ranjan P, Singan VR, Engle N, et al. 2018. Genome-wide association studies and expression-based quantitative trait loci analyses reveal roles of HCT2 in caffeoylquinic acid biosynthesis and its regulation by defense-responsive transcription factors in *Populus*. New Phytologist 220: 502–516. 10.1111/nph.15297

Zhang B, Zhu W, Diao S, Wu X, Lu J, Ding CJ, Su X. 2019. The poplar pangenome provides insights into the evolutionary history of the genus. Commun Biol 2: 215. 10.1038/s42003-019-0474-7

Zhang C, Dong SS, Xu J-Y, He W-M, Yang TL. 2019. PopLDdecay: a fast and effective tool for linkage disequilibrium decay analysis based on variant call format files. Bioinformatics 35: 1786–1788. 10.1093/bioinformatics/bty875

Zhang Z, Chen Y, Zhang J, Ma X, Li Y, Li M, Wang D, Kang M, Wu H, Yang Y, et al. 2020. Improved genome assembly provides new insights into genome evolution in a desert poplar (Populus euphratica). Molecular Ecology Resources 20: 781– 794. 10.1111/1755-0998.13142

Zheng Z, Guo Y, Novák O, Chen W, Ljung K, Noel JP, Chory J. 2016. Local auxin metabolism regulates environment-induced hypocotyl elongation. Nat Plants. 21: 16025. doi: 10.1038/nplants.2016.25

Zhong R, Allen JD, Xiao G, Xie Y. 2014. Ensemble-Based Network Aggregation Improves the Accuracy of Gene Network Reconstruction. PLoS ONE 9: e106319. 10.1371/JOURNAL.PONE.0106319

Zhou R, Jenkins JW, Zeng Y, Shu S, Jang H, Harding SA, Williams M, Plott C, Barry KW, Koriabine M, Amirebrahimi M, Talag J, Rajasekar S, Grimwood J, Schmitz RJ, Dawe RK, Schmutz J, Tsai C-J. 2023. Haplotype-resolved genome assembly of Populus tremula × P. alba reveals aspen-specific megabase satellite DNA. Plant J. 10.1111/tpj.16454s

Zhou X, Carbonetto P, Stephens M. 2013. Polygenic Modeling with Bayesian Sparse Linear Mixed Models. PLoS Genetics 9: 1003264. 10.1371/journal.pgen.1003264

Zhou X, Stephens M. 2012. Genome-wide efficient mixed-model analysis for association studies. Nature genetics 44: 821–4. 10.1038/ng.2310

