## Supplementary material for "An Improved Chromosome-scale Genome Assembly and Population Genetics resource for *Populus tremula*": Appendix S1 References to Table S1.docx

**Appendix S1.** References to Table S1 of the plant genomes included in the gene family analysis.

**Chaw SM, Liu YC, Wu YW, Wang HY, Lin CYI, Wu CS, Ke HM, Chang LY, Hsu CY, Yang HT, et al. 2019.** Stout camphor tree genome fills gaps in understanding of flowering plant genome evolution. *Nature Plants* **5**: 63–73.

**DePamphilis CW, Palmer JD, Rounsley S, Sankoff D, Schuster SC, Ammiraju JSS, Barbazuk WB, Chamala S, Chanderbali AS, Determann R, et al. 2013.** The Amborella genome and the evolution of flowering plants. *Science* **342:** 1241089.

**Dohm JC, Minoche AE, Holtgräwe D, Capella-Gutiérrez S, Zakrzewski F, Tafer H, Rupp O, Sörensen TR, Stracke R, Reinhardt R, et al. 2014.** The genome of the recently domesticated crop plant sugar beet (Beta vulgaris). *Nature* **505:** 546–549.

**Edwards KD, Fernandez-Pozo N, Drake-Stowe K, Humphry M, Evans AD, Bombarely A, Allen F, Hurst R, White B, Kernodle SP, et al. 2017.** A reference genome for Nicotiana tabacum enables map-based cloning of homeologous loci implicated in nitrogen utilization efficiency. *BMC Genomics* **18:** 448.

**Filiault DL, Ballerini ES, Mandáková T, Aköz G, Derieg NJ, Schmutz J, Jenkins J, Grimwood J, Shu S, Hayes RD, et al. 2018.** The Aquilegia genome provides insight into adaptive radiation and reveals an extraordinarily polymorphic chromosome with a unique history. *eLife* **7:** e36426.

**Guan R, Zhao Y, Zhang H, Fan G, Liu X, Zhou W, Shi C, Wang J, Liu W, Liang X, et al. 2016.** Draft genome of the living fossil Ginkgo biloba. *GigaScience* **5:** 49.

**Hellsten U, Wright KM, Jenkins J, Shu S, Yuan Y, Wessler SR, Schmutz J, Willis JH, Rokhsar DS. 2013.** Fine-scale variation in meiotic recombination in Mimulus inferred from population shotgun sequencing. *Proceedings of the National Academy of Sciences of the United States of America* **110:** 19478–19482.

**Jaillon O, Aury JM, Noel B, Policriti A, Clepet C, Casagrande A, Choisne N, Aubourg S, Vitulo N, Jubin C, et al. 2007.** The grapevine genome sequence suggests ancestral hexaploidization in major angiosperm phyla. *Nature* **449:** 463–467.

**Kaul S, Koo HL, Jenkins J, Rizzo M, Rooney T, Tallon LJ, Feldblyum T, Nierman W, Benito MI. et al. 2000.** Analysis of the genome sequence of the flowering plant *Arabidopsis thaliana*. *Nature* **408,** 796–815.

**Li X, Kui L, Zhang J, Xie Y, Wang L, Yan Y, Wang N, Xu J, Li C, Wang W, et al. 2016.** Improved hybrid de novo genome assembly of domesticated apple (Malus x domestica). *GigaScience* **5:** 35.

**Lin X, et al. 2000.** Analysis of the genome sequence of the flowering plant Arabidopsis thaliana. *Nature* **408:** 796–815.

**Lin YC, Wang J, Delhomme N, Schiffthaler B, Sundström G, Zuccolo A, Nystedt B, Hvidsten TR, de la Torre A, Cossu RM, et al. 2018.** Functional and evolutionary genomic inferences in Populus through genome and population sequencing of American and European aspen. *Proceedings of the National Academy of Sciences of the United States of America* **115:** E10970–E10978.

**Myburg AA, Grattapaglia D, Tuskan GA, Hellsten U, Hayes RD, Grimwood J, Jenkins J, Lindquist E, Tice H, Bauer D, et al.** **2014.** The genome of Eucalyptus grandis. *Nature* **510:** 356–362.

**Olsen JL, Rouzé P, Verhelst B, Lin YC, Bayer T, Collen J, Dattolo E, De Paoli E, Dittami S, Maumus F, et al. 2016.** The genome of the seagrass Zostera marina reveals angiosperm adaptation to the sea. *Nature* **530:** 331–335.

**Neale DB, Wegrzyn JL, Stevens KA, Zimin A V., Puiu D, Crepeau MW, Cardeno C, Koriabine M, Holtz-Morris AE, Liechty JD, et al. 2014.** Decoding the massive genome of loblolly pine using haploid DNA and novel assembly strategies. *Genome Biology* **15:** R59.

**Neale DB, McGuire PE, Wheeler NC, Stevens KA, Crepeau MW, Cardeno C, Zimin A V., Puiu D, Pertea GM, Sezen UU, et al. 2017.** The Douglas-Fir genome sequence reveals specialization of the photosynthetic apparatus in Pinaceae. *G3: Genes, Genomes, Genetics* **7:** 3157–3167.

**Nystedt B, Street N, Wetterbom A, Zuccolo A, Lin Y-C, Scofield D, Vezzi F, Delhomme N, Giacomello S, Alexeyenko A, et al. 2013.** The Norway spruce genome sequence and conifer genome evolution. *Nature* **497:** 579–584.

**Rensing SA, Lang D, Zimmer AD, Terry A, Salamov A, Shapiro H, Nishiyama T, Perroud PF, Lindquist EA, Kamisugi Y, et al. 2008.** The Physcomitrella genome reveals evolutionary insights into the conquest of land by plants. *Science* **319:** 64–69.

**Salojärvi J, Smolander OP, Nieminen K, Rajaraman S, Safronov O, Safdari P, Lamminmäki A, Immanen J, Lan T, Tanskanen J, et al. 2017.** Genome sequencing and population genomic analyses provide insights into the adaptive landscape of silver birch. *Nature Genetics* **49:** 904–912.

**Shen Q, Zhang L, Liao Z, Wang S, Yan T, Shi P, Liu M, Fu X, Pan Q, Wang Y, et al. 2018.** The Genome of Artemisia annua Provides Insight into the Evolution of Asteraceae Family and Artemisinin Biosynthesis. *Molecular Plant* **11:** 776–788.

**Silva-Junior OB, Grattapaglia D, Novaes E, Collevatti RG. 2018.** Genome assembly of the Pink Ipê (Handroanthus impetiginosus, Bignoniaceae), a highly valued, ecologically keystone Neotropical timber forest tree. *GigaScience* **7:** 1–16.

**Stevens KA, Wegrzyn JL, Zimin A, Puiu D, Crepeau M, Cardeno C, Paul R, Gonzalez-Ibeas D, Koriabine M, Holtz-Morris AE, et al. 2016.** Sequence of the sugar pine megagenome. *Genetics* **204:** 1613–1626.

**Tang H, Krishnakumar V, Bidwell S, Rosen B, Chan A, Zhou S, Gentzbittel L, Childs KL, Yandell M, Gundlach H, et al. 2014.** An improved genome release (version Mt4.0) for the model legume Medicago truncatula. *BMC Genomics* **15**: 312.

**Tuskan GA, DiFazio S, Jansson S, Bohlmann J, Grigoriev I, Hellsten U, Putnam M, Ralph S, Rombauts S, Salamov A, et al. 2006.** The genome of black cottonwood, Populus trichocarpa (Torr. & Gray). *Science* **313:** 1596–1604.

**Wan T, Liu ZM, Li LF, Leitch AR, Leitch IJ, Lohaus R, Liu ZJ, Xin HP, Gong YB, Liu Y, et al. 2018.** A genome for gnetophytes and early evolution of seed plants. *Nature Plants* **4:** 82–89.

**Wei C, Yang H, Wang S, Zhao J, Liu C, Gao L, Xia E, Lu Y, Tai Y, She G, et al. 2018.** Draft genome sequence of Camellia sinensis var. sinensis provides insights into the evolution of the tea genome and tea quality. *Proceedings of the National Academy of Sciences of the United States of America* **115:** E4151–E4158.

**Xu C-Q, Liu H, Zhou S-S, Zhang D-X, Zhao W, Wang S, Chen F, Sun Y-Q, Nie S, Jia K-H, et al. 2019.** Genome sequence of Malania oleifera, a tree with great value for nervonic acid production. *GigaScience* **8:** 1-14.

**Zhang Z, Chen Y, Zhang J, Ma X, Li Y, Li M, Wang D, Kang M, Wu H, Yang Y, et al. 2020.** Improved genome assembly provides new insights into genome evolution in a desert poplar (Populus euphratica). *Molecular Ecology Resources* **20:** 781– 794.
