## Supplementary material for "An Improved Chromosome-scale Genome Assembly and Population Genetics resource for *Populus tremula*": Appendix S2 LAMINA traits.docx

**Appendix S1.**

**1. An overview of the metrics measured along the proximodistal and centrolateral leaf axis by default in the LAMINA software.**

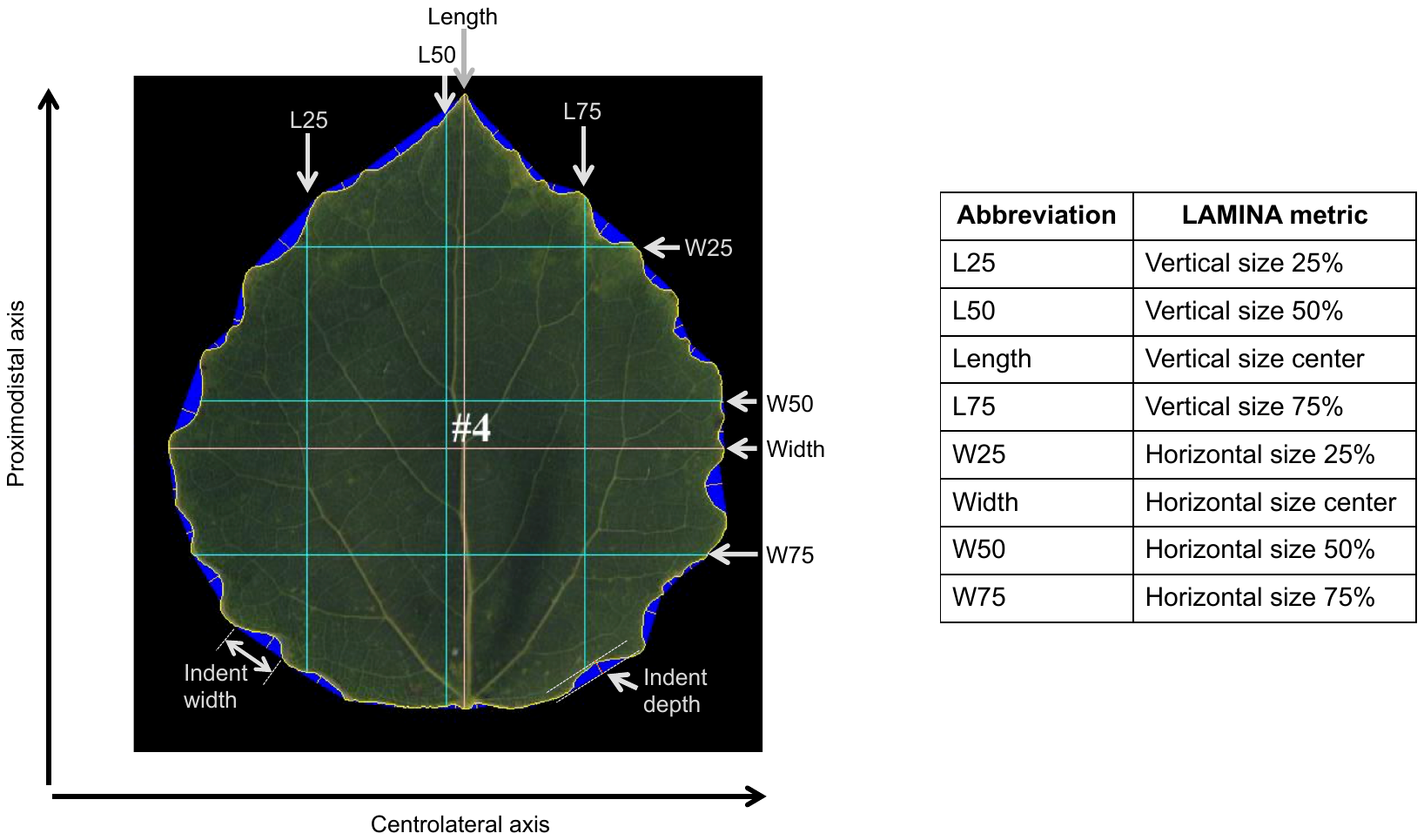

For more details of the measured parameters, see Bylesjö M, Segura V, Soolanayakanahally RY, Rae AM, Trygg J, Gustafsson P, Jansson S, Street NR. 2008. LAMINA: a tool for rapid quantification of leaf size and shape parameters. BMC Plant Biol. 81: 1–9. https://doi.org/10.1186/1471-2229-8-82

**2. Composite leaf physiognomy metrics in addition to the LAMINA defaults.**

Some of the default metrics from LAMINA were combined to calculate some composite metrics, with the aim of capturing additional variation in leaf shape in aspen.

| **Name** | **Description** | **Abbreviation in data files** | **Formula** |
| --- | --- | --- | --- |
| Indent density | Density of leaf margin serration | Indent.density | Perimeter 2 (excl. cavities)]/ Num. Indents |
| Indent width as a percentage of leaf perimeter | Width of leaf margin indent as percentage of the leaf perimeter | Ind.width.percent.perim | 100*[Indent width median / Perimeter 2 (excl. cavities)] |
| Indent depth as a percentage of leaf length | Median indent depth as percentage of length of central proximodistal leaf axis | Ind.depth.percent.leaf.length | 100*[Indent depth median/ Vert. size center] |
| Indent depth as a percentage of leaf width | Median indent depth as percentage of length of central centrolateral leaf axis | Ind.depth.percent.leaf.width | 100*[Indent depth median/ Horiz. size center] |
| Tip width: leaf length ratio | Ratio of leaf width at 25% along proximodistal leaf axis to length of proximodistal leaf axis | Horiz.25.vert.center (W25/Length) | Horiz. size 25%/Vert. size center |
| Apex angle | Ratio of leaf width at 25% along proximodistal leaf axis from tip, to width at centre proximodistal leaf axis | Horiz.25.horiz.center (W25/Width) | Horiz. size 25%/Horiz. size center |
| Basal angle | Ratio of leaf width at 75% along proximodistal leaf axis from tip, to width at centre proximodistal leaf axis | Horiz.75.horiz.center  (W75/Width) | Horiz. size 75%/Horiz. size center |

**3. All leaf metrics in the LAMINA analyses, phenotype data, trait names, types, and inclusion Genome-Wide Association study.**

| **Trait name in LAMINA analyses** | **Abbreviated trait name** | **Trait type** | **Metric** | **Trait included in GWAS** |
| --- | --- | --- | --- | --- |
| Area.2..cavities.filled. | Area | Size | LAMINA metric | Yes |
| Perimeter.2..excl..cavities. | Perimeter | Size | LAMINA metric | Yes |
| Squared.Perimeter.1.Area.1 |  | Shape | LAMINA metric | No |
| Squared.Perimeter.2.Area.2 | Squared Perimeter/Area | Shape | LAMINA metric | Yes |
| Circularity |  | Shape | LAMINA metric | Yes |
| Horiz..size.center | Width | Size | LAMINA metric | Yes |
| Horiz..size.25. | W25 | Size | LAMINA metric | Yes |
| Horiz..size.50. | W50 | Size | LAMINA metric | Yes |
| Horiz..size.75. | W75 | Size | LAMINA metric | Yes |
| Vert..size.center | Length | Size | LAMINA metric | Yes |
| Vert..size.25. | L25 | Size | LAMINA metric | Yes |
| Vert..size.50. | L50 | Size | LAMINA metric | Yes |
| Vert..size.75. | L75 | Size | LAMINA metric | Yes |
| Vert..size.center...Horiz..size.center | Length/Width | Shape | LAMINA metric | Yes |
| Horiz..size.25....Horiz..size.75. | H25/H75 | Shape | LAMINA metric | Yes |
| Vert..size.25....Vert..size.75. |  | Shape | LAMINA metric | No |
| Num..Indents | No. Indents | Shape | LAMINA metric | Yes |
| Indent.width.mean |  | Indent/Shape | LAMINA metric | No |
| Indent.width.median | Indent width | Indent/Shape | LAMINA metric | Yes |
| Indent.width.SD | Indent width SD | Indent/Shape | LAMINA metric | Yes |
| Indent.depth.mean |  | Indent/Shape | LAMINA metric | No |
| Indent.depth.median | Indent depth | Indent/Shape | LAMINA metric | Yes |
| Indent.depth.SD |  | Indent/Shape | LAMINA metric | Yes |
| Indent.density |  | Indent/Shape | Composite metric | Yes |
| Ind.width.percent.Perim |  | Shape | Composite metric | Yes |
| Ind.depth.percent.Leaflength |  | Shape | Composite metric | Yes |
| Ind.depth.percent.Leafwidth |  | Shape | Composite metric | Yes |
| Horiz.25.Vert.centre | W25/Length | Shape | Composite metric | Yes |
| Horiz.25.Horiz.centre | W25/Width | Shape | Composite metric | Yes |
| Horiz.75.Horiz.centre | W75/Width | Shape | Composite metric | Yes |

**4. Narrow-sense or Chip-heritability of leaf size and shape traits measured in the Umeå Aspen (UmAsp) collection.**

| **Collection** | **Type** | **Trait** | **h2** | **Lower 95 % C.I.** | **Upper 95 % C.I.** |
| --- | --- | --- | --- | --- | --- |
| UmAsp | Size | Area.2..cavities.filled. | 0.31 | 0.21 | 0.41 |
| UmAsp | Size | Horiz..size.25. | 0.48 | 0.39 | 0.57 |
| UmAsp | Size | Horiz..size.50. | 0.39 | 0.29 | 0.48 |
| UmAsp | Size | Horiz..size.75. | 0.43 | 0.34 | 0.52 |
| UmAsp | Size | Horiz..size.center | 0.38 | 0.28 | 0.48 |
| UmAsp | Size | Indent.depth.median | 0.65 | 0.58 | 0.72 |
| UmAsp | Size | Indent.width.median | 0.63 | 0.55 | 0.70 |
| UmAsp | Size | Perimeter.2..excl..cavities. | 0.37 | 0.27 | 0.46 |
| UmAsp | Size | Vert..size.25. | 0.31 | 0.21 | 0.41 |
| UmAsp | Size | Vert..size.50. | 0.31 | 0.21 | 0.41 |
| UmAsp | Size | Vert..size.75. | 0.30 | 0.20 | 0.40 |
| UmAsp | Size | Vert..size.center | 0.33 | 0.23 | 0.43 |
| UmAsp | Shape | Circularity | 0.66 | 0.60 | 0.73 |
| UmAsp | Shape | Horiz..size.25....Horiz..size.75. | 0.96 | 0.95 | 0.97 |
| UmAsp | Shape | Horiz.25.Horiz.centre | 0.86 | 0.83 | 0.89 |
| UmAsp | Shape | Horiz.25.Vert.centre | 0.78 | 0.73 | 0.83 |
| UmAsp | Shape | Horiz.75.Horiz.centre | 0.81 | 0.78 | 0.85 |
| UmAsp | Shape | Ind.depth.percent.Leaflength | 0.79 | 0.75 | 0.84 |
| UmAsp | Shape | Ind.depth.percent.Leafwidth | 0.79 | 0.74 | 0.84 |
| UmAsp | Shape | Ind.width.percent.Perim | 0.88 | 0.86 | 0.91 |
| UmAsp | Shape | Indent.density | 0.14 | 0.04 | 0.24 |
| UmAsp | Shape | Indent.depth.SD | 0.95 | 0.94 | 0.96 |
| UmAsp | Shape | Indent.width.SD | 0.62 | 0.55 | 0.70 |
| UmAsp | Shape | Num..Indents | 0.74 | 0.68 | 0.79 |
| UmAsp | Shape | Squared.Perimeter.2.Area.2 | 0.85 | 0.81 | 0.88 |
| UmAsp | Shape | Vert..size.center...Horiz..size.center | 0.97 | 0.97 | 0.98 |
