## Supplementary material for "An Improved Chromosome-scale Genome Assembly and Population Genetics resource for *Populus tremula*": Comprehensive materials and Methods.docx

**DNA extraction**

DNA for sequence data generation was extracted from fresh, young leaves of greenhouse-grown clonal replicates of the individual used to generate the v1.1 assembly presented in (Lin *et al.*, 2018). DNA was extracted using a CTAB-buffer based protocol based on Tibbits *et al.* (2006).

**Sequence Data**

For genome assembly and correction, we generated two libraries: “PacBio data”: 28,874,072,954 bases (filtered subreads, ~6̃0x coverage), Pacific Biosciences on the RSII platform (sequencing performed by Science for Life Laboratory, Uppsala, Sweden); and “Illumina data”: 108,353,739,802 bases (2̃26x coverage), Illumina HiSeq2500. Both datasets are available as ENA accession PRJEB41363. We also utilised six existing RNA-Seq datasets for use in the genome annotation: “AspWood” (Sundell *et al.*, 2017, ENA: ERP016242); “Sex” (Robinson *et al.*, 2014, ENA: ERP002471); “SwAsp” (Mähler *et al.*, 2017, ENA: ERP014886); “Assembly version 1 tissue atlas” (Lin *et al.*, 2018, ENA: PRJEB23585); “Xylem/Leaf” (Lin *et al.*, 2018b); and “Leaf Development” (Mähler *et al*., 2020, sequenced by BGI, China). Unless otherwise specified the Science for Life Laboratory in Stockholm generated all sequence data using standard Illumina TruSeq kits. The genome sequence and feature files are available from the PlantGenIE.org resource (Sundell *et al.,* 2015).

**Plant material, nuclei purification and ATAC-library**

To purify nuclei for the Assay of Transposase Accessible Chromatin sequencing (ATAC-Seq), 2 g of snap-frozen leaves from greenhouse grown *Populus tremula* were harvested and grounded in liquid nitrogen using a mortar and pestle. The nuclei were purified following method described in Zhang *et al.* (2012) with certain modifications. Briefly, the resulting powder was suspended ice cold 20 mL 1xNIB

(Nuclear Isolation Buffer containing [10 mM Trizma base, 0.5 M sucrose, 80 mM KCl, 10 mM EDTA, 1 mM spermidine trihydrochloride, 1 mM spermine tetrahydrochloride, 0.5% Triton X-100, 0.15% (vol/vol) 2-mercaptoethanol and protease inhibitor × 1 (Sigma P2714)] and incubated with gentle shaking at 4 °C for 15 min. The samples were then filtered through 4-layers of cheese cloth followed by 2-layers muslin cloth under gravity. The filtrate was centrifuged for 20 min at 3500 g and 4 °C. The pellet was washed 3-5 times after gentle resuspension in 10 mL 1xNIB. The resulting nuclei were counted using a BLAUBRAND Neubauer-improved chamber (depth of 0.1 mm, Sigma BR717810), after DAPI-staining under Zeiss Axioplan (fluorescence) microscopes.

Forty thousand nuclei were used for tagmentation and ATAC-library preparation at the Science for Life Laboratory (SciLife lab, Stockholm) using a standard Illumina Tagment DNA Enzyme and Buffer, and a Nextera DNA Library Prep Kit, respectively.

**Assembly**

All code and configuration related to the assembly process can be found at the associated GitHub repository: <https://github.com/bschiffthaler/aspen-v2>. Unless otherwise specified all software was run using default settings for the provided version.

Initially, we assembled the genome using FALCON v0.3 (Chin *et al.*, 2016). We include the FALCON config file in the Git repository. Subsequently, we aligned all the Illumina data to the initial assembly and used in-house scripts (available upon request) to correct homozygous SNPs and small indel (insertion/deletion) issues. Briefly, 100x coverage of Illumina short read data were aligned, variants were detected using GATK (version 3.4-46-gbc02625, McKenna *et al.*, 2010) and filtered to select homozygous variants only. We then selected high confidence alignments and replaced the reference sequence with the aligned consensus. We then aligned 100x of the Illumina data to the fixed assembly and repeated the aforementioned steps to correct homozygous SNPs and INDELs a second time. For a third and final round of fixing, we used the Illumina data as input to Pilon (v2.11-1.18, Walker *et al*., 2014) to correct assembly issues per contig. In order to reduce the presence of split haplotypes, we used HaploMerger2 (retrieved: 2015-11-06, Huang *et al*., 2017). We include all HaploMerger2 scripts in the Git repository. We subsequently created an optical map and hybrid assembly of the genome using the BioNano Irys system and software (produced by Nucleomics Core, Leuven, Belgium), which initially scaffolded and oriented the assembly. The map was constructed using fresh, young leaf material from clonal copies of the same individual used for genome assembly. For optical map assembly only molecules >150 kb and containing at least five labels and at most 0.6 average intensity were used. The optical map assembly was produced using the options ‘haplotype-OptArgs’. Subsequently, a hybrid-scaffold assembly was performed using the optical map and sequence assembly using the option ‘resolve conflicts’. This hybrid assembly and all unplaced contigs from the sequence-based assembly were then combined with the high-density genetic linkage map from Apuli *et al.* (2020) as input to ALLMAPS (v1.0.1, Tang *et al.*, 2015) to place the scaffolds into pseudo-chromosomes.

**Chloroplast and mitochondrion**

To flag sequences originating from the chloroplast, we matched all unplaced scaffolds to published chloroplast sequences (Kersten *et al.,* 2016), using blast+ (v2.9.0, e-value cutoff 10e-1). All scaffolds with greater than 85% cumulative coverage were flagged as “putative chloroplast”. The same was performed for the mitochondrion, however no assembly contigs were identified using this approach.

**Transcriptome Assembly**

We used five RNA-Seq datasets from *P. tremula* as supporting evidence for gene annotation. Four of the datasets had already been used for the annotation of the previous genome version (Lin *et al.*, 2018): exAtlas, exDiversity, Xylem/Leaf and Leaf Development, while the fifth was derived from our AspWood resource (Sundell *et al.*, 2017). These five datasets are available from the ENA (<https://ebi.ac.uk/ena>) under the accessions PRJEB5040, PRJEB1790, PRJEB28867, PRJEB28866, PRJEB14593, respectively. The reads were pre-processed as described in Lin *et al.* (2018) and Sundell *et al.* (2017). Briefly, the raw reads were filtered for rRNA using SortMeRNA (Kopylova *et al.*, 2012) version 2.1 and trimmed for adapter sequences and lower quality using Trimmomatic (Bolger *et al.*, 2014) version 0.39. The filtered reads were then assembled using Trinity (Haas *et al.*, 2013) version 2.8.4 using default settings. The resulting transcript fasta files were then used as expressed sequence tag (EST) evidence for Maker-P.

**Annotation**

We first collected a set of diverse RNA-Seq datasets from previous studies. These we aligned to the genome using STAR in 2-pass mode. For the first pass, we used the following arguments:

STAR --outFilterType BySJout --outFilterMultimapNmax 20 --alignSJoverhangMin 8 --alignSJDBoverhangMin 1 --outFilterMismatchNmax 999 --outFilterMismatchNoverReadLmax 0.1 --alignIntronMin 20 --alignIntronMax 20000 --alignMatesGapMax 5000 --outSAMtype BAM SortedByCoordinate --chimOutType WithinBAM

For the second pass, we provided the splice junctions from the first pass as pass-1-SJ.out.tab and used:

STAR --outFilterType BySJout --outFilterMultimapNmax 20 --alignSJoverhangMin 8 --alignSJDBoverhangMin 1 –outFilterMismatchNmax 999 --outFilterMismatchNoverReadLmax 0.1 --alignIntronMin 20 --alignIntronMax 20000 --alignMatesGapMax 5000 --outSAMtype BAM SortedByCoordinate --chimOutType WithinBAM --limitSjdbInsertNsj 2000000 --sjdbFileChrStartEnd pass-1-SJ.out.tab

We then provided these alignments to BRAKER1 (Hoff, 2016) along with protein sequences of version 1 of the assembly, running BRAKER1 in hybrid mode with arguments:
braker.pl --genome=genome.fa --prot_seq=protein.fa --prg=gth –softmasking --AUGUSTUS_ab_initio

In order to prepare the genome for annotation, we created a custom *de novo* repeat library using the genome sequence with RepeatModeler v1.0.11. We then concatenated the custom repeats with known repeats in *Viridiplantae* (RepBase database retrieved 2018-10-26, https://pubmed.ncbi.nlm.nih.gov/26045719/) and the first assembly of the *P. tremula* genome. We masked the genome using RepeatMasker4.0.8. (http://www.repeatmasker.org). We ran MAKER v2.31.10 (Campbell *et al.*, 2014) on the masked genome in three passes. We include the MAKER config files in the Git repository. We used Trinity assemblies from all RNA-Seq datasets in conjunction with all transcripts from the v1 assembly as expressed sequence tag (EST) evidence. Furthermore, we provided proteins from the v1 assembly and the v3.0 assembly of *P. trichocarpa* (Tuskan *et al.*, 2006) as protein evidence. In order to train AUGUSTUS v3.0.2 (Stanke *et al.*, 2008) and SNAP v2013-11-29 (Korf, 2004) we extracted confident predictions from the first run of MAKER using maker2zff from the MAKER suite and zff2augustus_gbk.pl from an external source (https://github.com/hyphaltip/genome-scripts/blob/master/gene_prediction/zff2augustus_gbk.pl). We then proceeded with another round of MAKER including AUGUSTUS, SNAP and GeneMark-ES (Lomsadze *et al.*, 2005). We repeated this process of training AUGUSTUS and SNAP once more for a third and final round of MAKER. MAKER config files at the Git repository contain exact settings used.

**Lift-over Alignments and Structural Variant Detection**

We performed the lift-over alignments of v1.1 scaffolds to the v2.2 assembly using minimap 2 (v2.2.2, Li, 2018).

We aligned the PacBio data to both the v1.1 and v2.2 assemblies using pbmm2 (v1.1.0), which uses minimap2 internally. We then performed variant detection using pbsv (v2.2.2, <https://github.com/PacificBiosciences/pbsv>).

**Functional Annotation**

We aligned the transcripts and protein-coding sequences retrieved from MAKER to the NCBI nt (Wheeler *et al.*, 2007) and UniRef90 (The UniProt Consortium, 2019) databases. For transcripts, we used Blast+ version 2.6.0+ (Altschul *et al.*, 1990) with the non-default parameters: -evalue 1e^-5^. For proteins, we used Diamond version 0.9.26 (Buchfink *et al.*, 2014). We identified and extracted the sequences aligning solely to the NCBI nt database to complement the UniRef90 alignments using an ad-hoc script (available upon request). We then imported the resulting alignment files into Blast2GO (Götz *et al.*, 2008) version 5.2. Finally, we used Blast2GO to generate Gene Ontology (both GO and GO-Slim), Pfam (El-Gebali *et al.*, 2018) and KEGG (Kanehisa & Goto, 2000) annotations.

**Genome assembly evaluation**

To calculate summary statistics of the assembly, we used QUAST v5.0.2 (Gurevich *et al.*, 2013), aligning a 20X coverage subset (generated by truncating the library to a total count of 8 * 10e^9^ nucleotides) of the aspen V1 2x150 PE library data (ENA: PRJEB23581) to calculate mapping percentages. We ran BUSCO v3.0.2 for both the genomic and transcript sequences. We retrieved the “embryophyta_odb10” dataset from <https://busco.ezlab.org/>. coreGF completeness scores were computed with TRAPID2.0 (Bucchini *et al.,* 2021), using Phylogenetic clade eudicotyledons [71240] and default settings (conservation threshold:0.9, Top hits:1).

**ATAC-Seq peak calling**

Raw reads were trimmed from adapter contamination, low quality bases and short fragments using Trimmomatic v.039 (Bolger *et al.*, 2014) (ILLUMINACLIP:$TRIMMOMATIC_HOME/adapters/NexteraPE-PE.fa:2:30:10:1:TRUE SLIDINGWINDOW:5:20 MINLEN:38). BWA-MEM v0.7.17 (Li, 2013) was then run with default parameters for aligning clean reads to the *Populus tremula* reference genome (Schiffthaler *et al.*, 2019) to which mitochondrial and chloroplast reference genomes (Kersten *et al.*, 2016) had been added for later filtering. Reads were then sorted, indexed and PCR duplicates were marked using SAMtools v1.9 (Danecek *et al*., 2021) followed by the removal of reads flagged as unmapped, not in proper pairs, marked as duplicate or with a MAPQ < 30 (-f 3 -F 12 -F 512 -F 1024 -q 30) or aligning to chloroplast or mitochondrial (grep -v -e 'KP861984.1' -e 'KT337313.1'). Peaks were called using MACS2 v2.2.7.1 (Zhang *et al.*, 2008) (-g 381973357 --nomodel --keep-dup all --format BAMPE).

**Transcriptome sequencing and data pre-processing of terminal leaf data**

RNA isolated from the terminal leaf series of *Poplulus tremula* was sequenced as described by Mähler *et al.* (2020). The data pre-processing was performed following the guidelines described by Delhomme et al. (2015). The quality of the raw sequence data was assessed using FastQC (v.11.4; Andrews, 2012). Sequence reads originating from ribosomal RNAs (rRNA) were identified and removed using SortMeRNA (v. 2.1b; Kopylova et al., 2012; settings --log --paired_in --fastx--sam --num_alignments 1) using the rRNA sequences provided with SortMeRNA (rfam-5s-database-id98.fasta, rfam-5.8s-database-id98.fasta, silva-arc-16s-database-id95.fasta, silva-bac-16s-database-id85.fasta, silva-euk-18s-database-id95.fasta, silva-arc-23s-database-id98.fasta, silva-bac-23s-database-id98.fasta and silva-euk-28s-database-id98.fasta). Data were then filtered to remove adapters and trimmed for quality using Trimmomatic (v. 0.46; Bolger et al., 2014; settings TruSeq3-PE-2.fa:2:30:10 SLIDINGWINDOW:5:20 MINLEN:50). After both filtering steps, FastQC was run again to ensure that no technical artefacts were introduced. Read counts were obtained using Salmon (v. 0.11.2; Patro et al., 2017).

**Identification of long intergenic non-coding RNAs (lincRNAs)**

We implemented a pipeline to identify putative lincRNAs on the pre-processed data, where default settings were used unless specified. We first *in silico*-normalised the reads to reduce data redundancy and then reconstructed the transcriptome using a de-novo assembler, Trinity (v. 2.8.3; Grabherr et al., 2011; Haas et al., 2013).

On the set of transcripts assembled by Trinity, we ran the following programs, which are detailed below: TransDecoder (version 2.8.3; https://github.com/TransDecoder/TransDecoder/wiki; Haas et al., 2013), GMAP (Genomic Mapping and Alignment Program; v. 2020-11-15; settings -i 70000; T. D. Wu & Watanabe, 2005), Salmon Index (v. 0.11.2; Patro et al., 2017), PLEK (predictor of long non-coding RNAs and messenger RNAs based on an improved k-mer scheme; v. 1.2; settings -minlength 200; A. Li et al., 2014), CNCI (Coding-Non-Coding Index; v. 2; Sun et al., 2013), CPC2 (Coding Potential Calculator version 2; v. 2.0 beta; settings -r TRUE; Kang et al., 2017), and BEDTools closest (v. 2.30.0; https://bedtools.readthedocs.io/en/latest/content/tools/closest.html; Quinlan & Hall, 2010).

TransDecoder was run to evaluate transcript coding potential. GMAP was used to map the transcriptome to the genome reference. A Salmon Index was used to subsequently run Salmon (Patro et al., 2017), a pseudoalignment tool used to map RNASeq reads to the assembled transcripts for expression quantification. PLEK, CNCI, and CPC2 were run to classify transcripts as coding or non-coding. Only GMAP uniq results were kept, pre-sorted by chromosome and then start position, to run BEDTools closest on them to exclude the overlap between the identified transcripts and a reference annotation. We then filtered all these results, considering lincRNAs characteristics. We retained only transcripts being expressed in the dataset, that were identified as having no coding potential by the three classification programs and being longer than 200 nt. We kept only transcripts identified as having no coding potential by TransDecoder and having a distance > 1000 nt from the reference genome according to BEDTools closest.

**Gene network inference**

The DESeq2 package (v. 1.42.0; Love et al., 2014) was used for differentially expression analysis with the formula based on consecutive leaf developmental series, both for genes and lincRNAs. Then, lincRNAs and genes expression data were transformed to homoscedastic, asymptotically log2 counts using the variance stabilising transformation as implemented in DESeq2 (settings blind=FALSE). We retained all the genes and lincRNAs being different from na in the dds results and used their values from the aware variance stabilising transformation as an input for the co-expression network. Then, ten network inference methods were run using the Seidr toolkit (Schiffthaler et al., 2023). The networks were aggregated using the inverse rank product method (Zhong et al., 2014) and edges were filtered according to the noise corrected backbone (Coscia & Neffke, 2017). We selected backbone8 to be used for further analysis for example, this is the AspLeaf network in the exNet tool in PlantGenIE.

**Genome assembly figures and data processing**

We used the karyoplotR (Gel & Serra, 2017) package for the R statistical framework (R Core Team, 2019) to generate karyoplots. If not otherwise specified, we omitted irrelevant arguments (such as file paths, parallelism) from command lines in the interests of clarity.
If no arguments are specified, we did not make any changes to the defaults. Unless otherwise specified, we aligned genomic data with BWA mem v0.7.8-r455 (Li, 2013) and RNA-Seq data with STAR v2.6.1d (Dobin *et al.*, 2013). Coverage values assume a genome size of 480Mbp. All scripts and config files can be found in the Git repository: <https://github.com/bschiffthaler/aspen-v2>

**Structural rearrangements between *Populus tremula* and *P. trichocarpa***

To identify homologous chromosomes between *P. tremula* and *P. trichocarpa* genomes, we used minimap2 (Li, 2018) to align the *P. trichocarpa* genome to the v2.2 *P. tremula* assembly. Dot plots were used to visualise the alignments using a custom R script. The syntenic regions, structural rearrangements (inversions, translocations, and duplications), and the sequence differences (SNPs, indels, highly divergent regions, and so on) between homologous chromosomes were identified using SyRI v1.0 (Goel *et al.*, 2019) with default parameters.

**Detection of synteny and collinearity**

We performed an all-versus-all BLASTP using protein sequences of *P. tremula* and *P. trichocarpa* to identify homologous gene pairs between the two species. MCscanX (Wang *et al.*, 2012) was used to identify syntenic gene blocks based on collinear genes requiring at least five gene pairs per syntenic block. For each duplicate gene pair within a syntenic block, we aligned the protein sequences using MAFFT (Katoh & Standley, 2013) and converted the protein alignments into the corresponding codon alignments. Then, we used the maximum likelihood-based program codeml in the PAML package (Yang, 2007), with the model of one *Ka*/*Ks* (non-synonymous/synonymous substitution) ratio for all branches, i.e., model=0, to estimate the *Ks* and *Ka*/*Ks* for each gene pair.

**Phylogenetic and gene family analysis**

We used 32 plant genomes, including 14 Rosid species, seven Superasterid species, one Early-diverging eudicotyledon species, one Magnoliidae species, one Monocotyledoneae species, one basal Angiosperm species, six Acrogymnospermae, and one Bryophyta species (Table S1, Appendix S1) to perform gene family analysis. To remove redundancy caused by alternative splicing variations, only the longest protein sequence at each gene locus was retained for downstream analysis. Orthofinder v2.2.7 (Emms & Kelly, 2015) with default settings was used to cluster the genes into gene families. Gene family trees were constructed using the PLAZA pipeline (Van Bel *et al.*, 2018), which combines MSA_TOOL and TREE_TOOL for multiple sequence alignment and tree inference. We used muscle as the multiple sequence alignment method and fasttree as the tree construction method. The species tree was inferred using STAG (https://github.com/davidemms/STAG) with the gene family tree with a maximum of five copies for each gene. To estimate the divergence time, we first calibrated the species tree based on the divergence dates from Timetree (<http://www.timetree.org/>) by fixing the Embryophyta clade at 532 Mya (million years ago), the Spermatophyta clade at 313 Mya, the Magnoliophyta clade at 181 Mya, the Mesangiospermae clade at 160 Mya, the eudicotyledons clade at 128 Mya, the Pentapetalae clade at 117 Mya, the Pinus clade at 73 Mya, the Salicaceae at 36 Mya, the *Populus* at 16.9 Mya, and inferred the divergence time on each clade using r8s (Sanderson, 2003). We inferred the expansion and contraction of the gene families using CAFÉ (De Bie *et al.*, 2006) and the species tree.

**Common garden experiments**

We measured phenotypic traits in three aspen (*P. tremula*) collections, two of which originate from Sweden and one from Scotland. The Swedish Aspen (SwAsp) collection of 113 individuals, collected across ten degrees of latitude and longitude (Luquez *et al.*, 2008), is replicated in two common gardens in Sweden, one in the north (Sävar, ~64 °N) and one in the south (Ekebo, ~56 °N). The Umeå Aspen (UmAsp) collection comprises 245 individuals originating from the Umeå municipality (~5,200 km^2^) in northern Sweden (Fracheboud *et al.*, 2009; Robinson *et al.*, 2014) growing in a common garden locally, at Sävar. For monitoring purposes, UmAsp also includes some outgroups, the first comprising two genotypes (L1 and L2) from ~66 °N, the second comprising one surviving genotype (K2, K1 having died) from ~62 °N and, the third comprising three genotypes (SwAsp89, SwAsp 95, and SwAsp 97, excluded from the UmAsp GWAS) from SwAsp, and the fourth an aspen from Vindeln (V, 64.2 °N, 19.8 °E), the subject of a previous detailed study of gene expression during wood development (Sundell *et al.*, 2017) and leaf development (Mähler *et al.*, 2020). The Scottish Aspen (ScotAsp) collection of trees distributed across ~3.5 latitudinal and longitudinal degrees in Scotland, was cloned and grown in plots of five trees per clone, in a common garden at Forest Research, Roslin, UK (~56 °N, with 126 genotypes, Harrison, 2009). The original locations of the SwAsp, UmAsp and ScotAsp genotypes are provided in Fig. 2 of the main text.

**Sample collection and sequencing**

Details of the SwAsp and ScotAsp DNA sequencing and SNP calling have been described previously (Rendón-Anaya *et al.*, 2021, Table S2). Samples sequenced from the previously generated data set comprising 94 genotypes from the SwAsp collection (Wang *et al.*, 2018) have been complemented with a further five genotypes re-sequenced for *P. tremula* v2.2. Additional leaf material was sampled from the UmAsp collection common garden at Sävar, northern Sweden. UmAsp is an expansion of the sub-population of the ten SwAsp individuals in the Umeå region (~64 °N). Since the UmAsp collection is drawn from a local geographic area it is not constrained by the growth-confounding effect of latitude that we observe in SwAsp. DNA from these additional samples (ENA study accession PRJEB47451, Table S2) was extracted and sequenced exactly as detailed by Rendón-Anaya *et al.* (2021). Historical climate variables (from 1970 to 2000) at the geographic origins of the samples (where available) were downloaded from WorldClim 2 (Fick *et al.,* 2017) at 10 minute resolution and mean values calculated for each of ScotAsp, SwAsp and UmAsp (Supplementary table S11).

**Mapping and Single Nucleotide Polymorphism calling**

Re-sequenced accessions were mapped against the reference genome of *P. tremula* v2.0 using BWA (v0.7.17) – mem using default parameters. Post-mapping filtering removed reads with MQ<20 (samtools v1.10); depth and breadth of coverage were assessed in order to confirm all samples had a minimum coverage of 8X; finally, we tagged duplicate reads (picard MarkDuplicates v2.10.3; http://broadinstitute.github.io/picard/) before the variant calling step. Duplicated reads did not exceed 14% in any sample (range 3 to 13.8%).

We used GATK v3.8 to call variants. First, we performed a local realignment around indels with RealignerTargetCreator and IndelRealigner (default parameters). We called per-sample variants using HaplotypeCaller to produce gVCF files (-ERC GVCF). Given the large number of individuals, we produced intermediate gVCF files of 50-60 samples using CombineGVCFs that were finally used in the joint-call step with GenotypeGVCFs. We joint-called SNPs for individuals belonging to SwAsp, UmAsp and ScotAsp independently, producing three VCF files from which we selected the SNPs using SelectVariants and filtered them with VariantFiltration (QD < 2.0; FS > 60.0; MQ < 40.0; ReadPosRankSum < -8.0; SOR > 3.0; MQRankSum < -12.5). At this point, the filtered VCF was lifted over to version 2.2 of the genome of *P. tremula*, using picard LiftoverVcf. Further SNP pruning with VCF/BCFtools (Danecek *et al.,* 2021) removed positions with extreme depth values (for consistency with the filtering thresholds described by Rendón-Anaya *et al.* (2021) we set min-meanDP 10, max-meanDP 25), absent in more than 30% of the samples, non-biallelic or displaying an excess of heterozygosity (FDR <0.01).

We used PLINK (v1.90b4.9) to compute the relatedness between samples from the UmAsp collection by calculating genome-wide estimates of identity by descent (IBD) on pruned, unlinked SNPs. We removed from our downstream analyses one sample of each closely related pair at a threshold of 0.4 resulting in 227 unrelated UmAsp individuals.

We called VCF files independently for SwAsp, UmAsp and ScotAsp (with 99, 227 and 105 unrelated individuals, respectively), containing biallelic, high quality sites along the 19 chromosomes using BCFtools and using the BCFtools plugin fill-tags we re-calculated important INFO statistics for the newly generated VCF file (AF, AC, AC_Hemi, AC_Hom, AC_Het, AN, ExcHet, HWE, MAF, NS) that we used to filter excessively heterozygous (FDR < 0.01) sites, low frequency sites, and sites with Hardy-Weinberg equilibrium exact test *P*-values below 1e^-6^ with PLINK (--geno 0.05 -- hwe 0.000001).

**SNP pruning to open chromatin regions**

Detection of phenotype-SNP associations is challenging for complex traits and theoretically requires populations of thousands of individuals to detect loci of small effects. The high-quality genotyping in our aspen populations identified several million SNPs, resulting in a considerable challenge to detect associations after correction for multiple testing. To overcome this obstacle, SNP pruning is an obvious approach to reduce the number of SNPs. We made a subset of all SNPs by intersecting with a bed file of the open chromatin regions identified by ATAC-Seq. The resulting distribution had greater proportions of SNPs in exons and regulatory regions, and a smaller proportion of SNPs in intergenic regions. Based on the interpretation that the subset of SNPs in accessible chromatin regions, as sites of DNA regulatory elements, would have more biologically relevant effects on the phenotype, we conducted genome-wide association analyses on these sets of SNPs, hereafter termed ‘SNPs in Open Chromatin regions’ or ‘OCR SNPs’. These subsets comprised 185,616 SNPs in SwAsp, 220,009 in UmAsp and 212,902 in ScotAsp.

**SwAsp collection RNA-Sequencing analysis and eQTL mapping**

We used existing RNA-Seq data from Mähler *et al*. (2017; ENA accession ERP014886), which was reprocessed using the v2.2 genome assembly. The RNA-Seq data were processed using salmon/1.0.0 (Patro *et al.*, 2017) in mapping-based mode, specifically utilising the “genome-aware” quantification index. Decoy metadata were extracted from the *P. tremula* v2.2 genome using grep -E "^>" genome.fa | cut -f 1, then genome and transcriptome (of the same assembly) were concatenated to form the salmon “gentrome”. Salmon was then run using options -l A --seqBias --gcBias -p 8 --validateMappings -g transcript_gene_map.tsv, where “transcript_gene_map.tsv” is a tab delimited file mapping transcript IDs to gene IDs for the assembly. The gene expression values for the population were applied as the phenotypic gene expression data to map expression-QTL (eQTL) in the SwAsp population, following the methods applied in Mähler *et al.* (2017) updated to the v2.2 SNP data set and named “Matrix eQTLs”. Owing to the potential sensitivity of GWAS analyses to phenotypic data distributions, we also mapped eQTL for the same gene expression data set using fastJT (Lin *et al.,* 2019). For the non-parametric genome-transcriptome eQTL mapping, the mean expression of genotypes was associated with SNPs using the R package fastJT v. 1.0.6 (Lin *et al.,* 2019). The analysis required converting the Variant Call Format (VCF) file into .raw format via PLINK v. 1.9, then transformation into .tsv format. To account for multiple testing, we applied an FDR adjustment to the p-values using the R package stats. Only SNPs significantly associated (FDR < 0.05) were kept for downstream analyses. For simplicity, eQTLs were defined as *cis-*acting (local) if the gene and SNP collocated to the same chromosome, and *trans-*acting (distant) if the expressed gene was on a different chromosome to its associated SNP.

**Leaf physiognomy phenotyping**

Twenty-eight leaf physiognomy (shape and size) parameters (Appendix S2) were measured in six leaves of three clonal replicate trees in the UmAsp common garden on 9th July 2019, and fifteen leaves sampled across five clonal replicates per genotype in the ScotAsp Roslin clone garden on 19th June 2012. Mature, undamaged leaves were sampled, scanned using a flatbed scanner, and measured using LAMINA software (Bylesjö *et al.,* 2008) following methods described in Mähler *et al.* (2020). In addition to the default metrics in LAMINA, seven additional composite metrics were calculated (Appendix S2). Leaves sampled from the SwAsp common gardens as reported in Mähler *et al.* (2020) for leaf area, leaf circularity and leaf indent depth, are presented here for the further 25 leaf shape and size metrics for the analysis with SNPs called from *P. tremula* v2.2.

**SwAsp, UmAsp and ScotAsp genome-wide association mapping**

Inspired by the pre-GWAS steps described in Slaten *et al*. (2020), a pipeline (available at <https://github.com/sarawestman/Genome_paper>) was developed to calculate Best Linear Unbiased Predictor (BLUP) values for the phenotypes used in the GWAS; the main steps are as follows. Outlier values in the phenotype data were identified using the ‘OutlierTest’ function of the R package ‘car’ (version 3.0.10, Fox & Weisberg, 2019) and subsequently removed. Each trait was then tested for non-normally distributed random effects and/or residual error terms using the Shapiro-Wilk test in R, and those at *P*-value < 0.05 were transformed using the ‘orderNorm’ function of the R package ‘bestNormalize’ (version 1.6.1, Peterson & Cavanaugh, 2019). For all SwAsp individuals, we calculated a best unbiased linear predictor (BLUP) with a restricted maximum likelihood approach to estimate the genotypic effect of each phenotype, as described in Wang *et al.* (2018). Briefly, the model used was:

z*_jkl_* = u + b*_j_* + g*_k_* + e*_jkl_*

Where in the equation z_jkl_ is the phenotype of the *lth* individual in the *jth* block from the *kth* genotype, where u denotes the grand mean and e_jkl_ is the residual error term. The genotype and residual terms were treated as random effects, whereas block was a fixed effect. For UmAsp individuals, where some individuals were replanted in a subsequent year, the model was:

z*_ijkl_* = u + b*_i_* + y_j_ + g*_k_* + e*_ijkl_*

Where in the equation z_ijkl_ is the phenotype of the *lth* individual in the *ith* block in the *j*th planting year from the *kth* genotype, where u denotes the grand mean and e_ijkl_ is the residual error term. The genotype and residual terms were treated as random effects, whereas the block and planting year were fixed effects. For ScotAsp (Roslin garden) leaf physiognomy data, the model used was:

z*_ij_* = u + g*_i_* + e*_ij_*

Where in equation z_ij_ is the phenotype of the i*th* genotype, where u denotes the grand mean and e_ij_ is the residual error term. The genotype and residual terms were treated as random effects.

For each phenotype, BLUP data were tested for departure from a normal distribution using a Shapiro-Wilk test (shapiro.test in R) and residuals plotted to check for homogeneity of variance. Effects were considered significant at *P*-value < 0.05.

We used these BLUP estimates as leaf physiognomy phenotypic values for genome-wide association (GWA) mapping in each of the SwAsp, UmAsp and ScotAsp collections separately. GWAS was carried out on the SwAsp (maxiumum 99 genotypes), UmAsp (max. 227 genotypes), and ScotAsp (max. 105 genotypes) for which both BLUP values and SNP data were available. SNPs were filtered for a minor allele frequency of 5% and Hardy-Weinberg equilibrium *P-*value threshold of 1e^-6^ using PLINK version 1.9. (Purcell *et al.,* 2007). Genome-wide associations were investigated using GEMMA v0.98.1 (Zhou & Stephens, 2012). Linear mixed models were used as described by Wang *et al.* (2018) with the centred relatedness matrix of all individuals applied as a covariate. For all traits we cautiously applied latitude of genotype origin as an additional covariate to account for the potential influence of latitude-driven bud set that strongly influences growth and some phenological phenotypes (Luquez *et al.*, 2008; Wang *et al.*, 2018). A 5% false discovery rate (*q*-value) was used to determine significant associations, calculated in the 'qvalue' package in R (Storey *et al.,* 2021). The proportion of phenotypic variation explained (PVE) by an individual SNP was estimated using the equation stated in Wang *et al.* (2018). SNPs were filtered for plotting purposes using ‘dplyr’ version 0.8.4 in R (Wickham *et al.*, 2020). GWAS Manhattan plots were generated in R using ‘qqman’ version 0.1.4 (Turner, 2017).

**Linkage disequilibrium (LD) analyses**

We calculated linkage-disequilibrium *r*^2^ values among SNPs of interest using --r2 with-freqs in PLINK.

**Gene ontology and Pfam enrichments of GWAS results**

We used the Enrichment tool at PlantGenIE.org to conduct gene ontology (GO) and Pfam enrichments for lists of genes of interest, selecting “AspLeaf expessed genes (32832)” as the test background. In addition to the default PlantGenIE GO tables, we used the “Send to REVIGO” button to reduce and visualise the GO results using the REVIGO tool (Supek *et al.,* 2011). Default settings were used in REVIGO except for list size, which was set to 0.4, and species, set to *Arabidopsis thaliana*. We then viewed the Scatterplot for Biological Process GO terms and selected “Export to R script for plotting” to plot the GO visualisation in R and followd the default, exported scripts.

**Heritability and genetic correlations**

We estimated heritability as ‘SNP heritability’, an approximation of narrow sense heritability or *h^2^* (Kruijer *et al.*, 2015) based on both phenotypic data and the relatedness matrix obtained from GEMMA (see details of genome-wide association mapping above). This marker-based heritability (*h*^2^) was determined using ‘marker_h2’ function in the ‘heritability’ package version 1.3 (Kruijer, 2019) in R. Genetic correlations among the 26 phenotypes in the GWAS were calculated for the UmAsp collection following the method described in Mähler *et al.* (2020).

**Signatures of positive and balancing selection**

SNPs in SwAsp, UmAsp, and ScotAsp were discarded if the missing rate was >5% and failed the Hardy–Weinberg equilibrium test (*P*-value < 1e^-6^). SNPs were annotated using ANNOVAR v2019Oct24 with the parameters --neargene 2000 -geneanno -dbtype refGene. SNPs with minor allele frequency >10% and missing rate <20% were used for linkage disequilibrium (LD) analysis. We calculated the squared correlation coefficients (*r*^2^) between all pairs of SNPs that were within 50 Kbp using PopLDdecay v3.41 (Zhang *et al*., 2018). To analyse the population structure based on the PCA, SNPs were pruned by removing one SNP from each pair of SNPs with a between SNP correlation coefficient (*r*^2^) > 0.2 in windows of 50 SNPs with a step of 5 SNPs using PLINK v1.90b6.16 (Purcell *et al*., 2007). The smartpca program in EIGENSOFT v6.1.4 (Patterson *et al.,* 2006) was then used to perform a principal components analysis (PCA) on the reduced set of genome-wide independent SNPs. The significance of the principal components was determined using a Tacey-Widom test in smartpca in EIGENSOFT. To identify signals of positive selection, we calculated genome-wide Weir and Cockerham’s *F*_ST_ (Weir & Cockerham, 1984), π ratios (i.e., π·_Um_/π·_Sw_, π·_Scot_/π·_Sw_, π·_Sw_/π·_Um_, and π·_Scot_/π·_Um_) and Tajima’s D (Tajima, 1989) in 10 Kbp non-overlapping windows across the chromosomes in SwAsp, UmAsp, and ScotAsp using VCFtools v0.1.15 (Danecek *et al.*, 2011). We removed the genotypes from SwAsp population 9 when calculating the *F*_ST_ and π ratios as this population contains individuals from Umeå that overlap with UmAsp.

We also calculated the composite likelihood ratio (CLR) statistic in 10 Kbp non-overlapping windows using SweepFinder2 and iHH12 (Integrated Haplotype Homozygosity Pooled) using selscan v1.3., after inferring the haplotype phase and imputing the missing alleles with Beagle v5.1 (Browning *et al.,* 2018) with default parameters. We averaged the normalised iHH12 of each biallelic SNP site in 10 Kbp non-overlapping windows.

To identify regions under positive selection, sliding windows containing at least 10 SNPs were used as input to a range of inference methods. Windows with the lowest 5% Tajima’s D values or highest 5% values for other measures were considered as those windows displaying evidence of signals of positive selection. To limit false positives, we required that the sliding windows classified as being under selection were identified by at least three measures and that one of those measures must be either CLR or iHH12. Adjacent windows were merged. Genes or SNPs within these selected regions were assumed to be under selection.

We ran betascan (Siewert *et al.*, 2017) (-fold -m 0.1) to detect possible signals of balancing selection in the ScotAsp, UmAsp and SwAsp collections. We filtered GWAS results for all traits in SwAsp, UmAsp and ScotAsp at -log_10_(*P*_wald_) ≥ 5 and compared the SNP frequencies in 10 Kbp windows with the ß score frequencies (P_FDR_ ≤ 0.01) for the three collections. Associated gene functions were examined for SNPs in coincident high frequency regions for GWAS and ß score signals.

**References**

**Altschul SF, Gish W, Miller W, Myers EW, Lipman DJ**. **1990**. Basic local alignment search tool. *Journal of Molecular Biology* **215**: 403–410. https://doi.org/10.1016/S0022-2836(05)80360-2

**Andrews S. 2012.** FastQC: A quality control application for high throughput sequence data. Babraham Institute Project Page: http://www.bioinformatics.bbsrc.ac.uk/Projects/Fastqc.

**Apuli R-P, Bernhardsson C, Schiffthaler B, Robinson KM, Jansson S, Street NR, Ingvarsson PK**. **2020**. Inferring the genomic landscape of recombination rate variation in European Aspen (*Populus tremula*). *Genes|Genomes|Genetics* **10**: 299–309. https://doi.org/10.1534/g3.119.400504

**Bolger AM, Lohse M, Usadel B**. **2014**. Trimmomatic: A flexible trimmer for Illumina sequence data. *Bioinformatics* **30**: 2114–2120. https://doi.org/10.1093/bioinformatics/btu170

**Browning BL, Zhou Y, Browning SR**. **2018**. A one-penny imputed genome from next generation reference panels. *Am. J. Hum. Genet.* **103**: 338–348. doi:10.1016/j.ajhg.2018.07.015

**Bucchini F, Del Cortona A, Kreft L, Botzki A, Van Bel M, Vandepoele K**. **2021**.TRAPID 2.0: a web application for taxonomic and functional analysis of de novo transcriptomes. *Nucleic Acids Research* **49**: e101. https://doi.org/10.1093/nar/gkab565

**Buchfink B, Xie C, Huson DH**. **2014**. Fast and sensitive protein alignment using DIAMOND. *Nature Methods* **12**: 59–60. https://doi.org/10.1038/s41592-021-01101-x

**Bylesjö M, Segura V, Soolanayakanahally RY, Rae AM, Trygg J, Gustafsson P, Jansson S, Street NR.** **2008**. LAMINA: a tool for rapid quantification of leaf size and shape parameters. *BMC Plant Biol.* **81**: 1–9. https://doi.org/10.1186/1471-2229-8-82

**Campbell MS, Holt C, Moore B, Yandell M**. **2014**. Genome Annotation and Curation Using MAKER and MAKER-P. *Current Protocols in Bioinformatics* **2014**: 4.11.1-4.11.39. https://doi.org/10.1002/0471250953.bi0411s48

**Chin CS, Peluso P, Sedlazeck FJ, Nattestad M, Concepcion GT, Clum A, Dunn C, O’Malley R, Figueroa-Balderas R, Morales-Cruz A, *et al.*** **2016**. Phased diploid genome assembly with single-molecule real-time sequencing. *Nature Methods* **13**: 1050–1054. https://doi.org/10.1038/nmeth.4035

**Coscia M, Neffke FMH. 2017.** Network backboning with noisy data. *Proceedings - International Conference on Data Engineering,* 425–436. https://doi.org/10.1109/ICDE.2017.100

**Danecek P, Auton A, Abecasis G, Albers CA, Banks E, DePristo MA, Handsaker RE, Lunter G, Marth GT, Sherry ST, McVean G, Durbin R, 1000 Genomes Project Analysis Group**. 2011. The variant call format and VCFtools. *Bioinformatics* **27**: 2156–2158. https://doi.org/10.1093/bioinformatics/btr330

**Danecek P, Bonfield JK, Liddle J, Marshall J, Ohan V, Pollard MO, Whitwham A, Keane T, McCarthy SA, Davies RM, Li H. 2021.** Twelve years of SAMtools and BCFtools. *GigaScience* **10:** giab008, https://doi.org/10.1093/gigascience/giab008

**Davies C, Ellis CJ, Iason GR, Ennos RA**. **2014**. Genotypic variation in a foundation tree (*Populus tremula* L.) explains community structure of associated epiphytes. *Biol. Lett.* **10**: 20140190. https://doi.org/10.1098/rsbl.2014.0190

**De Bie T, Cristianini N, Demuth JP, Hahn MW**. **2006**. CAFE: a computational tool for the study of gene family evolution. *Bioinformatics* **22**: 1269–71. https://doi.org/10.1093/bioinformatics/btl097

**Delhomme, N., Mähler, N., Schiffthaler, B., Sundell, D., Mannapperuma, C., Hvidsten, T. R., & Street, N. R. (2015).** Guidelines for RNA-Seq data analysis. Retrieved September 5, 2023, from http://www.epigenesys.eu

**Dobin A, Davis CA, Schlesinger F, Drenkow J, Zaleski C, Jha S, Batut P, Chaisson M, Gingeras TR**. **2013**. STAR: ultrafast universal RNA-seq aligner. *Bioinformatics* **29**: 15–21. https://doi.org/10.1093/bioinformatics/bts635

**El-Gebali S, Mistry J, Bateman A, Eddy SR, Luciani A, Potter SC, Qureshi M, Richardson LJ, Salazar GA, Smart A, *et al.*** **2018**. The Pfam protein families database in 2019. *Nucleic Acids Research* **47**: 427–432. https://doi.org/10.1093/nar/gkaa913

**Emms DM, Kelly S**. **2015**. OrthoFinder: solving fundamental biases in whole genome comparisons dramatically improves orthogroup inference accuracy. *Genome Biology* **16**. https://doi.org/10.1186/s13059-015-0721-2

**Fick SE, and Hijmans RJ. 2017.** WorldClim 2: new 1km spatial resolution climate surfaces for global land areas. *International Journal of Climatology* **37:** 4302-4315**.** https://doi.org/10.1002/joc.5086

**Hoff KJ, Lange S, Lomsadze A, Borodovsky M, Stanke M**. **2016**. BRAKER1: Unsupervised RNA-Seq-Based Genome Annotation with GeneMark-ET and AUGUSTUS *Bioinformatics* **32**: 767–769. https://doi.org/10.1093/bioinformatics/btv661

**Huang S, Kang M, Xu A**. **2017**. HaploMerger2: rebuilding both haploid sub-assemblies from high-heterozygosity diploid genome assembly. *Bioinformatics* **33**: 2577–2579. https://doi.org/10.1093/bioinformatics/btx220

**Kanehisa M, Goto S**. **2000**. KEGG: Kyoto Encyclopedia of Genes and Genomes. *Nucleic Acids Research* **28**: 27–30. https://doi.org/10.1093/nar/28.1.27

**Kang YJ, Yang DC, Kong L, Hou M, Meng YQ, Wei L, Gao G. 2017.** CPC2: a fast and accurate coding potential calculator based on sequence intrinsic features. *Nucleic Acids Research* **45:** W12–W16. https://doi.org/10.1093/NAR/GKX428

**Katoh K, Standley DM**. **2013**. MAFFT multiple sequence alignment software version 7: Improvements in performance and usability. *Molecular Biology and Evolution* **30**: 772–780. https://doi.org/10.1093/molbev/mst010

**Kersten B, Faivre Rampant P, Mader M, Le Paslier M-C, Bounon R, Berard A, Vettori C, Schroeder H, Leplé J-C, Fladung M**. **2016**. Genome Sequences of *Populus tremula* Chloroplast and Mitochondrion: Implications for Holistic Poplar Breeding. *PLoS ONE* **11**: e0147209. https://doi.org/10.1371/journal.pone.0147209. https://doi.org/10.1371/journal.pone.0147209

**Kopylova E, Noé L, Touzet H**. **2012**. SortMeRNA: fast and accurate filtering of ribosomal RNAs in metatranscriptomic data. *Bioinformatics* **28**: 3211–7. https://doi.org/10.1093/bioinformatics/bts611

**Korf I**. **2004**. Gene finding in novel genomes. *BMC Bioinformatics* **5**: 59. https://doi.org/10.1186/1471-2105-5-59

**Kruijer W**. **2019**. heritability: Marker-Based Estimation of Heritability Using Individual Plant or Plot Data. R package version 1.3. *https://CRAN.R-project.org/package=heritability*.

**Kruijer W, Boer MP, Malosetti M, Flood PJ, Engel B, Kooke R, Keurentjes JJB, Van Eeuwijk FA**. **2015**. Marker-based estimation of heritability in immortal populations. *Genetics* **199**: 379–398. https://doi.org/10.1534/genetics.114.167916

**Li H**. **2013**. Aligning sequence reads, clone sequences and assembly contigs with BWA-MEM. arXiv:1303.3997v2 [q-bio.GN]

**Li H**. **2018**. Minimap2: Pairwise alignment for nucleotide sequences. *Bioinformatics* **34**: 3094–3100. https://doi.org/10.1093/bioinformatics/bty191

**Li A, Zhang J, Zhou Z. 2014.** PLEK: A tool for predicting long non-coding RNAs and messenger RNAs based on an improved k-mer scheme. *BMC Bioinformatics* **15:** 1–10. https://doi.org/10.1186/1471-2105-15-311

**Lin J, Sibley A, Shterev I, Nixon A, Innocenti F, Chan C, Owzar K. 2019.** fastJT: An R package for robust and efficient feature selection for machine learning and genome-wide association studies. *BMC Bioinformatics* **20:** 333. https://doi.org/10.1186/s12859-019-2869-3

**Lin YC, Wang J, Delhomme N, Schiffthaler B, Sundström G, Zuccolo A, Nystedt B, Hvidsten TR, de la Torre A, Cossu RM, *et al.*** **2018**. Functional and evolutionary genomic inferences in Populus through genome and population sequencing of American and European aspen. *Proceedings of the National Academy of Sciences of the United States of America* **115**: E10970–E10978. https://doi.org/10.1073/pnas.1801437115

**Lomsadze A, Ter-Hovhannisyan V, Chernoff YO, Borodovsky M**. **2005**. Gene identification in novel eukaryotic genomes by self-training algorithm. *Nucleic acids research* **33**: 6494–6506. https://doi.org/10.1093/nar/gki937

**Love MI, Huber W, Anders S. 2014.** Moderated estimation of fold change and dispersion for RNA-seq data with DESeq2. *Genome Biology* **15:** 1–21. https://doi.org/10.1186/s13059-014-0550-8

**Luquez V, Hall D, Albrectsen BR, Karlsson J, Ingvarsson P, Jansson S**. **2008**. Natural phenological variation in aspen (*Populus tremula*): The SwAsp collection. *Tree Genetics and Genomes* **4**: 279–292. https://doi.org/10.1007/s11295-007-0108-y

**Mähler N, Wang J, Terebieniec BK, Ingvarsson PK, Street NR, Hvidsten TR**. **2017**. Gene co-expression network connectivity is an important determinant of selective constraint. *PLoS Genetics* **13**: e1006402. https://doi.org/10.1371/journal.pgen.1006402

**Mähler N, Schiffthaler B, Robinson KM, Terebieniec BK, Vučak M, Mannapperuma C, Bailey MES, Jansson S, Hvidsten TR, Street NR**. **2020**. Leaf shape in *Populus tremula* is a complex, omnigenic trait. *Ecol Evol* **10**:11922–11940. https://doi.org/10.1002/ece3.6691

**McKenna, A., Hanna, M., Banks, E., et al. 2010.** The Genome Analysis Toolkit: A MapReduce framework for analyzing next-generation DNA sequencing data. *Genome Res.* **20**: 1297–1303. https://doi.org/10.1101/gr.107524.110

**Patro R, Duggal G, Love MI, Irizarry RA, Kingsford C**. **2017**. Salmon provides fast and bias-aware quantification of transcript expression. *Nature Methods* **14**: 417–419. https://doi.org/10.1038/nmeth.4197

**Patterson N, Price AL, Reich D**. **2006**. Population Structure and Eigenanalysis. *PLoS Genet* **2**: e190. https://doi.org/10.1371/journal.pgen.0020190

**Peterson RA, Cavanaugh JE**. **2020**. Ordered quantile normalization: a semiparametric transformation built for the cross-validation era. *Journal of Applied Statistics* **47**: 2312-2327. doi: 10.1080/02664763.2019.1630372.

**Purcell S, Neale B, Todd-Brown K, Thomas L, Ferreira MAR, Bender D, Maller J, Sklar P, De Bakker PIW, Daly MJ, *et al.*** **2007**. PLINK: A tool set for whole-genome association and population-based linkage analyses. *American Journal of Human Genetics* **81**: 559–575. https://doi.org/10.1086/519795

**Quinlan AR, Hall IM. 2010.** BEDTools: a flexible suite of utilities for comparing genomic features. *Bioinformatics* **26:** 841–842. https://doi.org/10.1093/BIOINFORMATICS/BTQ033

**R Core Team**. **2019**. A Language and Environment for Statistical Computing. *R Foundation for Statistical Computing*, Vienna, Austria. https://www.R-project.org/

**Rendón-Anaya M, Wilson J, Sveinsson S, Cottrell J, Bailey ME, Ruņģis D, Lexer C, Jansson S, Robinson KM, Street NR, Ingvarsson PK**. **2021**. Adaptive Introgression Facilitates Adaptation to High Latitudes in European Aspen (*Populus tremula* L.). *Molecular Biology and Evolution* **38**: 5034–5050. https://doi.org/10.1093/molbev/msab229

**Robinson KM, Delhomme N, Mähler N, Schiffthaler B, Onskog J, Albrectsen BR, Ingvarsson PK, Hvidsten TR, Jansson S, Street NR**. **2014**. *Populus tremula* (European aspen) shows no evidence of sexual dimorphism. *BMC plant biology* **14**: 276. https://doi.org/10.1186/s12870-014-0276-5

**Sanderson MJ**. **2003**. r8s: inferring absolute rates of molecular evolution and divergence times in the absence of a molecular clock. *Bioinformatics* **19**: 301–2. https://doi.org/10.1093/bioinformatics/19.2.301

**Schiffthaler B, Van Zalen E, Serrano AR, Street NR, Delhomme N. 2023.** SeiÃ°r: Efficient calculation of robust ensemble gene networks. *Heliyon*, **9:** e16811. https://doi.org/10.1016/j.heliyon.2023.e16811

**Siewert KM, Voight BF. 2017.** Detecting Long-Term Balancing Selection Using Allele Frequency Correlation. *Mol Biol Evol* 2017, **34:** 2996–3005. https://doi.org/10.1093/molbev/msx209

**Slaten ML, Chan YO, Shrestha V, Lipka AE, Angelovici R**. **2020**. HAPPI GWAS: Holistic Analysis with Pre- and Post Integration GWAS. *Bioinformatics* **36**: 4655–4657. https://doi.org/10.1093/bioinformatics/btaa589

**Stanke M, Diekhans M, Baertsch R, Haussler D**. **2008**. Using native and syntenically mapped cDNA alignments to improve de novo gene finding. *Bioinformatics* **24**: 637–44. https://doi.org/10.1093/bioinformatics/btn013

**Storey JD, Bass AJ, Dabney A, Robinson D**. **2021**. qvalue: Q-value estimation for false discovery rate control. R package version 2.26.0, http://github.com/jdstorey/qvalue.

**Sun L, Luo H, Bu D, Zhao G, Yu K, Zhang C, Liu Y, Chen R, Zhao Y. (2013).** Utilizing sequence intrinsic composition to classify protein-coding and long non-coding transcripts. *Nucleic Acids Research* **41:** e166–e166. https://doi.org/10.1093/NAR/GKT646

**Sundell D, Mannapperuma C, Netotea S, Delhomme N, Lin Y-C, Sjödin A, Van de Peer Y, Jansson S, Hvidsten TR, Street NR**. **2015**. The Plant Genome Integrative Explorer Resource: PlantGenIE.org. *New Phytologist* **208**: 1149–1156. https://doi.org/10.1111/nph.13557

**Sundell D, Street NR, Kumar M, Mellerowicz EJ, Kucukoglu M, Johnsson C, Kumar V, Mannapperuma C, Delhomme N, Nilsson O, *et al.*** **2017**. Aspwood: High-spatial-resolution transcriptome profiles reveal uncharacterized modularity of wood formation in *Populus tremula*. *Plant Cell* **29**: 1585–1604. https://doi.org/10.1105/tpc.17.00153

**Supek F, Bošnjak M, Škunca N, Šmuc T. 2011**. REVIGO summarizes and visualizes long lists of gene ontology terms. *PLoS One*. **6:** e21800. doi: 10.1371/journal.pone.0021800

**Tajima F**. **1989**. Statistical method for testing the neutral mutation hypothesis by DNA polymorphism. *Genetics* **123**: 585-595. https://doi.org/10.1093/genetics/123.3.585

**Tang H, Zhang X, Miao C, Zhang J, Ming R, Schnable JC, Schnable PS, Lyons E, Lu J**. **2015**. ALLMAPS: Robust scaffold ordering based on multiple maps. *Genome Biology* **16**: 3. https://doi.org/10.1186/s13059-014-0573-1

**The UniProt Consortium**. **2019**. UniProt: a worldwide hub of protein knowledge The UniProt Consortium. *Nucleic Acids Research* **47**: D506–D515. https://doi.org/10.1093/nar/gky1049

**Tibbits JFG, McManus LJ, Spokevicius A V., Bossinger G**. **2006**. A rapid method for tissue collection and high-throughput isolation of genomic DNA from mature trees. *Plant Molecular Biology Reporter* **24**: 81–91. https://doi.org/10.1007/BF02914048

**Tuskan GA, DiFazio S, Jansson S, Bohlmann J, Grigoriev I, Hellsten U, Putnam M, Ralph S, Rombauts S, Salamov A, *et al.*** **2006**. The genome of black cottonwood, Populus trichocarpa (Torr. & Gray). *Science* **313**: 1596–1604. https://doi.org/10.1126/science.1128691

**Van Bel M, Diels T, Vancaester E, Kreft L, Botzki A, Van de Peer Y, Coppens F, Vandepoele K**. **2018**. PLAZA 4.0: an integrative resource for functional, evolutionary and comparative plant genomics. *Nucleic acids research* **46**: D1190–D1196. https://doi.org/10.1093/nar/gkx1002

**Walker BJ, Abeel T, Shea T, Priest M, Abouelliel A, Sakthikumar S, Cuomo CA, Zeng Q, Wortman J, Young SK, *et al.*** **2014**. Pilon: An integrated tool for comprehensive microbial variant detection and genome assembly improvement. *PLoS ONE* **9**: e112963. https://doi.org/10.1371/journal.pone.0112963

**Wang Y, Tang H, Debarry JD, Tan X, Li J, Wang X, Lee T, Jin H, Marler B, Guo H, *et al.*** **2012**. MCScanX: a toolkit for detection and evolutionary analysis of gene synteny and collinearity. *Nucleic acids research* **40**: e49. https://doi.org/10.1093/nar/gkr1293

**Wang J, Ding J, Tan B, Robinson KM, Michelson IH, Johansson A, Nystedt B, Scofield DG, Nilsson O, Jansson S, *et al.*** **2018**. A major locus controls local adaptation and adaptive life history variation in a perennial plant. *Genome Biology* **19**: 1–17. https://doi.org/10.1186/s13059-018-1444-y

**Weir BS, Cockerham CC**. **1984**. Estimating F-statistics for the analysis of population structure. *Evolution* **38**: 1358-1370. https://doi.org/10.1111/j.1558-5646.1984.tb05657.x

**Wickham H, François R, Henry L, Müller K**. **2020**. dplyr: A Grammar of Data Manipulation. R package version 0.8.4. *https://CRAN.R-project.org/package=dplyr*.

**Wu TD, Watanabe CK. 2005.** GMAP: a genomic mapping and alignment program for mRNA and EST sequences. *Bioinformatics* **21:** 1859–1875. https://doi.org/10.1093/BIOINFORMATICS/BTI310

**Yang Z**. **2007**. PAML 4: Phylogenetic Analysis by Maximum Likelihood. *Molecular Biology and Evolution* **24**: 1586–1591. https://doi.org/10.1093/molbev/msm088

**Zhang Y, Liu T, Meyer CA, Eeckhoute J, Johnson DS, Bernstein BE, Nusbaum C, Myers RM, Brown M, Li W. 2008.** Model-based analysis of ChIP-Seq (MACS). *Genome biology* **9**: 1- 9.

**Zhang, M., Zhang, Y., Scheuring, C. et al. 2012.** Preparation of megabase-sized DNA from a variety of organisms using the nuclei method for advanced genomics research. *Nat. Protoc.* **7**: 467–478

**Zhang J, Yang Y, Zheng K, Xie M, Feng K, Jawdy SS, Gunter LE, Ranjan P, Singan VR, Engle N, *et al.*** **2018**. Genome-wide association studies and expression-based quantitative trait loci analyses reveal roles of HCT2 in caffeoylquinic acid biosynthesis and its regulation by defense-responsive transcription factors in *Populus*. *New Phytologist* **220**: 502–516. https://doi.org/10.1111/nph.15297

**Zhou X, Stephens M**. **2012**. Genome-wide efficient mixed-model analysis for association studies. *Nature genetics* **44**: 821–4. https://doi.org/10.1038/ng.2310
