## Supplementary material for "An Improved Chromosome-scale Genome Assembly and Population Genetics resource for *Populus tremula*": Legends for supplementary figures.docx

**Supplementary figure S1.** Density plot of synonymous mutation substitution rates (Ks) distribution. potra = *P. tremula*, potri = *P. trichocarpa*.

**Supplementary figure S2.** Phylogenetic tree constructed from protein sequences from 32 plant genomes used to infer the analyse expansion and contraction of gene families.

**Supplementary figure S3.** Linking the genome-wide associations (GWAS), phenotype and gene expression data in SwAsp. **(A)** Manhattan plot view of the GWAS, showing the two SNPs significant at *q*-value < 0.05 in SwAsp for leaf base angle (Width 75%/Width or in the LAMINA output files, “Horizontal 75%/ Horizontal center”, see full details of phenotypic metrics in Appendix 2). The inset shows the position of the SNPs in a zoomed window from JBrowse in PlantGenIE, with the nearest Potra v2.2 gene models, positions of *cis* eQTL (from the Matrix eQTL method), regions of open chromatin, and the SNP positions. **(B)** Alleles for one of the significant SNPs, chr5_13736867_T_G are plotted with the phenotypic values obtained from files available at the Figshare repository. Comparisons were made with t-tests between the homozygous major reference allele group (TT) and each of the heterozygous allele group (TG) and homozygous minor allele group (GG). Individual data points are superimposed on the boxplot. From the phenotype file and this boxplot view, the SwAsp genotypes with the greatest and smallest values of the phenotype, SwAsp 114 and SwAsp 4, were selected and representative leaf images (also available at Figshare) examined; note the differences in leaf base angle. **(C)** The allelic groups for chr5_13736867_T_G were also plotted with the gene expression (variance normalised transformation, VST, values) of Potra2n5c11907, the gene associated with this SNP from the SwAsp buds gene expression datea set available at Figshare. The *P-*values represent t-test comparisons between the TT (reference) and each of the TG and GG allele groups. Individual data points are shown overlaying the boxplot.

**Supplementary figure S4.** A JBrowse view exported from PlantGenIE.org of a 379 kbp region of chromosome 9 showing how tracks can be loaded to display intersecting *P. tremula* v2.2 gene models and GWAS results. The top bar shows the distance along the chromosome and immediately below are the Potra v2.2 gene models. The top GWAS track is the ScotAsp All-SNP GWAS for leaf indent density SD, with points in dark blue showing the SNPs with significant associations to the trait above the *q*-value threshold of 0.05. The three tracks below this are the ScotAsp OCR-GWAS associations for indent density, indent depth SD and indent width SD, with points in red showing the SNPs with significant associations to the trait above the *q*-value threshold of 0.05.
