## Supplementary figures and images for "An Improved Chromosome-scale Genome Assembly and Population Genetics resource for *Populus tremula*"

### Supplementary figure S1. Ks distribution.pdf

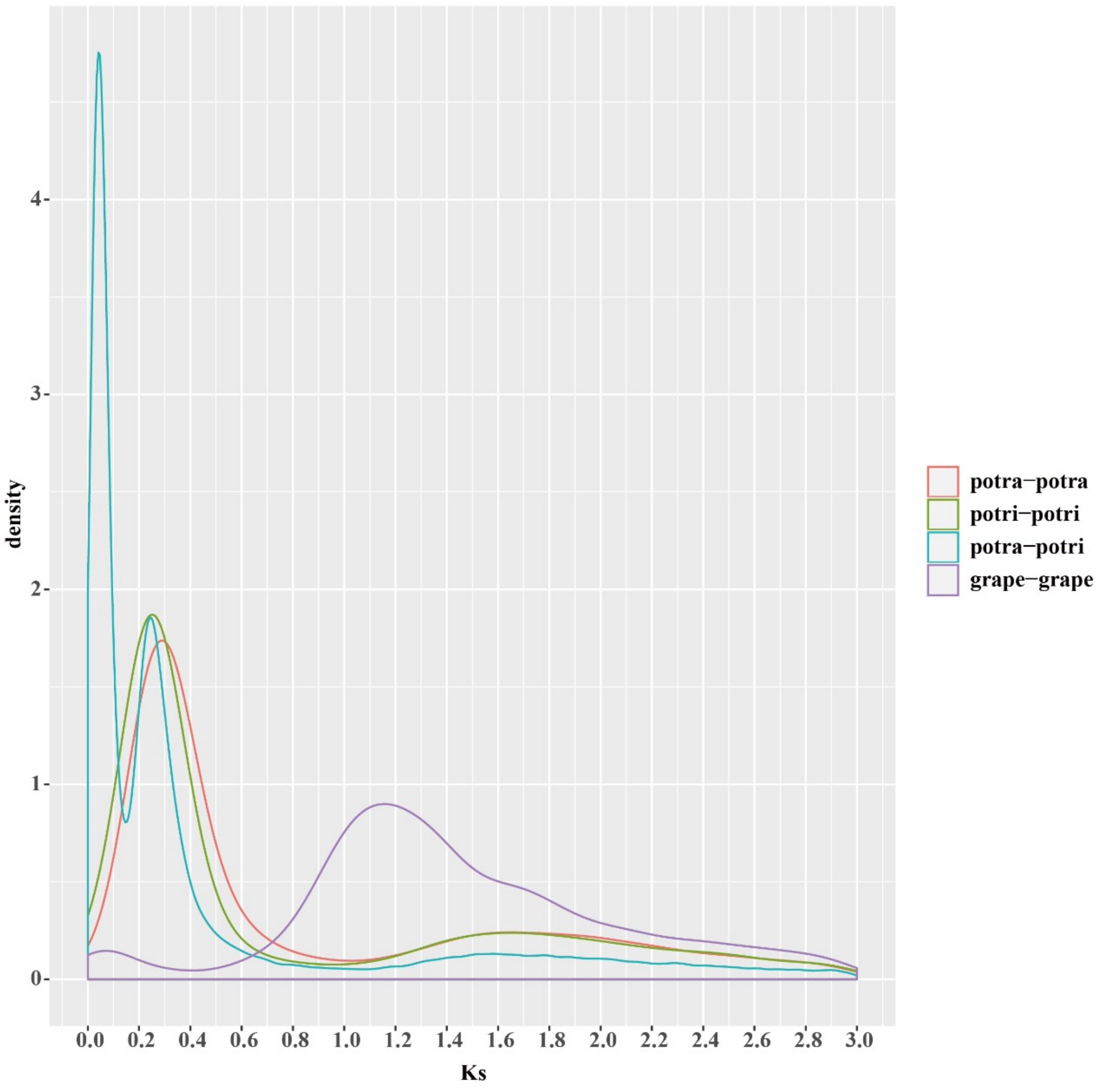

### Supplementary figure S2. Phylogenetic tree.pdf

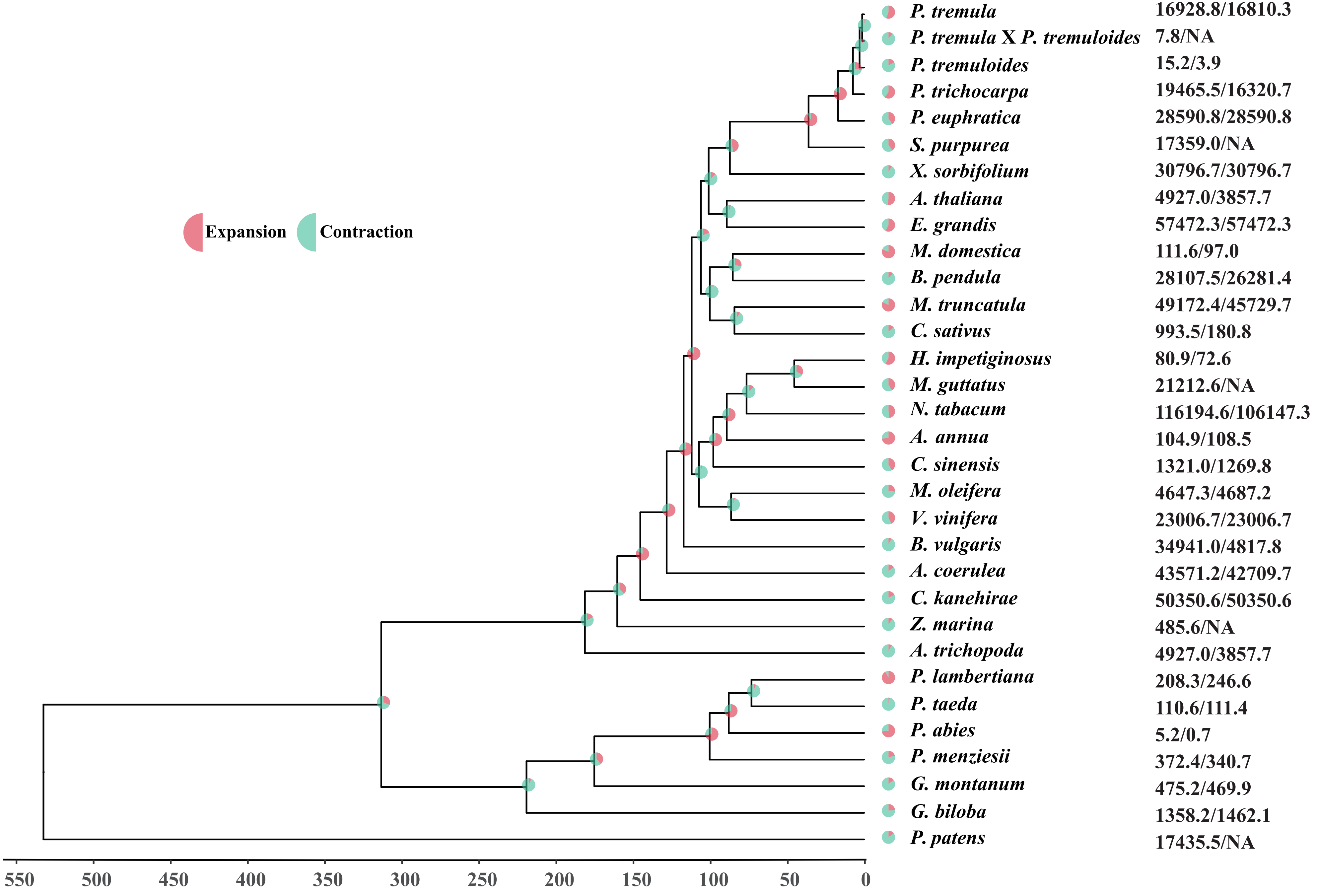

### Supplementary Figure S3 WD40 draft.pdf

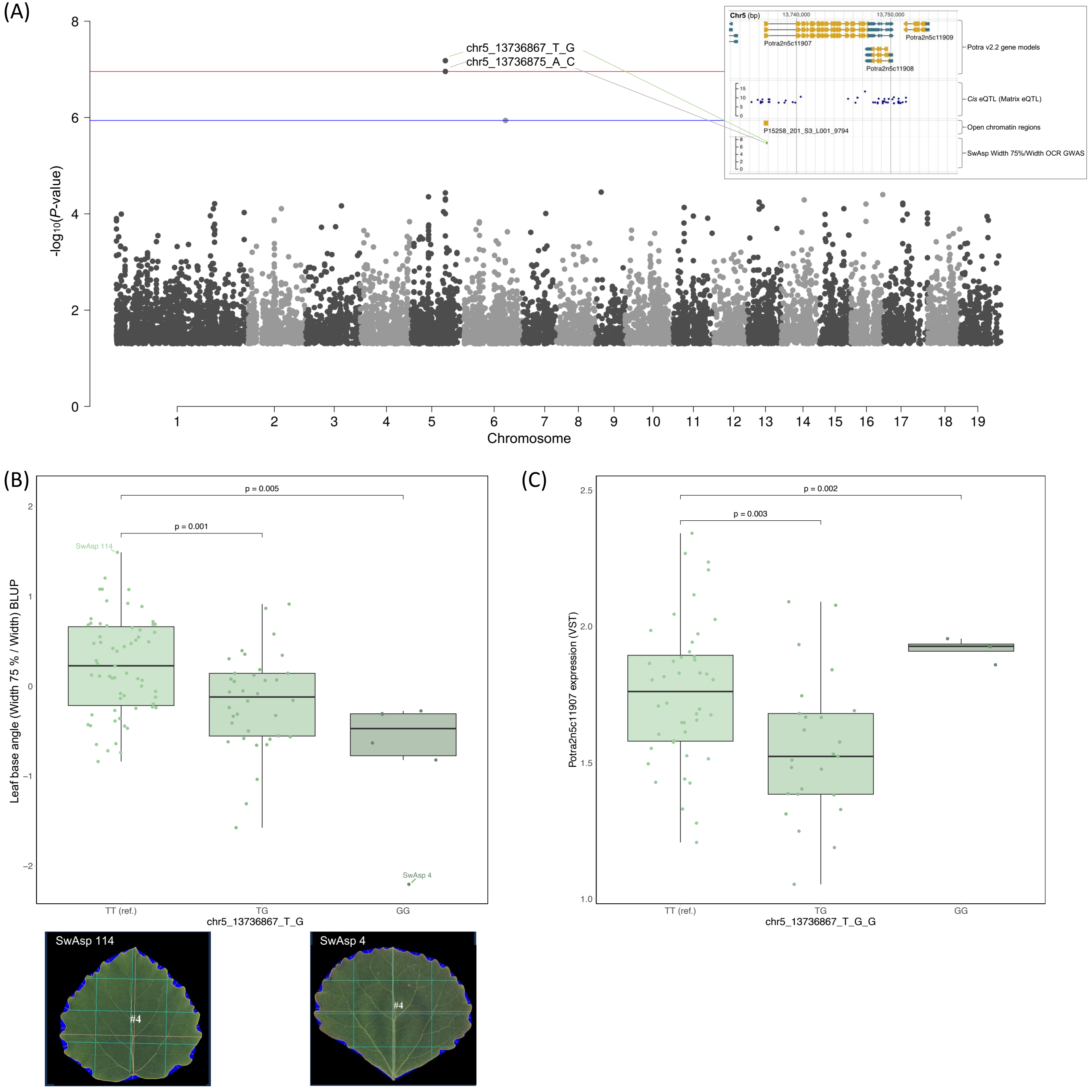

### Supplementary Figure S4. Chromosome 9 JBrowse tracks.pdf

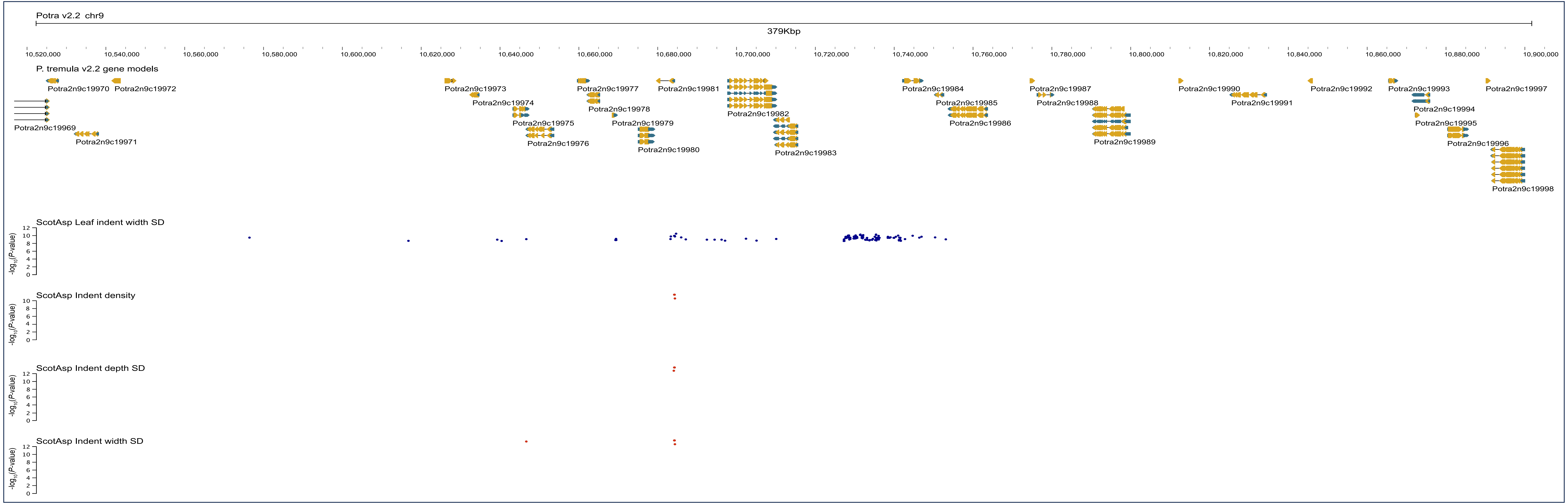
